# Favipiravir, umifenovir and camostat mesylate: a comparative study against SARS-CoV-2

**DOI:** 10.1101/2022.01.11.475889

**Authors:** Mehnmet Altay Unal, Omur Besbinar, Hasan Nazir, Gokce Yagmur Summak, Fatma Bayrakdar, Lucia Gemma Delogu, Tambay Taskin, Sibel Aysil Ozkan, Kamil Can Akcali, Acelya Yilmazer

## Abstract

Since the first cases the coronavirus disease caused by SARS-CoV-2 (COVID-19) reported in December 2019, worldwide continuous efforts have been placed both for the prevention and treatment of this infectious disease. As new variants of the virus emerge, the need for an effective antiviral treatment continues. The concept of preventing SARS-CoV-2 on both pre-entry and post-entry stages has not been much studied. Therefore, we compared the antiviral activities of three antiviral drugs which have been currently used in the clinic. In silico docking analyses and in vitro viral infection in Vero E6 cells were performed to delineate their antiviral effectivity when used alone or in combination. Both in silico and in vitro results suggest that the combinatorial treatment by favipiravir and umifenovir or camostat mesylate has more antiviral activity against SARS-CoV-2 rather than single drug treatment. These results suggest that inhibiting both viral entry and viral replication at the same time is much more effective for the antiviral treatment of SARS-CoV-2.

## Introduction

SARS-CoV-2 has spread all over the world, over 191 countries with more than 181 million confirmed cases and 4 million deaths (Johns Hopkins University, www.coronavirus.jhu.edu/map.html, accessed 29. June. 2021). Despite the ongoing preclinical and clinical studies, unfortunately, there is no other drug approved for SARS-CoV-2 infection rather than FDA approved Remdesivir. Researchers have been focused on drug repurposing to find fast and effective treatment against SARS-CoV-2 infection. Drug repurposing is a technique to search new indication for an approved drug rather than its original indication.

Favipiravir (FPV) (T-705; 6-fluoro-3-hydroxy-2-pyrazinecarboxamide) is a pro-drug which was licensed by the Fujifilm Toyama Chemical Company. FPV prevents viral replication by inhibiting RNA dependent RNA polymerase (RdRP). It has a broad spectrum anti-viral activity in a variety of RNA viruses. It is shown that FPV effectively inhibits the SARS-CoV-2 infection in Vero E6 cells (EC50 = 61.88 μmol·L−1, CC50 > 400 μmol·L−1, SI > 6.46) (Wang et al., 2020a). Cai et al. have compared the effects of favipiravir and Lopinavir/ritonavir against Covid-19. In a non-randomized clinical trial in which 35 patients were treated with Favipiravir and 45 patients were treated with Lopinavir/ritonavir, they found that Favipiravir was associated with faster viral cleansing and higher rates of recovery in chest imaging (Cai et al., 2020).

Umifenovir (Arbidol) is an antiviral drug that is generally used for the treatment of influenza and the major feature is not to have intolerable side effects. Umifenovir interacts with the envelope protein of hepatitis C virus (HCV) leading inhibition of host cell membrane fusion activity (Pécheur et al., 2007). Also umifenovir interacts with viral hemagglutinin (HA) of Influenza virus and inhibits the transition of HA into its functional form (Wang et al., 2020b). S protein of SARS-CoV-2 virus have a structural similarity HA of Influenza virus. Therefore, umifenovir suggested for the antiviral treatment of SARS-CoV-2 virus for the inhibition of viral membrane and host cell membrane fusion (Halboub et al., 2020; Vankadari, 2020; Wang et al., 2020b). Therefore, fusogenic changes in spike protein by Umifenovir in endosomes prevent fusion of viral membrane to the host-cell membrane. Fusogenic mechanisms are common for most of the viruses, therefore umifenovir is being regarded as a wide-spectrum antiviral agent. A randomized, controlled, open-label multicenter trial was conducted among patients with COVID-19. It is shown that umifenovir treatment neither shortened the time of SARS-CoV-2 nor the length of hospitalization of patients with COVID-19 (Lian et al., 2020). Contrary to this finding, there are several reports showing that umifenovir therapy contributes significantly to the clinical and laboratory improvements in COVID-19 patients (Huang et al., 2020) (Nojomi M, 2020). Huang et al. indicated that umifenovir might not only make shorter the viral infection time but also decreased the hospitalization duration of non-severe patients. Therefore they pointed out umifenovir was promising for the clinical outcome of COVID-19 (Huang et al., 2020). In an open-label randomized controlled trial, Nojomi et al., found that use ofumifenovir in hospitalized COVID-19 patients was significantly better against lopinavir/ritonavir combination in terms of clinical and laboratory improvements, including peripheral oxygen saturation, requiring ICU admissions, duration of hospitalization, chest CT involvements, WBC, and ESR (Nojomi M, 2020). Another umifenovir monotherapy trial against lopinavir ritonavir was supportive for this finding as no viral load was detected after 14 days of admission in umifenovir group versus viral load was still present in 44.1% of the patients with lopinavir/ritonavir group (Zhu Z, 2020).

In a Randomized Clinical Trial, monotherapy of 116 patients in Favipiravir and 120 patients in Umifenovir was compared. It was shown that although the comparison of 7 day’s clinical recovery rate of drugs was not significantly different, Favipiravir significantly advanced the latency to cough relief and reduced the duration of pyrexia in moderate COVID-19 patients. It has been mentioned that favipiravir was not related with any differences in ICU admission, AOT/NMV, dyspnea, respiratory failure or all-cause mortality but associated with reduced auxiliary oxygen therapy or noninvasive mechanical ventilation rate with marginal significance (Chen et al., 2020). In another clinical trial patients were treated with Umifenovir-Lopinavir/ritonavir combination and LPV/r only for 5-21 days. Analysis were performed of 16 patients who received Umifenovir-LPV/r in the combination group and 17 who LPV/r only. LPV/r has some gastrointestinal symptoms like elevated levels of bilirubin and it was mentioned that treatment with Umifenovir and LPV/r was well tolerated symptoms in this study (Deng et al., 2020). Prophylactic Umifenovir was shown to have a lower incidence of SARS-CoV-2 infection in health care providers (Yang C, 2020). In this study the cumulative infection rate in health professionals using Umifenovir was found to be significantly lower that of individuals not using Umifenovir. In another retrospective cohort study on family members and health care providers exposed to COVID-19 previously, it was shown thatumifenovir could reduce the risk of infection in hospital and family environments (Zhang JN, 2020).

Camostat mesylate is an oral serine protease inhibitor used for the treatment of acute symptoms related to chronic pancreatitis and postoperative reflux esophagitis, shows promise in cell cultures in combating SARS-CoV-2 through limiting viral entry via inhibiton of TMPRSS2 priming activity with SARS-CoV-2 virus. In a study; patients treated with camostat mesylate showed a decrease in disease severity assessed by the SOFA score. The SOFA score includes sepsis defining parameters, suggesting that camostat mesylate may reduce virus spread from the lung to other organs and/or may dampen the inflammatory response (Hofmann-Winkler et al., 2020). The efficacy and safety of 600 mg QID are currently being evaluated in an phase III study in patients with COVID-19 (Kitagawa et al., 2021). The results from an other double-blind randomized placebo-controlled trial show that among patients hospitalized with Covid-19 camostat mesylate treatment did not significantly improve time to clinical improvement, the risk of intubation or death, time to discontinuation of supplemental oxygen,or any other efficacy outcomes (Gunst et al., 2021).

In this study, we aimed to compare the antiviral activities of these three promising antiviral drugs against SARS-CoV-2. In vitro viral infection and in silico docking analyses were performed to delineate their effectivity when used alone or in combination.

## Material and Methods

### In silico calculations

The chemical structures of umifenovir, camostat mesylate and favipiravir were downloaded from the PubChem (https://pubchem.ncbi.nlm.nih.gov) with the CID numbers 131411, 5284360 and 492405 respectively. Before docking calculations, the structures of chemicals were optimized using B3LYP/6-31G(d) method and basis set of density functional theory (DFT). As a second step, frequency analysis calculations were performed, and stabilization of the optimized structures were checked by the number of image (NImag=0). In addition, the electrostatic potential maps of the optimized structures were determined by using the same method with the basis set 6-311G(d,p). Gaussian 09W and GausView program packages were used for these calculations (Frisch et al. 2009). PDB files of all proteins that model the SARS-CoV-2 virus were downloaded from the PDB databank (https://www.rcsb.org): 6VYB, 6VXX (open and closed forms of spike glycoproteins respectively), 6VWW (Nsp 15 endoribonuclease), 6LU7, 6M03, 6Y84 (main protease), 6VYO (nucleocapsid phosphoprotein), 6M71 (RNA polymerase), 5×29 (envelope protein), 6LXT (fusion protein), 6M0J (ACE2-bounded spike protein) and 1R42 (ACE2 protein). Before using PDB files for docking calculations, first, water and drug molecules were removed from all protein structures using the BIOVIA Discovery Studio 2021 software (Dassault Systèmes 2021) (BIOVIA Discovery Studio - BIOVIA - Dassault Systèmes®), and then the PDB file format of proteins were converted into PDBQT file format using default parameters of AutoDock (ver.1.5.7) software (Trott and Olson, 2009). The binding affinity energies (ΔG, kcal·mol^−1^) between drugs (umifenovir, camostat mesylate, favipiravir) and proteins were calculated, with single and multiple drug simulation docking (Li et al., 2012) approaches by blind docking (supplementary material, RawData.zip) using AutoDock 4.2.6 program. As a result of these calculations, the top 10 conformations with the highest affinity energies were taken into consideration. BIOVIA and UCSF Chimera ver. 1.15 (Pettersen et al., 2004) softwares were used for two- and three-dimensional analysis and imaging of complexes. Furthermore, to detect the cavity and druggability of cavities of prepared protein models, CAVITY software (http://repharma.pku.edu.cn) was used (Yuan et al., 2013). In order to predict ADME-T (absorption, distribution, metabolism, excretion and toxicity) profiles of umifenovir, camostat mesylate and favipiravir, OSIRIS Property Explorer (https://www.organic-chemistry.org/prog/peo) and SwissADME online web server (http://www.swissadme.ch) were used.

### Cell Culture

African green monkey kidney Vero E6 cell line was purchased from ATCC and maintained in DMEM media containing 10% FBS and 1% antibiotics. Local SARS-CoV-2 isolates (hCoV-19/Turkey/HSGM-302/2020 (Clade GR) with GISAID accession number of EPI_ISL_437313|2020-03-27 was used in this study. Viruses were propagated in Vero E6 cells by using DMEM media containing 2% FBS and 1% antibiotics. All virus-related experiments were performed at Biosafety Level 3 laboratories.

### Viral Infection

Vero E6 cells were seeded onto 96-well plates at confluency. Cells were first incubated with drugs at different concentrations and later infected with SARS-CoV-2 (MOI 0.1) After the final treatment, plates were incubated at 5% CO_2_ at 37ºC. Plates were monitored for cytopathic activity for 5 days.

### qRT-PCR analysis

At the end of day 5, cell culture supernatants were collected. Total RNA was isolated via MPLC Total Nucleic Acid Isolation Kit (Roche) using automated MagNA Pure LC Instrument (Roche). One-step qRT-PCR was performed using the Transcriptor One-Step RT-PCR Kit (Roche) by using 5μL of samples per each 20 μL reaction volume. In order to calculate viral copies per μL, a standard curve was constructed by using 5 different standards of known copy numbers according to the viral N gene.

### Statistical Analysis

All values are expressed as mean ± ST.D. Comparison between groups was performed by one-way ANOVA, followed by a Tukey’s post hoc multiple comparisons by the statistics program GraphPad. At least three independent replicated were analyzed.

## Results

First, we evaluated the in vitro antiviral activity of these drug molecules based on the docking results. First, Vero E6 cells were infected following exposure with drug molecules. Cells were monitored for 5 days, and at the end of the experiment, cell culture supernatants were collected and used to quantify viral copy numbers via qRT-PCR. According to Figures 2A and 2B, umifenovir and favipiravir showed a similar antiviral activity profile, where more than 80% of viral inhibition was observed at 650 μM drug concentration. On the other hand, camostate mesylate treatment was only able to achieve the same inhibition levels at a much higher dose (Figure 2C). Later, when drugs were used in combination, it can be seen that treating Vero E6 cells with umifenovir and favipiravir at the same time, SARS-CoV-2 copy numbers were significantly reduced compared to the cells treated with either of the drugs at each dilution (Figure 3A. Furthermore, combining camostat mesylate with umifenovir or favipiravir also significantly inhibited viral infection compared to the cells treated only with camostate mesylate (Figure 3B and 3C). Considering that umifenovir and camostat mesylate are both acting at the entry of viral particles into host cells, combining these drugs with a post-entry drug such as favipiravir gives the most promising antiviral activity in Vero E6 cells. Therefore, in parallel to the docking analysis, these results further support the importance of using drug combinations to achieve better antiviral activity.

**Figure 1.**
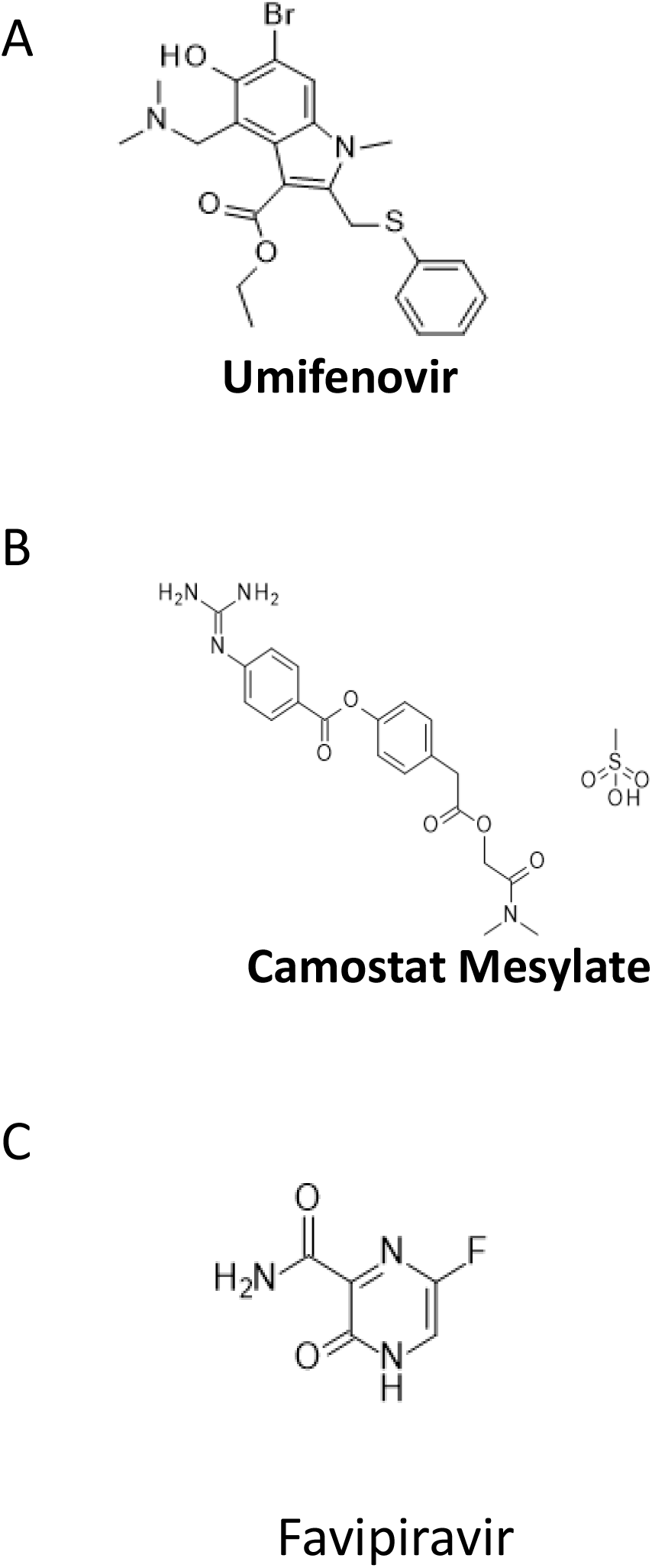
The structure of antiviral drug molecules used in this study. A) Umifenovir, B) Camostat mesylate, C) Umifenovir.

**Figure 2.**
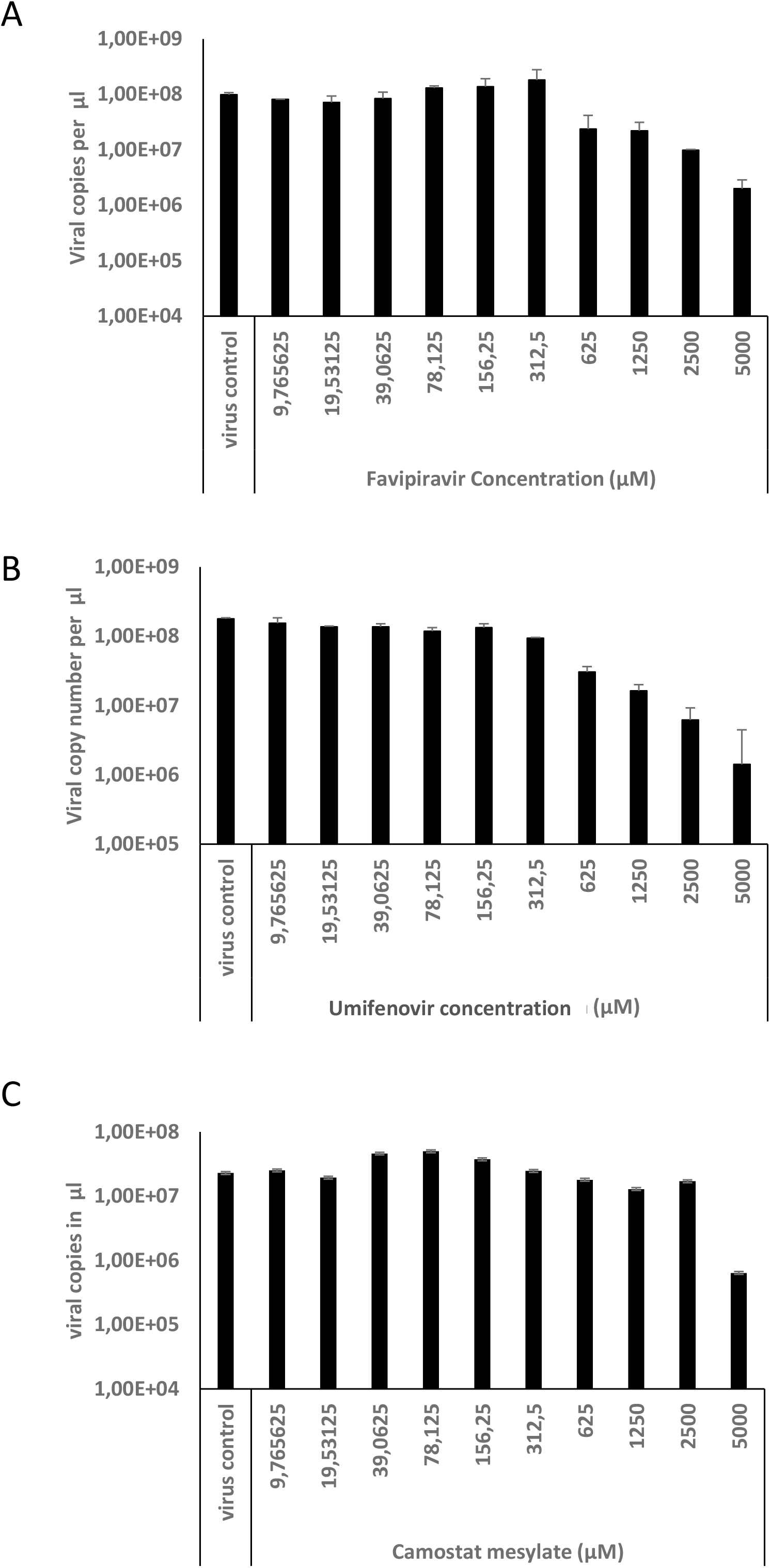
In vitro antiviral activity of drug molecules when used alone. Vero E6 cells were pre-treated with antiviral drugs at different concentrations and after 1 hour, viral particles were added into the culture media. At the end of day 5, cell culture supernatants were collected and number of viral particles were quantified by qRT-PCR.

**Figure 3.**
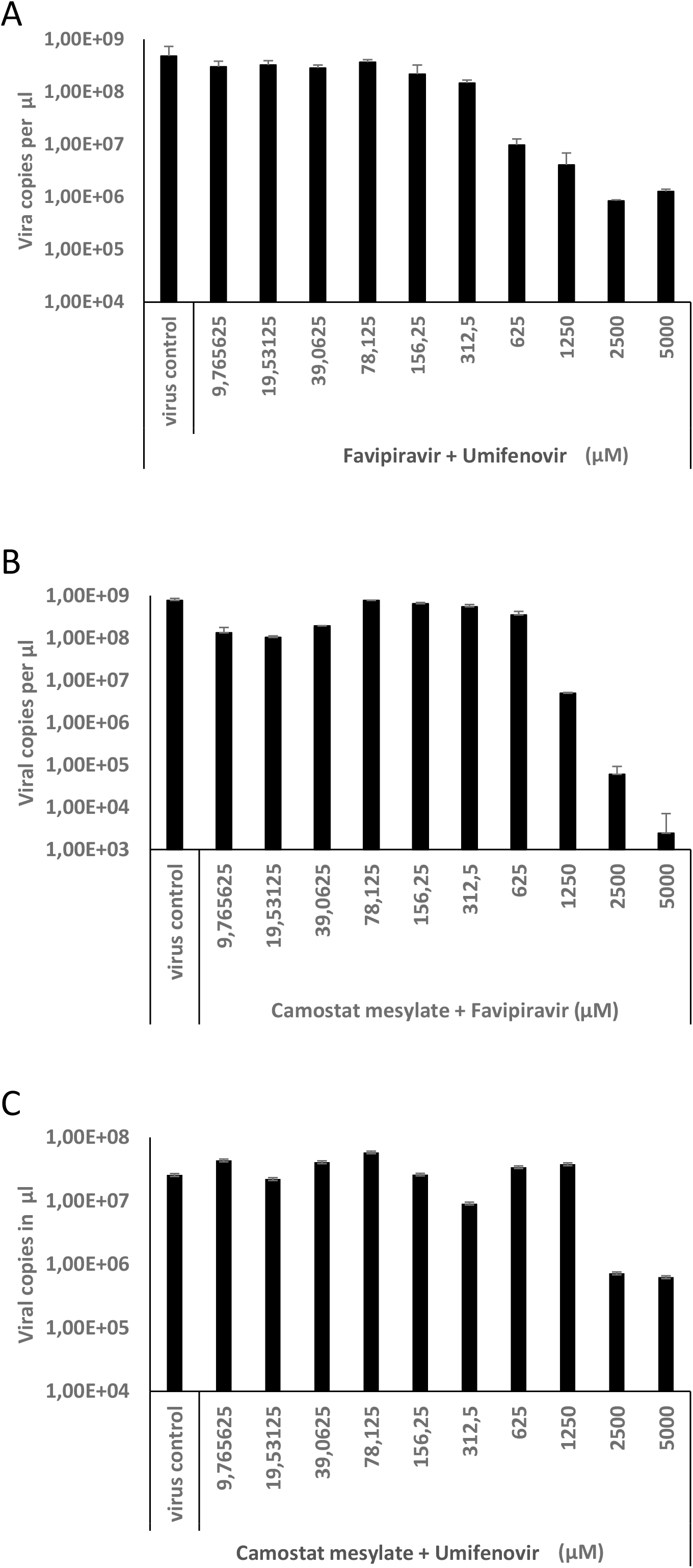
In vitro antiviral activity of drug molecules when used in combination. Vero E6 cells were pre-treated with antiviral drug combinations at different concentrations and after 1 hour, viral particles were added into the culture media. At the end of day 5, cell culture supernatants were collected and number of viral particles were quantified by qRT-PCR.

After performing in vitro experiments, molecular docking analtysis was performed to further support and delineate the antiviral activities observed above. ESP maps of umifenovir, camostat mesylate (CamostatM) and favipiravir molecules show nucleophilic and electrophilic attack regions (Figure S1). These characteristics indicates that the drug molecules are capable of being both hydrogen donor and acceptor, conduct non-bonding interaction with amino acids, and also held dynamically on the protein surface. In addition, ADME-T calculations suggest that these drug active chemicals are not mutagenic and other feature values are also in acceptable range (Figure S2). Calculated the highest single affinity docking scores of umifenovir, CamostatM, favipiravir and as duo umifenovir+camostatM, camostatM+favipiravir, umifenovir+favipiravir protein complexes were given in Table S1 (and supplementary materials, RawData.zip). If Table S1 results were analyzed, in the single effect, the highest affinity value with -8.35 kcal/mol against 6LU7 (main protease) and the lowest affinity value with -2.97 kcal/mol against 5×29 (envelope) was umifenovir and favipiravir respectively. As for duo effect, shows that the highest affinity values with -8.96 + -4.36 kcal/mol against 6VXX (closed form of spike) and the lowest affinity values with -3.20 + - 3.52 kcal/mol against 5×29 (envelope) was camostatM+favipiravir and favipiravir+umifenovir, each to each. If the data in Table S1 are analyzed in terms of the average effect of umifenovir, camostatM and favipiravir on proteins, it can be said that, the single and duo effect changes respectively as camostatM>umifenovir>favipiravir and camostatM+favipiravir> umifenovir+camostatM>favipiravir+umifenovir. However, when dual effects examined, it was clearly seen that especially camostatM+favipiravir increases the effectiveness. This effect was calculated to be highest against Spike proteins (6M0J, -8.26 + -5.20, 6VXX, -8.96 + -4.36 and 6VYB, -8.63 + -4.85). As a matter of fact, this calculated increase in efficiency is also consistent with the experimental results (Figure 3B). All three-dimensional shots of 10 highest energy conformations of single and duo combinations of umifenovir, camostatM and favipiravir on proteins are given in RawData.zip (supplementary material). And also, three-dimensional shots showing the protein affinity of the camostatM+favipiravir combination, in which the highest efficiency was observed, are shown in Figure S3. The effect of camostatM+favipiravir combination on 6VYB, 6VXX spike proteins and 6M0J spike ACE2 bounded protein, increasing their affinity energies compare to their single state (Table S1), is especially worth examining. In Figure S4, the effects of single camostatM, favipiravir and duo camostatM+favipiravir on 6VYB, 6VXX and 6M0J proteins are shown comparatively. In the duo combination of camostatM + favipiravir, it can be said that camostatM and favipiravir both protect their single affinity sites and also distributed uniformly on the protein surfaces, thus the ligands affect each other in the direction of expanding the affinity area. According to UniProtKB database (https://www.uniprot.org/uniprot/P0DTC2), 6VYB (open form) and 6VXX (close form) spike proteins have two binding areas. These are amino acids between 319-541 (Receptor Binding Domain, RBD) and 437-508 (receptor-binding motif). Besides, 30-41, 82-84 and 353-357 amino acids are active in binding of ACE2 protein (1R42) to spike proteins (https://www.uniprot.org/uniprot/Q9BYF1). If representations in Figure S4 are examined, it is seen that the combination of comastatM + favipiravir shows affinity for these active sites of the proteins. Indeed, the two- and three-dimensional representations of ligand-amino acid interactions which were given in Figure S5A-C confirm this situation. Considering its affinity for the 6M0J spike-ACE2 bounded protein (Figure S5C), it is seen that camostatM+favipiravir have affinity for both parts and interface of the protein, and to interact with amino acids in hydrogen, van der Waals, and non-covalent bonds: (6M0J; A(1R42), LSY:31, ASN:33, HIS:34, GLU:35, GLU:37, ASP:38, LEU:39, TRY:83, PRO:84, ARG:357; E(Spike), TRP:436, ASN:437, SER:438, ASN:439, ASN:440, LEU:441, LEU:452, TYR:453, GLU:484, TYR:489, PHE:490, LEU:492, GLN:493, SER:494, TYR:495, VAL:503, TYR:505, GLN:506, TYR:508, ARG:509). It can also be said that ligands interact with amino acids in areas where solvent accessibility is low (Figure S5, SAS shots, green area). On the other hand, the duo combinations of umifenovir+camostatM and favipiravir+umifenovir are also in agreement with the experimental data (Figure 3C and 3A). Figure S6A and S6B show the effect of camostatM+umifenovir and favipiravir+umifenovir on 6VYB, 6VXX and 6M0J proteins, respectively. In addition, 2D maps showing the effects of duo combinations at the 6M0J protein interface are also given. As seen from the 2D maps, dual drug combinations have effects on Spike-binding amino acids of ACE2 and the RBD regions of Spike proteins (open and closed forms), although their affinity energies are lower than CamostatM+favipiravir (TableS1).

## Discussion

SARS-CoV-2 virus classified as positive sense single stranded RNA (+ssRNA) virus. +ssRNA viral genome is enveloped with nucleocapsid (N) protein. The viral membrane of SARS-CoV-2 composed of envelope (E), membrane (M), and spike (S) proteins (Murgolo et al., 2021). S1 and S2 domains containing S protein mediates host cell entry via interacting with angiotensin converting enzyme 2 (ACE2) receptor located on host cell membrane. S1 domain of S protein mediates the interaction of ACE2 and S protein by containing receptor-binding domain (RBD) (Lan et al., 2020; Letko et al., 2020). ACE2 interacting S protein is primed by transmembrane protease serine 2 (TMPRSS2) protein to reveal active S2 domain of S protein. S2 protein mediates fusion of viral membrane with host cell membrane leading disclosure of +ssRNA in the host cell cytoplasm (Hoffmann et al., 2020a, 2020b). Viral translation begins with the disclosure of +ssRNA after viral cell entry leading t the translation of Open Reading Frames, ORF1a and ORF1b. Translation of ORF1a and ORF1b lead production of pp1a and pp1ab polyproteins respectively (Finkel et al., 2021). These polyproteins later produce non-structural proteins (nsps) by proteolytic cleaveage. Proteolytic cleaveage of pp1a and pp1ab leads to production of nsp1-10 and nsp12-16 proteins, respectively. The combination of nsp3, nsp5 and nsp8 leads to the expression of viral helicase protein whereas nsp12 leads to the expression of RNA-dependent RNA polymerase (RdRp) (Hillen et al., 2020). Host endomembranes diverted into replication organelles – mostly ER-derived double membrane vesicles (DMVs)-with the help of nsp3, nsp4 and nsp6 (Knoops et al., 2008). RdRp and replicase protein placed in DMVs which leading protective microenvironment for viral transcription and replication. For the formation of new viral particles, translated proteins in DMVs translocate into ER membrane, and then transit into ER-to-golgi intermediate compartment (ERGIC). The replicated genomic material encapsulated with the produced E, M, S and N proteins. Finally, the assembled virus is released by exocytosis through golgi apparatus from host cell (V’kovski et al., 2020).

As summarized above, the viral infection cycle is complex and dependent on various different viral proteins. In this study, we aimed to explore if the antiviral drugs that have been used in the clinic against SARS-CoV-2, can be used in combination to improve the antiviral activity (Figure 4). Umifenovir has been suggested for the antiviral treatment of SARS-CoV-2 for the inhibition viral entry (Halboub et al., 2020; Vankadari, 2020; Wang et al., 2020b). Camostat mesylate is a serine protease inhibitor which inhibits the catalytic activity of TMPRSS2 protein (Hoffmann et al., 2021). It has been suggested that inhibition of TMPRSS2 activity results reduction of SARS-CoV-2 viral entry by blocking exposure of S2 domain of S protein via TMPRSS2 priming (Uno, 2020; Breining et al., 2021), which refers to the combinatorial umifenovir-camostat mesylate treatment in our study. The possibility that camostat mesylate or other TMPRSS2 inhibitors administered in higher doses or combinations with other drugs might be effective in lowering the risk of disease progression. Hypothetically, combination of drugs which have different mechanism of action can eventuate to an effective antiviral thearpy for SARS-CoV-2 infection. Preclincal and clinical studies are ongoing for drug combinations to combat SARS-CoV-2.

**Figure 4.**
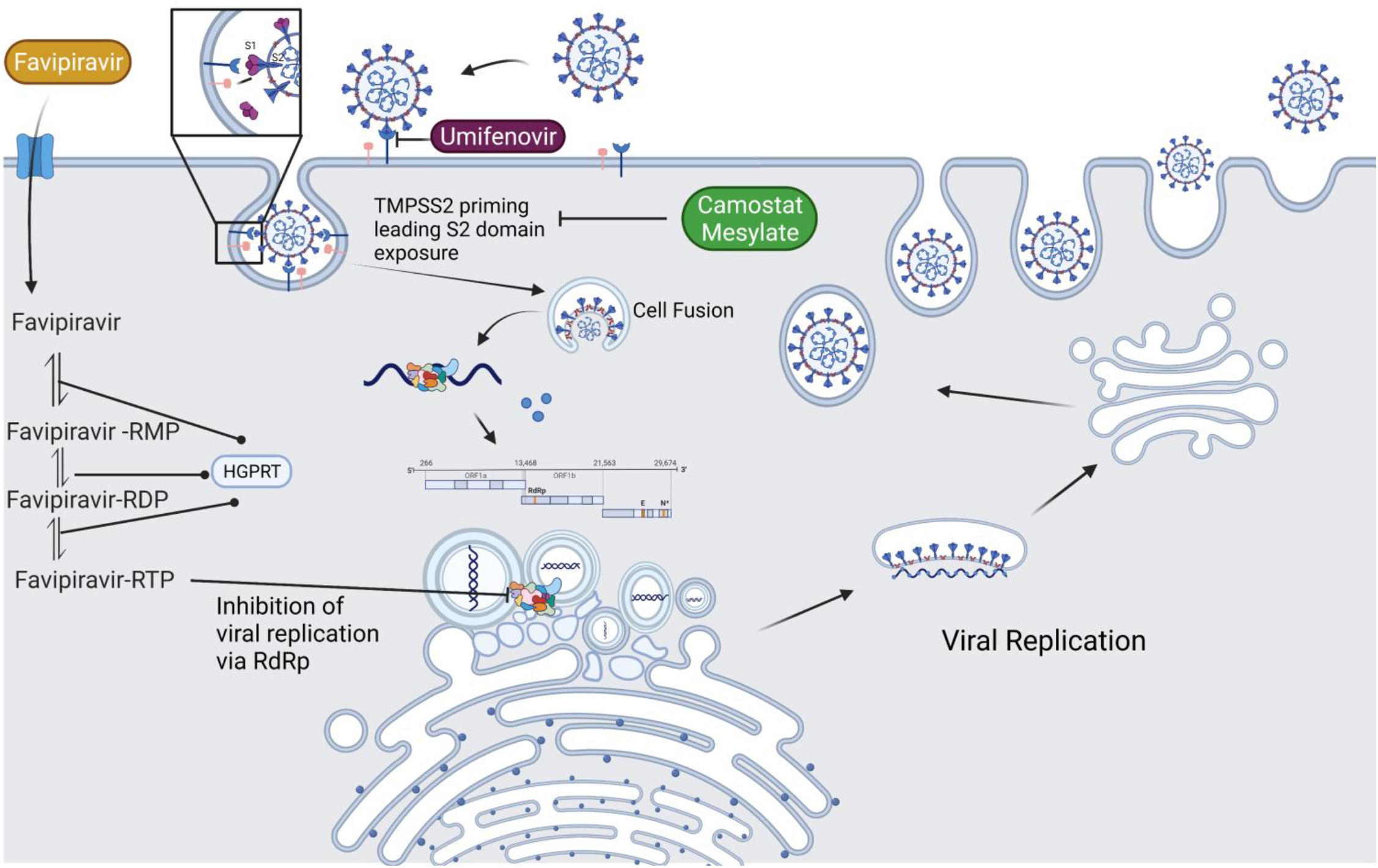
Mechanisms of antiviral drugs. SARS-CoV-2 virus is classified as +ssRNA virus. S protein of SARS-CoV-2 mediates host cell entry via interacting with ACE2 receptor located on host cell membrane. ACE2 interacting S protein is primed by TMPRSS2 protein to reveal active S2 domain of S protein. S2 protein mediates fusion of viral membrane with host cell membrane leading exposure of +ssRNA in the host cell cytoplasm which results viral replication. Umifenovir inhibits the interaction of S protein and ACE2 receptor. Camostat Mesylate inthibits the proteolytic activity of TMPRSS2. Translation leads to production of nsps. Host endomembranes diverted into replication organelles. RdRp and replicase protein placed in DMVs for tranlation of viral proteins. Translated viral proteins translocate into ER membrane and then transit to ERGIC. Viral genomic material wrapped with E, M, S, and N viral proteins and newly produced virus relased by exocytosis through Golgi apparatus. Favipiravir enters the cell and phoshorylated with HGPRT. The Favipiravir-RTP form of Favipiravir competes with purine bases and inhibits the replicative activity of RdRp.

The concept of preventing the virus on both pre-entry and post-entry stages has not been previously studied. Favipiravir is broad-spectrum antiviral pro-dug which inhibits viral replication by influencing the activiy of RdRp (Delang et al., 2018). Favipiravir enters cell through cell membrane and phophoribosylated by Hypoxanthine Guanine Phosphoribosyltransferase (HGPRT) to become Favipiravir-ribose-5’-monophosphate (Favipiravir-RMP) (Naesens et al., 2013). Favipiravir-RMP turns into Favipiravir-ribose-5-diphosphate (Favipiravir-RDP) and Favipiravir-ribose-5’-triphosphate (Favipiravir-RTP) respectively by phosphorylation. Favipiravir-RTP competes with purine bases – predominantly GTP- to influence viral RdRp mediated viral replication (Furuta et al., 2013). Favipiravir-RTP can base pair with both cysteine and uracil base pairs. In our study, the antiviral activity of favipiravir and umifenovir or favipiravir and camostat mesylate combinations are much more effective than mono drug threapies. Doi et al. showed that combination of favipiravir and another TMPRSS2 inhibitor nafamostat mesylate showed promising results indicating combinatorial treatment inhibits both viral replication and viral entry (Doi et al., 2020). Therefore, it is reasonable to target both viral entry and viral replication at the same time to eliminate the SARS-CoV-2 infection. In summary, the combinatiorial treatment via favipiravir and umifenovir or camostat mesylate has more antiviral activity against SARS-CoV-2 rather than single drug treatment by inhibiting both viral entry and viral replication.

## Supporting information

Supplemental Raw Data

## Acknowledgement

The research for this paper was financially supported by the Scientific and Technological Research Council of Turkey (TUBITAK) under grant number 18AG020. We thank the Ankara University Information Technologies Department for technical support.

## References

Liu, F., Marquardt, S., Lister, C., Swiezewski, S. & Dean, C. Targeted 3’ processing of antisense transcripts triggers Arabidopsis FLC chromatin silencing. Science 327, 94–97 (2010).

Rhinn, H. et al. Alternative α-synuclein transcript usage as a convergent mechanism in Parkinson’s disease pathology. Nat. Commun. 3, 1084 (2012).

Solana, J. et al. Conserved functional antagonism of CELF and MBNL proteins controls stem cell-specific alternative splicing in planarians. Elife 5, (2016).

Mudge, J. M. & Harrow, J. The state of play in higher eukaryote gene annotation. Nat. Rev. Genet. 17, 758–772 (2016).

Frankish, A. et al. GENCODE reference annotation for the human and mouse genomes. Nucleic Acids Res. 47, D766–D773 (2019).

McGarvey, K. M. et al. Mouse genome annotation by the RefSeq project. Mamm. Genome 26, 379–390 (2015).

Berardini, T. Z. et al. The Arabidopsis information resource: Making and mining the ‘gold standard’ annotated reference plant genome. Genesis 53, 474–485 (2015).

FANTOM Consortium and the RIKEN PMI and CLST (DGT) et al. A promoter-level mammalian expression atlas. Nature 507, 462–470 (2014).

Wu, P.-Y., Phan, J. H. & Wang, M. D. Assessing the impact of human genome annotation choice on RNA-seq expression estimates. BMC Bioinformatics 14 Suppl 11, S8 (2013).

SEQC/MAQC-III Consortium. A comprehensive assessment of RNA-seq accuracy, reproducibility and information content by the Sequencing Quality Control Consortium. Nat. Biotechnol. 32, 903–914 (2014).

Guigó, R. et al. EGASP: the human ENCODE Genome Annotation Assessment Project. Genome Biol. 7 Suppl 1, S2.1–31 (2006).

Stark, R., Grzelak, M. & Hadfield, J. RNA sequencing: the teenage years. Nat. Rev. Genet. 20, 631–656 (2019).

Levin, J. Z. et al. Comprehensive comparative analysis of strand-specific RNA sequencing methods. Nat. Methods 7, 709–715 (2010).

Murata, M. et al. Detecting expressed genes using CAGE. Methods Mol. Biol. 1164, 67–85 (2014).

Adiconis, X. et al. Comprehensive comparative analysis of 5’-end RNA-sequencing methods. Nat. Methods 15, 505–511 (2018).

Schon, M. A., Kellner, M. J. & Plotnikova, A. NanoPARE: parallel analysis of RNA 5′ ends from low-input RNA. Genome Res. (2018).

Cvetesic, N. et al. SLIC-CAGE: high-resolution transcription start site mapping using nanogram-levels of total RNA. Genome Res. 28, 1943–1956 (2018).

Jan, C. H., Friedman, R. C., Ruby, J. G. & Bartel, D. P. Formation, regulation and evolution of Caenorhabditis elegans 3’UTRs. Nature 469, 97–101 (2011).

Moll, P., Ante, M., Seitz, A. & Reda, T. QuantSeq 3′ mRNA sequencing for RNA quantification. Nat. Methods 11, i–iii (2014).

Pelechano, V., Wei, W. & Steinmetz, L. M. Extensive transcriptional heterogeneity revealed by isoform profiling. Nature 497, 127–131 (2013).

Wang, J. et al. TIF-Seq2 disentangles overlapping isoforms in complex human transcriptomes. Nucleic Acids Res. 48, e104 (2020).

Picelli, S. et al. Smart-seq2 for sensitive full-length transcriptome profiling in single cells. Nat. Methods 10, 1096–1098 (2013).

Hagemann-Jensen, M. et al. Single-cell RNA counting at allele and isoform resolution using Smart-seq3. Nat. Biotechnol. 38, 708–714 (2020).

Cao, J. et al. The single-cell transcriptional landscape of mammalian organogenesis. Nature 566, 496–502 (2019).

Cao, J. et al. Comprehensive single-cell transcriptional profiling of a multicellular organism. Science 357, 661–667 (2017).

Zheng, G. X. Y. et al. Massively parallel digital transcriptional profiling of single cells. Nat. Commun. 8, 14049 (2017).

Garalde, D. R. et al. Highly parallel direct RNA sequencing on an array of nanopores. Nat. Methods 15, 201–206 (2018).

Wan, Y. et al. Systematic identification of intergenic long-noncoding RNAs in mouse retinas using full-length isoform sequencing. BMC Genomics 20, 559 (2019).

Steijger, T. et al. Assessment of transcript reconstruction methods for RNA-seq. Nat. Methods 10, 1177–1184 (2013).

Tardaguila, M. et al. SQANTI: extensive characterization of long-read transcript sequences for quality control in full-length transcriptome identification and quantification. Genome Res. (2018) doi:10.1101/gr.222976.117.

Kuo, R. I. et al. Normalized long read RNA sequencing in chicken reveals transcriptome complexity similar to human. BMC Genomics 18, 323 (2017).

Tang, A. D. et al. Full-length transcript characterization of SF3B1 mutation in chronic lymphocytic leukemia reveals downregulation of retained introns. Nat. Commun. 11, 1438 (2020).

Kovaka, S. et al. Transcriptome assembly from long-read RNA-seq alignments with StringTie2. Genome Biol. 20, 278 (2019).

Cumbie, J. S., Ivanchenko, M. G. & Megraw, M. NanoCAGE-XL and CapFilter: an approach to genome wide identification of high confidence transcription start sites. BMC Genomics 16, 597 (2015).

de Rie, D. et al. An integrated expression atlas of miRNAs and their promoters in human and mouse. Nat. Biotechnol. 35, 872–878 (2017).

Thieffry, A. et al. Characterization of Arabidopsis thaliana Promoter Bidirectionality and Antisense RNAs by Inactivation of Nuclear RNA Decay Pathways. Plant Cell 32, 1845–1867 (2020).

Sherstnev, A. et al. Direct sequencing of Arabidopsis thaliana RNA reveals patterns of cleavage and polyadenylation. Nat. Struct. Mol. Biol. 19, 845–852 (2012).

Pertea, M. et al. StringTie enables improved reconstruction of a transcriptome from RNA-seq reads. Nat. Biotechnol. 33, 290–295 (2015).

Shao, M. & Kingsford, C. Accurate assembly of transcripts through phase-preserving graph decomposition. Nat. Biotechnol. 35, 1167–1169 (2017).

Trapnell, C. et al. Differential gene and transcript expression analysis of RNA-seq experiments with TopHat and Cufflinks. Nat. Protoc. 7, 562–578 (2012).

Amarasinghe, S. L. et al. Opportunities and challenges in long-read sequencing data analysis. Genome Biol. 21, 30 (2020).

Goodwin, S., McPherson, J. D. & McCombie, W. R. Coming of age: ten years of next-generation sequencing technologies. Nat. Rev. Genet. 17, 333–351 (2016).

Balázs, Z. et al. Template-switching artifacts resemble alternative polyadenylation. BMC Genomics 20, 824 (2019).

Tang, D. T. P. et al. Suppression of artifacts and barcode bias in high-throughput transcriptome analyses utilizing template switching. Nucleic Acids Res. 41, e44 (2013).

Gordon, S. P. et al. Widespread Polycistronic Transcripts in Fungi Revealed by Single-Molecule mRNA Sequencing. PLoS One 10, e0132628 (2015).

Xie, Z., Kasschau, K. D. & Carrington, J. C. Negative Feedback Regulation of Dicer-Like1 in Arabidopsis by microRNA-Guided mRNA Degradation. Current Biology vol. 13 784–789 (2003).

Rajagopalan, R., Vaucheret, H., Trejo, J. & Bartel, D. P. A diverse and evolutionarily fluid set of microRNAs in Arabidopsis thaliana. Genes Dev. 20, 3407–3425 (2006).

Westoby, J., Artemov, P., Hemberg, M. & Ferguson-Smith, A. Obstacles to detecting isoforms using full-length scRNA-seq data. Genome Biol. 21, 74 (2020).

Natarajan, K. N. et al. Comparative analysis of sequencing technologies for single-cell transcriptomics. Genome Biol. 20, 70 (2019).

Paul, L. et al. SIRVs: Spike-In RNA Variants as External Isoform Controls in RNA-Sequencing. bioRxiv 080747 (2016) doi:10.1101/080747.

Nam, J.-W. et al. Global analyses of the effect of different cellular contexts on microRNA targeting. Mol. Cell 53, 1031–1043 (2014).

Lagarde, J. et al. High-throughput annotation of full-length long noncoding RNAs with capture long-read sequencing. Nat. Genet. 49, 1731–1740 (2017).

Niknafs, Y. S., Pandian, B., Iyer, H. K., Chinnaiyan, A. M. & Iyer, M. K. TACO produces robust multisample transcriptome assemblies from RNA-seq. Nat. Methods 14, 68–70 (2017).

Song, L., Sabunciyan, S., Yang, G. & Florea, L. A multi-sample approach increases the accuracy of transcript assembly. Nat. Commun. 10, 5000 (2019).

Pertea, G. & Pertea, M. GFF Utilities: GffRead and GffCompare. F1000Res. 9, 304 (2020).

Wang, B. et al. Unveiling the complexity of the maize transcriptome by single-molecule long-read sequencing. Nat. Commun. 7, 11708 (2016).

Gupta, I. et al. Single-cell isoform RNA sequencing characterizes isoforms in thousands of cerebellar cells. Nat. Biotechnol. (2018) doi:10.1038/nbt.4259.

Philpott, M. et al. Nanopore sequencing of single-cell transcriptomes with scCOLOR-seq. Nat. Biotechnol. (2021) doi:10.1038/s41587-021-00965-w.

Zheng, Y. F., Chen, Z. C., Shi, Z. X., Hu, K. H. & Zhong, J. Y. HIT-scISOseq: High-throughput and high-accuracy single-cell full-length isoform sequencing for corneal epithelium. bioRxiv (2020).

Tabula Muris Consortium et al. Single-cell transcriptomics of 20 mouse organs creates a Tabula Muris. Nature 562, 367–372 (2018).

Quake, S. R. & Sapiens Consortium, T. The Tabula Sapiens: a single cell transcriptomic atlas of multiple organs from individual human donors. bioRxiv (2021).

Martin, M. Cutadapt removes adapter sequences from high-throughput sequencing reads. EMBnet.journal 17, 10–12 (2011).

Dobin, A. et al. STAR: ultrafast universal RNA-seq aligner. Bioinformatics 29, 15–21 (2013).

BIOVIA Discovery Studio - BIOVIA - Dassault Systèmes® Available at: https://www.3ds.com/products-services/biovia/products/molecular-modeling-simulation/biovia-discovery-studio/ [Accessed May 9, 2021].

Breining, P., Frølund, A. L., Højen, J. F., Gunst, J. D., Staerke, N. B., Saedder, E., et al. (2021). Camostat mesylate against SARS-CoV-2 and COVID-19—Rationale, dosing and safety. Basic \& Clin. Pharmacol. \& Toxicol. 128, 204–212. doi:https://doi.org/10.1111/bcpt.13533.

Cai, Q., Yang, M., Liu, D., Chen, J., Shu, D., Xia, J., et al. (2020). Experimental Treatment with Favipiravir for COVID-19: An Open-Label Control Study. Eng. (Beijing, China) 6, 1192–1198. doi:10.1016/j.eng.2020.03.007.

Chen, C., Zhang, Y., Huang, J., Yin, P., Cheng, Z., Wu, J., et al. (2020). Favipiravir versus Arbidol for COVID-19: A Randomized Clinical Trial. medRxiv, 2020.03.17.20037432. doi:10.1101/2020.03.17.20037432.

Delang, L., Abdelnabi, R., and Neyts, J. (2018). Favipiravir as a potential countermeasure against neglected and emerging RNA viruses. Antiviral Res. 153, 85–94. doi:10.1016/j.antiviral.2018.03.003.

Deng, L., Li, C., Zeng, Q., Liu, X., Li, X., Zhang, H., et al. (2020). Arbidol combined with LPV/r versus LPV/r alone against Corona Virus Disease 2019: A retrospective cohort study. J. Infect. 81, e1–e5. doi:10.1016/j.jinf.2020.03.002.

Doi, K., Ikeda, M., Hayase, N., Moriya, K., Morimura, N., Maehara, H., et al. (2020). Nafamostat mesylate treatment in combination with favipiravir for patients critically ill with Covid-19: a case series. Crit. Care 24, 1–4. doi:10.1186/s13054-020-03078-z.

Finkel, Y., Mizrahi, O., Nachshon, A., Weingarten-Gabbay, S., Morgenstern, D., Yahalom-Ronen, Y., et al. (2021). The coding capacity of SARS-CoV-2. Nature 589, 125–130. doi:10.1038/s41586-020-2739-1.

Frisch, M.J., Trucks, G.W., Schlegel, H.B., Scuseria, G.E., Robb, M.A., Cheeseman, J.R., Scalmani, G., Barone, V., Mennucci, B., Petersson, G.A., et al. 2009. Gaussian G09. Gaussian Inc., Wallingford, CT, USA. Available from https://gaussian.com/glossary/g09/.

Furuta, Y., Gowen, B. B., Takahashi, K., Shiraki, K., Smee, D. F., and Barnard, D. L. (2013). Favipiravir (T-705), a novel viral RNA polymerase inhibitor. Antiviral Res. 100, 446–454. doi:10.1016/j.antiviral.2013.09.015.

Gunst, J. D., Staerke, N. B., Pahus, M. H., Kristensen, L. H., Bodilsen, J., Lohse, N., et al. (2021). Efficacy of the TMPRSS2 inhibitor camostat mesilate in patients hospitalized with Covid-19-a double-blind randomized controlled trial. EClinicalMedicine 35, 100849. doi:https://doi.org/10.1016/j.eclinm.2021.100849.

Halboub, E., AL-Maweri;, S. A., and Al-Soneidar, W. A. (2020). COVID-19: A review of the proposed pharmacological treatments. Eur. J. Pharmacol., 1–5.

Hillen, H. S., Kokic, G., Farnung, L., Dienemann, C., Tegunov, D., and Cramer, P. (2020). Structure of replicating SARS-CoV-2 polymerase. Nature 584, 154–156. doi:10.1038/s41586-020-2368-8.

Hoffmann, M., Hofmann-Winkler, H., Smith, J. C., Krüger, N., Arora, P., Sørensen, L. K., et al. (2021). Camostat mesylate inhibits SARS-CoV-2 activation by TMPRSS2-related proteases and its metabolite GBPA exerts antiviral activity. EBioMedicine 65. doi:10.1016/j.ebiom.2021.103255.

Hoffmann, M., Kleine-Weber, H., and Pöhlmann, S. (2020a). A Multibasic Cleavage Site in the Spike Protein of SARS-CoV-2 Is Essential for Infection of Human Lung Cells. Mol. Cell 78, 779–784.e5. doi:10.1016/j.molcel.2020.04.022.

Hoffmann, M., Kleine-Weber, H., Schroeder, S., Krüger, N., Herrler, T., Erichsen, S., et al. (2020b). SARS-CoV-2 Cell Entry Depends on ACE2 and TMPRSS2 and Is Blocked by a Clinically Proven Protease Inhibitor. Cell 181, 271–280.e8. doi:10.1016/j.cell.2020.02.052.

Hofmann-Winkler, H., Moerer, O., Alt-Epping, S., Bräuer, A., Büttner, B., Müller, M., et al. (2020). Camostat Mesylate May Reduce Severity of Coronavirus Disease 2019 Sepsis: A First Observation. Crit. care Explor. 2, e0284–e0284. doi:10.1097/CCE.0000000000000284.

Huang, H., Guan, L., Yang, Y., Grange, J. M. Le, Tang, G., Xu, Y., et al. (2020). Chloroquine, arbidol (umifenovir) or lopinavir/ritonavir as the antiviral monotherapy for COVID-19 patients: a retrospective cohort study. Res. Sq. doi:10.21203/rs.3.rs-24667/v1.

Kitagawa, J., Arai, H., Iida, H., Mukai, J., Furukawa, K., Ohtsu, S., et al. (2021). A phase I study of high dose camostat mesylate in healthy adults provides a rationale to repurpose the TMPRSS2 inhibitor for the treatment of COVID-19. Clin. Transl. Sci. n/a. doi:https://doi.org/10.1111/cts.13052.

Knoops, K., Kikkert, M., Van Den Worm, S. H. E., Zevenhoven-Dobbe, J. C., Van Der Meer, Y., Koster, A. J., et al. (2008). SARS-coronavirus replication is supported by a reticulovesicular network of modified endoplasmic reticulum. PLoS Biol. 6, 1957–1974. doi:10.1371/journal.pbio.0060226.

Lan, J., Ge, J., Yu, J., Shan, S., Zhou, H., Fan, S., et al. (2020). Structure of the SARS-CoV-2 spike receptor-binding domain bound to the ACE2 receptor. Nature 581, 215–220. doi:10.1038/s41586-020-2180-5.

Letko, M., Marzi, A., and Munster, V. (2020). Functional assessment of cell entry and receptor usage for SARS-CoV-2 and other lineage B betacoronaviruses. Nat. Microbiol. 5, 562–569. doi:10.1038/s41564-020-0688-y.

Li, H., Li, C., Wang, G.-L., Poirier, M. G., and Huang, K. Multiple Ligand Simultaneous Docking (MLSD) and Its Applications to Fragment Based Drug Design and Drug Repositioning DISSERTATION.

Lian, N., Xie, H., Lin, S., Huang, J., Zhao, J., and Lin, Q. (2020). Umifenovir treatment is not associated with improved outcomes in patients with coronavirus disease 2019: a retrospective study. Clin. Microbiol. Infect. 26, 917–921. doi:10.1016/j.cmi.2020.04.026.

Murgolo, N., Therien, A. G., Howell, B., Klein, D., Koeplinger, K., Lieberman, L. A., et al. (2021). SARS-CoV-2 tropism, entry, replication, and propagation: Considerations for drug discovery and development. PLOS Pathog. 17, 1–18. doi:10.1371/journal.ppat.1009225.

Naesens, L., Guddat, L. W., Keough, D. T., Van Kuilenburg, A. B. P., Meijer, J., Vande Voorde, J., et al. (2013). Role of human hypoxanthine guanine phosphoribosyltransferase in activation of the antiviral agent T-705 (favipiravir). Mol. Pharmacol. 84, 615–629. doi:10.1124/mol.113.087247.

Pécheur, E. I., Lavillette, D., Alcaras, F., Molle, J., Boriskin, Y. S., Roberts, M., et al. (2007). Biochemical mechanism of hepatitis C virus inhibition by the broad-spectrum antiviral arbidol. Biochemistry 46, 6050–6059. doi:10.1021/bi700181j.

Pettersen, E. F., Goddard, T. D., Huang, C. C., Couch, G. S., Greenblatt, D. M., Meng, E. C., et al. (2004). UCSF Chimera - A visualization system for exploratory research and analysis. J. Comput. Chem. 25, 1605–1612. doi:10.1002/jcc.20084.

Trott, O., and Olson, A. J. (2009). AutoDock Vina: Improving the speed and accuracy of docking with a new scoring function, efficient optimization, and multithreading. J. Comput. Chem. 31, NA-NA. doi:10.1002/jcc.21334.

Uno, Y. (2020). Camostat mesilate therapy for COVID-19. Intern. Emerg. Med. 15, 1577–1578. doi:10.1007/s11739-020-02345-9.

V’kovski, P., Kratzel, A., Steiner, S., Stalder, H., and Thiel, V. (2020). Coronavirus biology and replication: implications for SARS-CoV-2. Nat. Rev. Microbiol. doi:10.1038/s41579-020-00468-6.

Vankadari, N. (2020). Arbidol: A potential antiviral drug for the treatment of SARS-CoV-2 by blocking trimerization of the spike glycoprotein. Int. J. Antimicrob. Agents 56, 105998. doi:10.1016/j.ijantimicag.2020.105998.

Wang, M., Cao, R., Zhang, L., Yang, X., Liu, J., Xu, M., et al. (2020a). Remdesivir and chloroquine effectively inhibit the recently emerged novel coronavirus (2019-nCoV) in vitro. Cell Res. 30, 269–271. doi:10.1038/s41422-020-0282-0.

Wang, X., Cao, R., Zhang, H., Liu, J., Xu, M., Hu, H., et al. (2020b). The anti-influenza virus drug, arbidol is an efficient inhibitor of SARS-CoV-2 in vitro. Cell Discov. 6, 4–8. doi:10.1038/s41421-020-0169-8.

Yuan, Y., Pei, J., and Lai, L. (2013). Binding Site Detection and Druggability Prediction of Protein Targets for Structure-Based Drug Design. Curr. Pharm. Des. 19, 2326–2333. doi:10.2174/1381612811319120019.

