## Supplemental Raw Data for "Favipiravir, umifenovir and camostat mesylate: a comparative study against SARS-CoV-2": ArbiComFavi_supplementary_MAU.docx

| 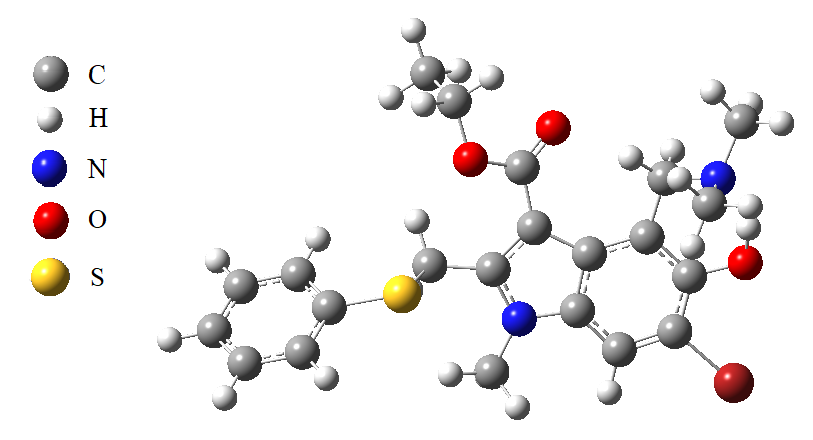 | 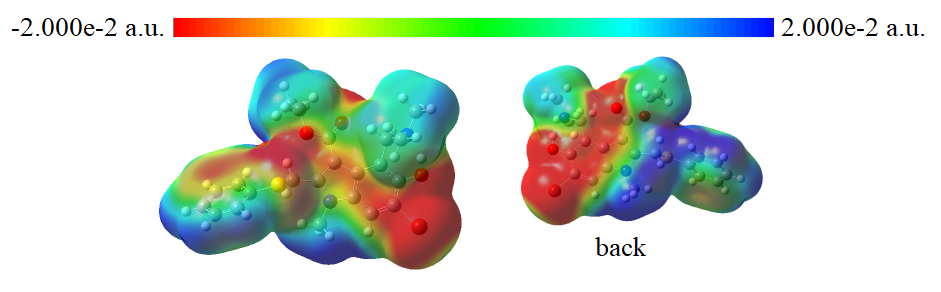 |
| --- | --- |
| A | |
| 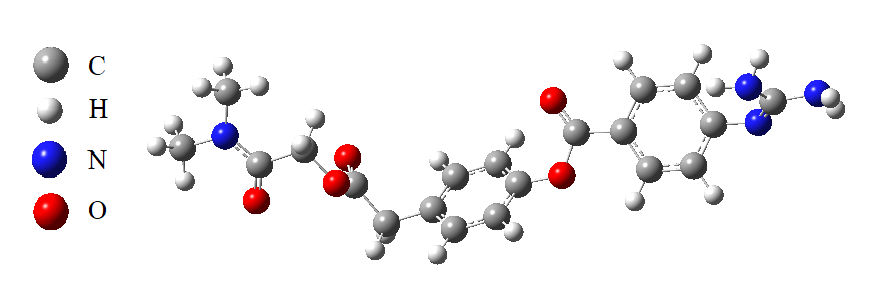 | 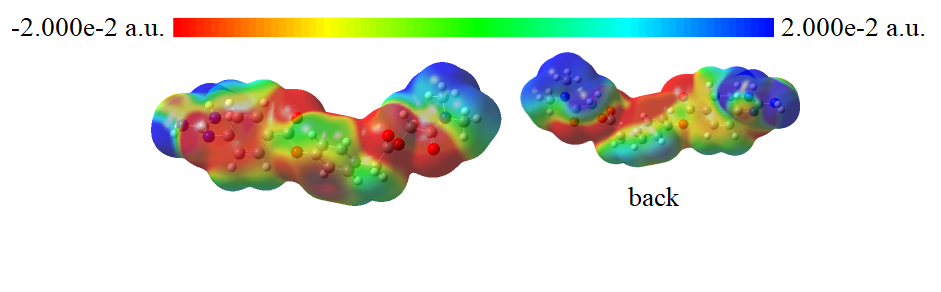 |
| B | |
| 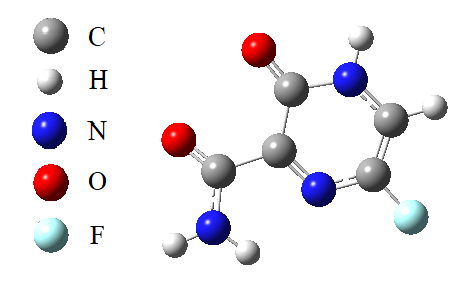 | 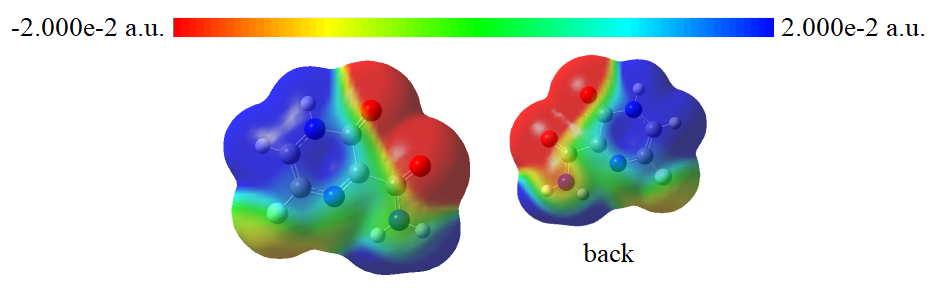 |
| C | |
| **Figure S1 –** Minimized structures and electrostatic potential (ESP) maps of A) Umifenovir, calculated electronic energy (E)= -4158.479720 Hartree (-113157.988398 eV) and dipole moment (μ)=5.277 Debye, B) Camostat Mesylate, calculated electronic energy (E)= -1370.541113 Hartree (-37294.320474 eV) and dipole moment (μ)=6.708 Debye, C) Favipiravir, calculated electronic energy (E)= -607.475744 Hartree (-16530.255723 eV) and dipole moment (μ)=5.297 Debye. | |

| 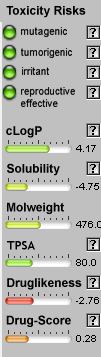 | 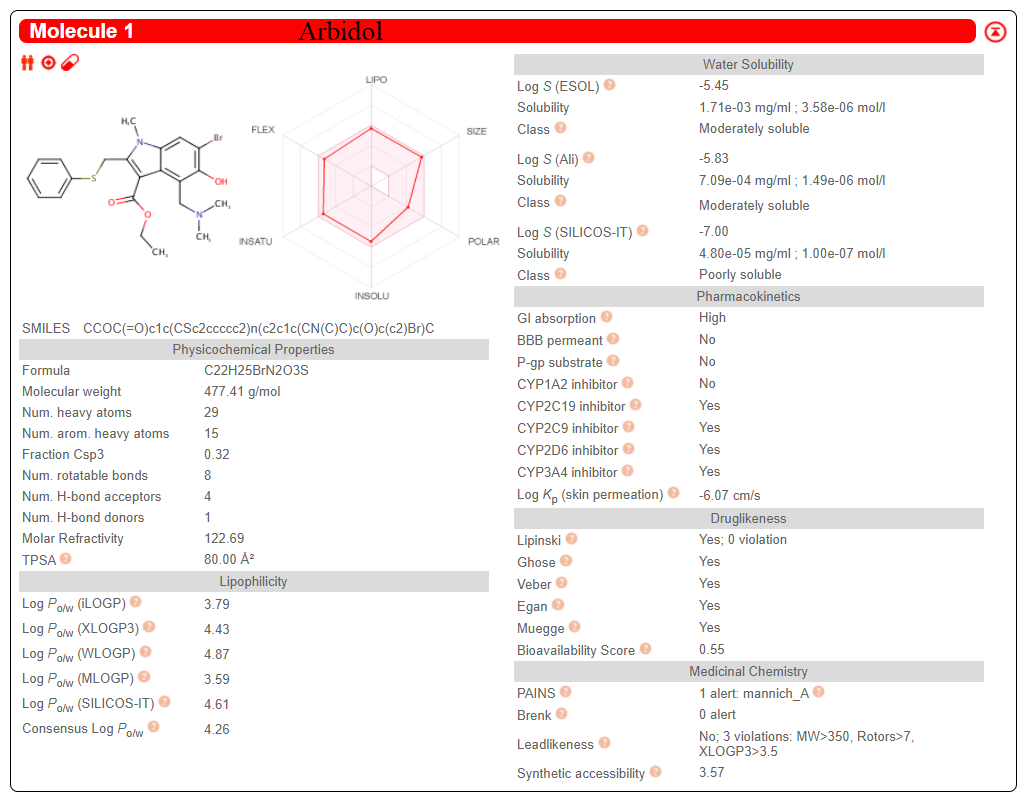 |
| --- | --- |
| A | |
| 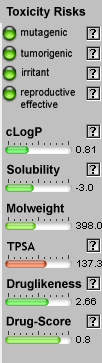 | 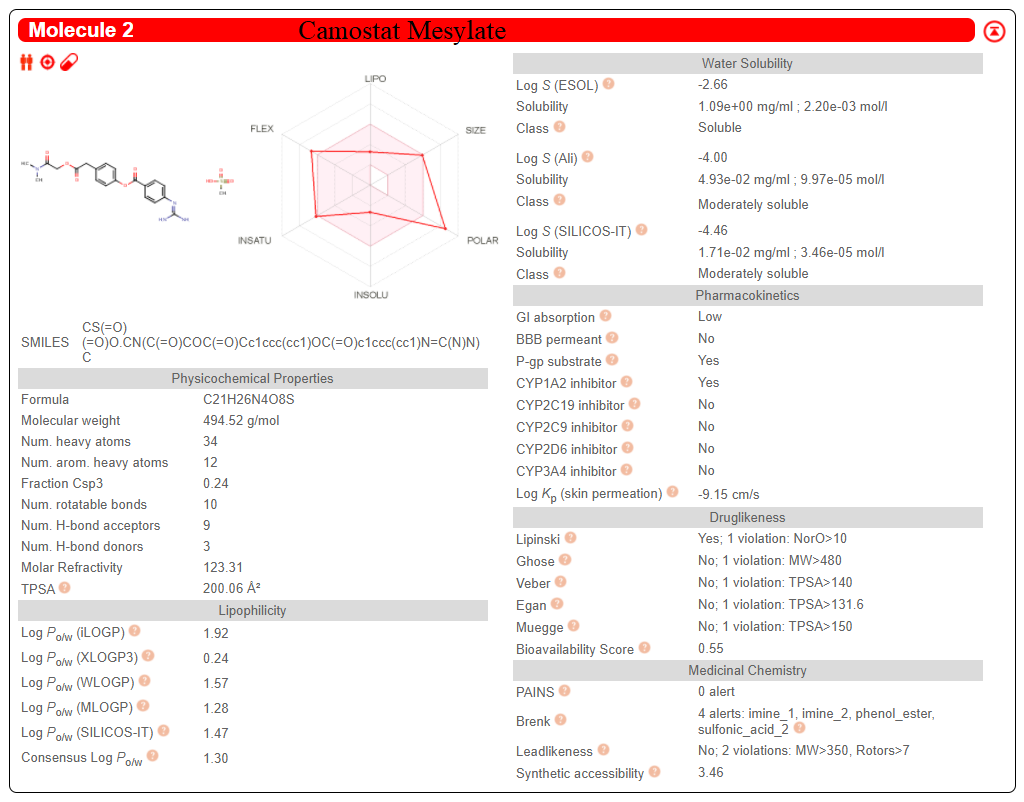 |
| B | |
| 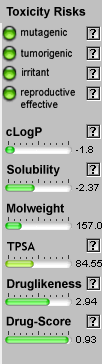 | 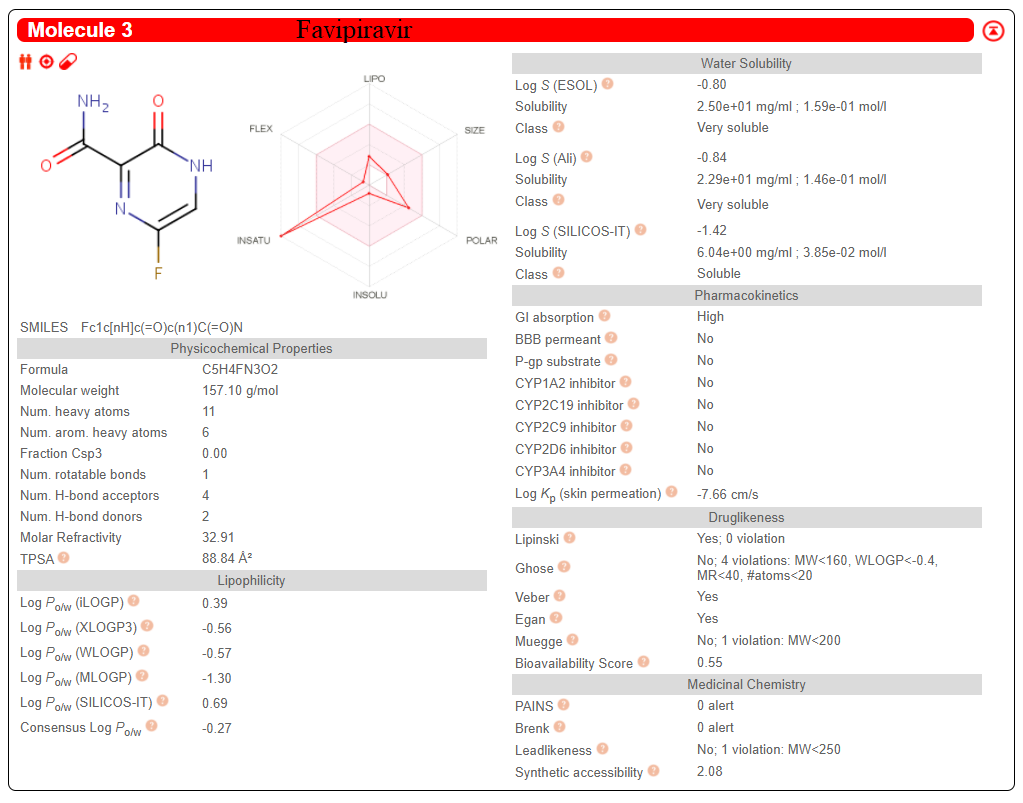 |
| C | |
| **Figure S2 –** ADME-T calculation results of A) Umifenovir, B) Camostat Mesylate and C) Favipiravir. All three drug- active chemicals do not show mutagenicity. Calculated properties are within acceptable values. | |

**Table S1 –**Highest affinity values of single and duo docking results of umifenovir, camostat mesylate and favipiravir-protein complexes (kcal/mol).

| **Protein ID** | **Umifenovir**  **ΔG*, kcal/mol** | **Camostat Mesylate**  **ΔG*, kcal/mol** | **Favipiravir**  **ΔG*, kcal/mol** |
| --- | --- | --- | --- |
| **Single** | | | |
| 1R42 | -7.17 | -7.99 | -5.25 |
| 5X29 | -3.99 | -4.55 | -2.97 |
| 6LU7 | -8.35 | -7.21 | -5.48 |
| 6LXT | -5.22 | -5.34 | -4.74 |
| 6M0J | -6.56 | -6.42 | -4.52 |
| 6M03 | -6.14 | -7.54 | -4.46 |
| 6M71 | -6.32 | -6.52 | -5.33 |
| 6VWW | -6.35 | -7.70 | -4.98 |
| 6VXX | -5.84 | -6.87 | -5.54 |
| 6VYB | -5.90 | -7.57 | -5.33 |
| 6VYO | -5.94 | -7.97 | -5.11 |
| 6Y84 | -6.95 | -6.31 | -4.44 |
| **Protein ID** | **Umifenovir + Camostat M.**  **ΔG*, kcal/mol** | **Camostat M. + Favipiravir**  **ΔG*, kcal/mol** | **Favipiravir + Umifenovir**  **ΔG*, kcal/mol** |
| **Duo** | | | |
| 1R42 | -6.55 + -6.94 | -7.65 + -5.51 | -5.23 + -6.99 |
| 5X29 | -3.29 + -4.29 | -4.75 + -3.46 | -3.20 + -3.52 |
| 6LU7 | -8.11 + -6.59 | -7.28 + -5.33 | -4.73 + -7.07 |
| 6LXT | -6.40 + -5.46 | -6.21 + -4.74 | -4.72 + -4.92 |
| 6M0J | -7.45 + -7.74 | -8.26 + -5.20 | -5.02 + -7.28 |
| 6M03 | -6.06 + -6.05 | -6.25 + -4.42 | -4.49 + -6.41 |
| 6M71 | -6.26 + -5.71 | -6.54 + -5.55 | -5.15 + -5.77 |
| 6VWW | -6.86 + -6.88 | -7.00 + -4.55 | -4.97 + -6.79 |
| 6VXX | -5.13 + -6.58 | -8.96 + -4.36 | -5.55 + -5.42 |
| 6VYB | -6.12 + -7.47 | -8.63 + 4.85 | -5.06 + -5.87 |
| 6VYO | -5.80 + -8.23 | -8.53 + -4.95 | -5.26 + -6.73 |
| 6Y84 | -6.97 + -6.39 | -6.96 + -4.35 | -4.65 + -5.97 |

*RMSD = 0.00

| 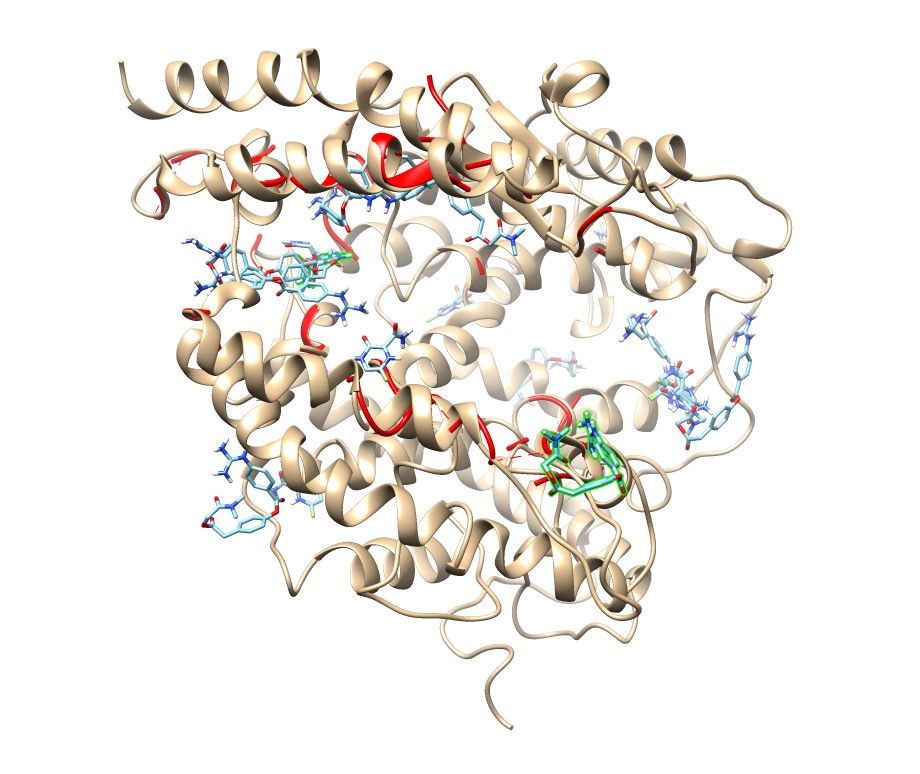  1R42  Druggability score: 6822 | 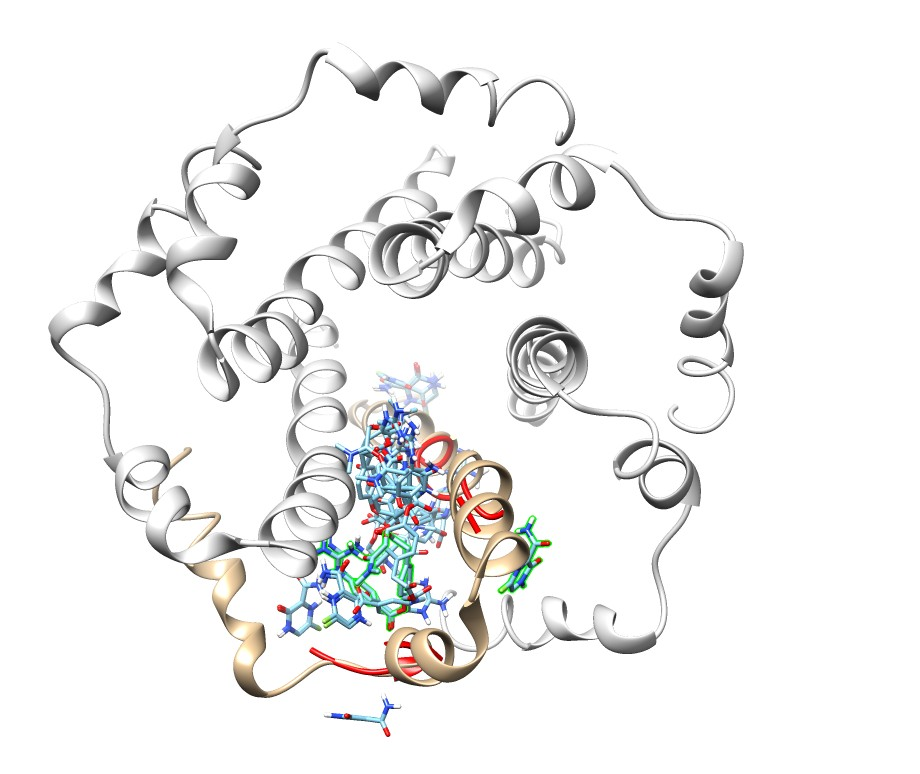  5X29  Druggability score: 4232 |
| --- | --- |
| 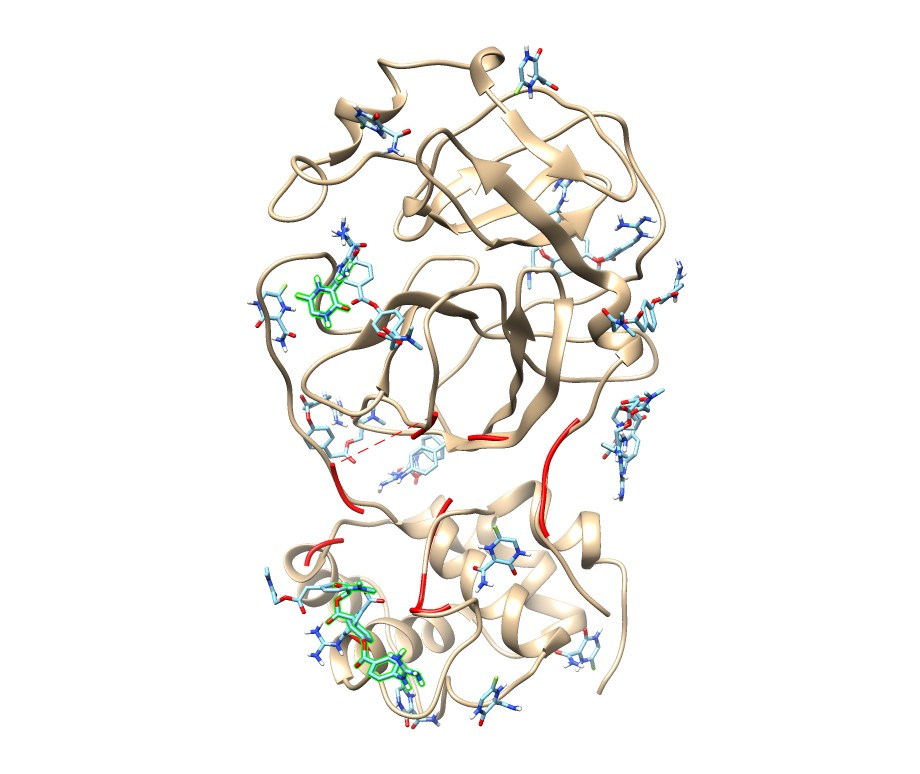  6LU7  Druggability score: -344 | 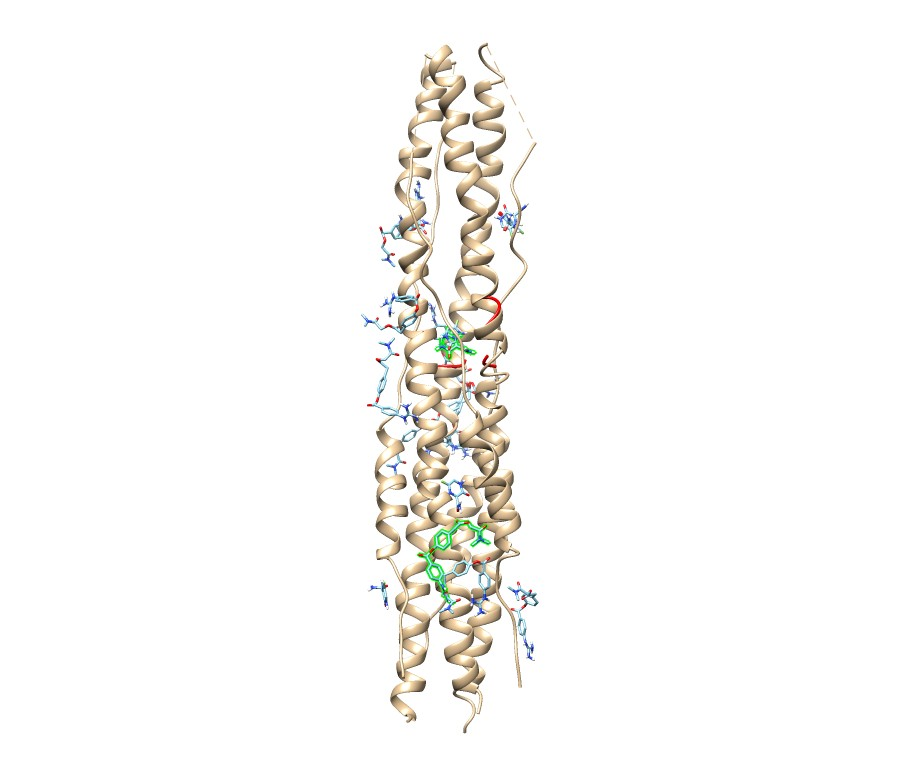  6LXT  Druggability score: 241 |
| 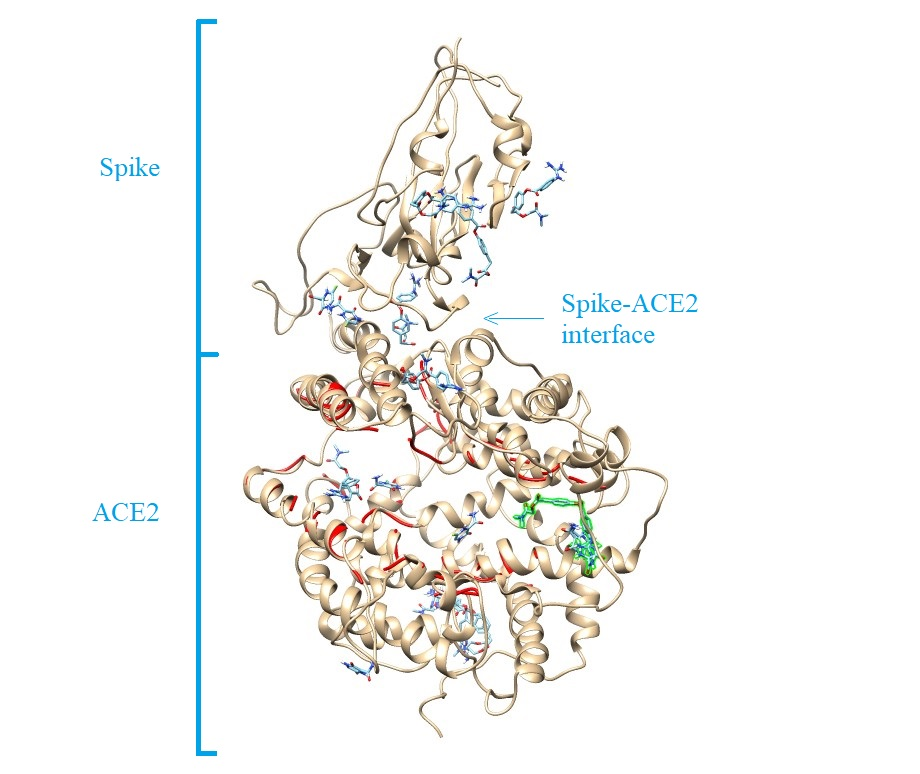  6M0J  Druggability score: 7414 | 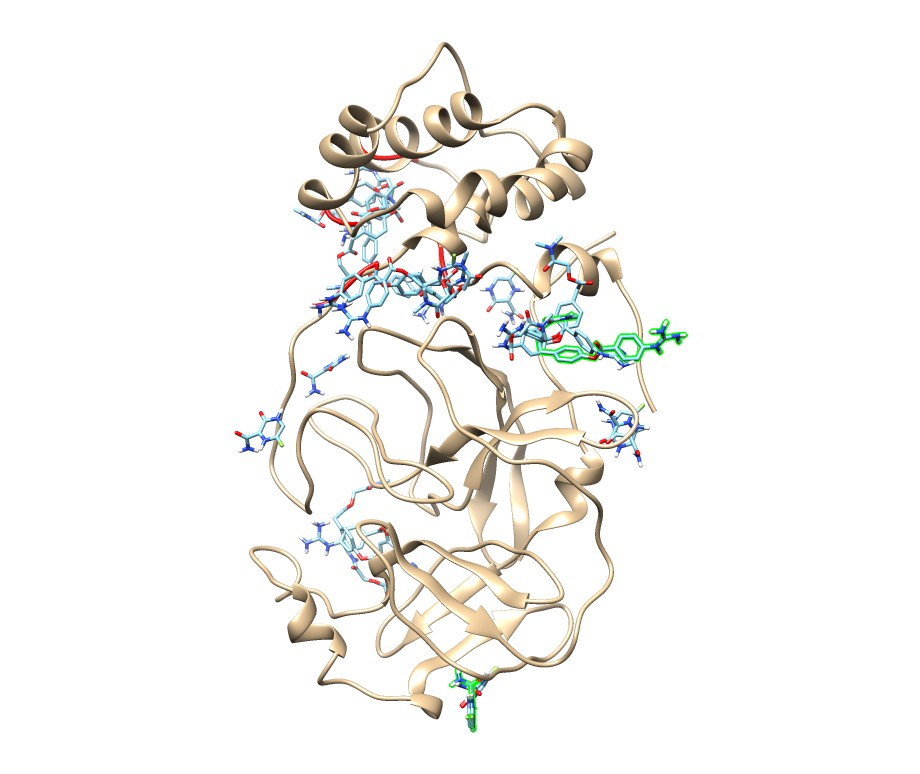  6M03  Druggability score: 51 |
| 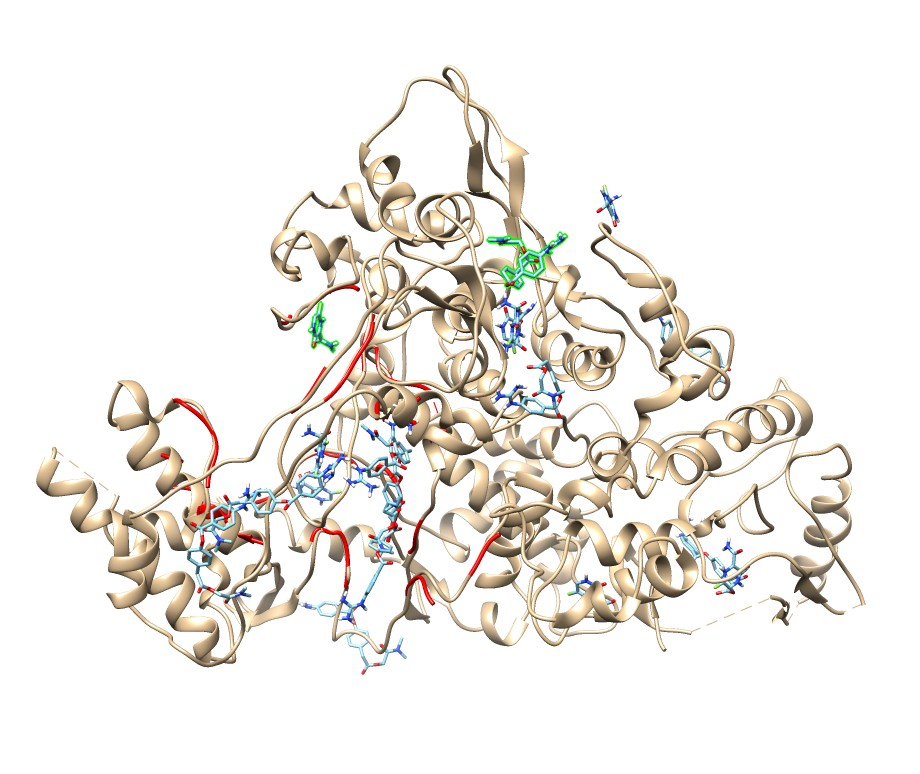  6M71  Druggability score: 2495 | 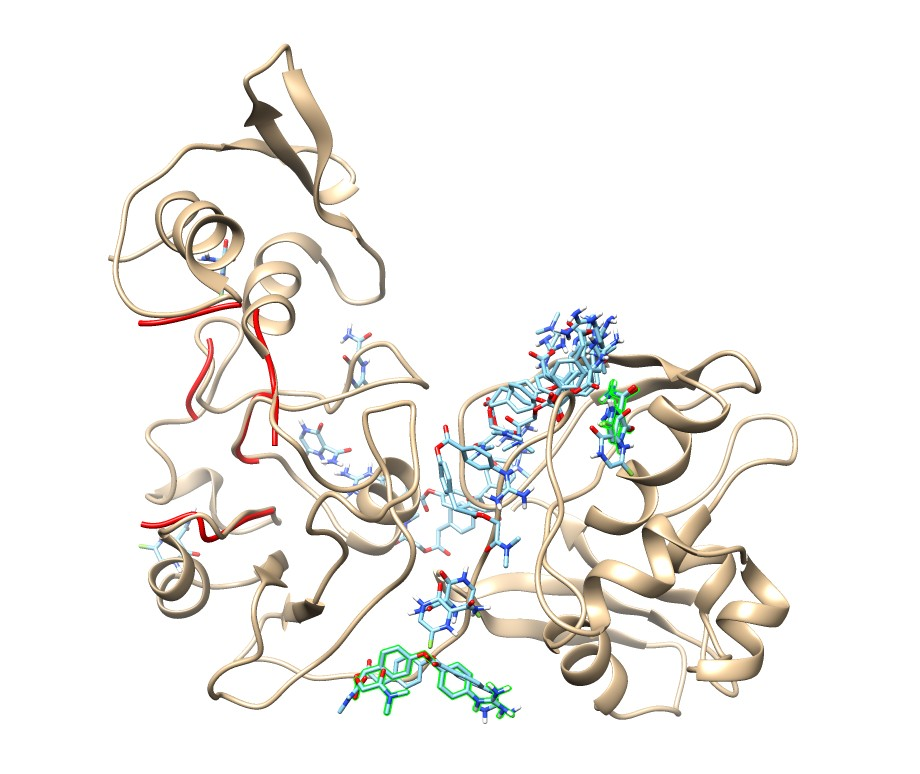  6VWW  Druggability score: -259 |
| 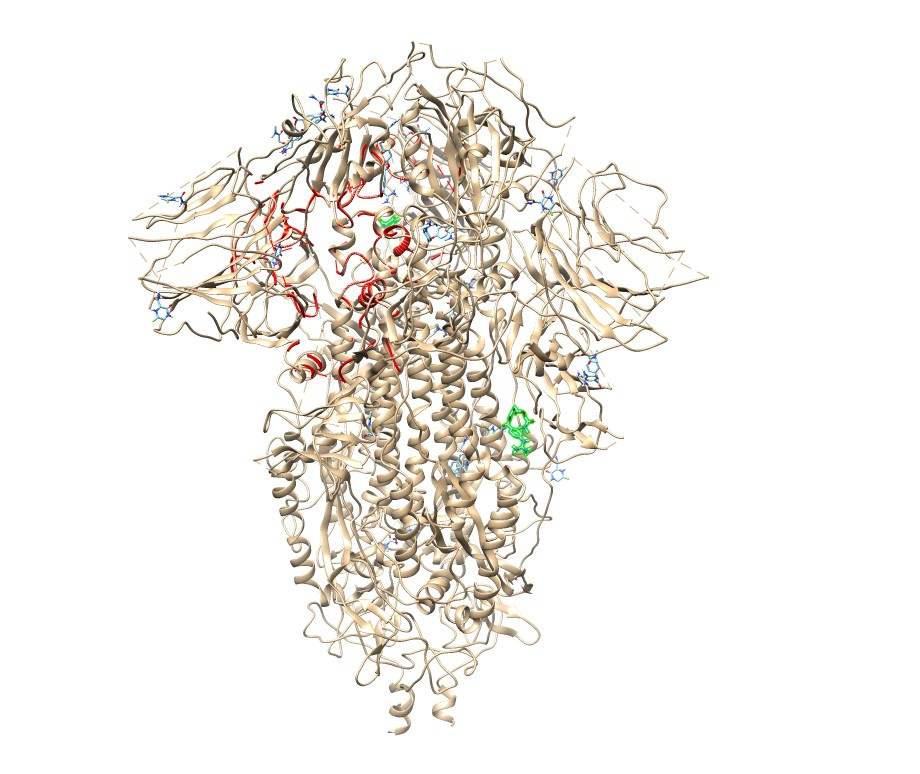  6VXX-spike close form  Druggability score: 6589; this is 3^rd^ druggable score but it is located on the RBD region. The other two scores are higher and located in the middle of the spike (1^rst^ score: 16603, 2^nd^ score: 13701; RawData.zip). | 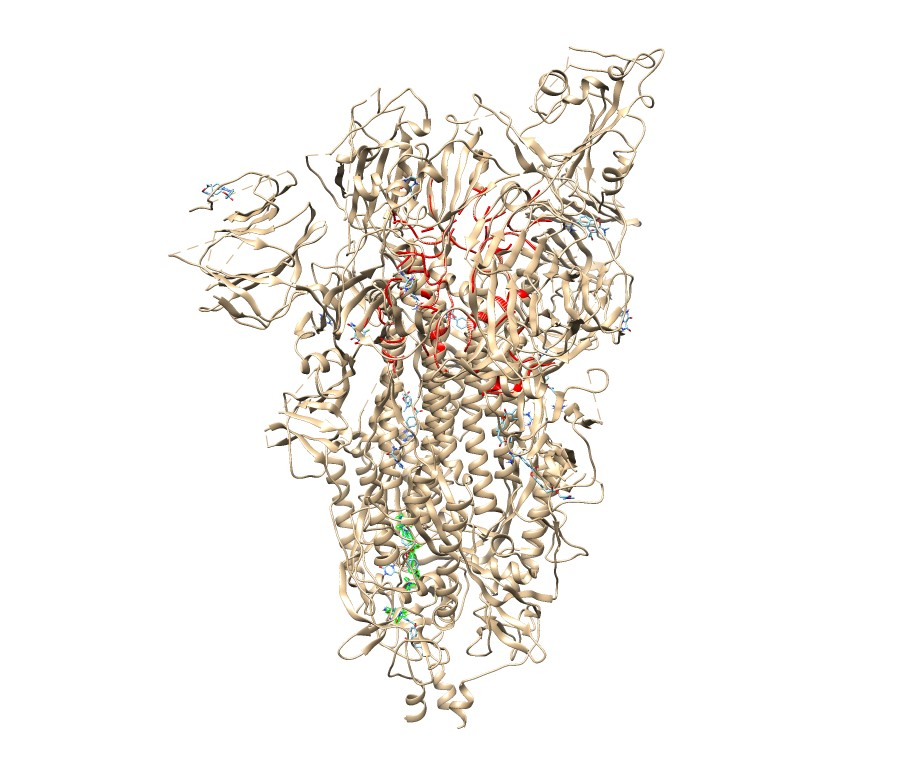  6VYB-spike open form  Druggability score: 4954, this is 3^rd^ druggable score but it is located on the RBD region. The other two scores are higher and located in the middle (1^st^ score: 9561) and in the bottom (2^nd^ score: 5089) of the spike (RawData.zip). |
| 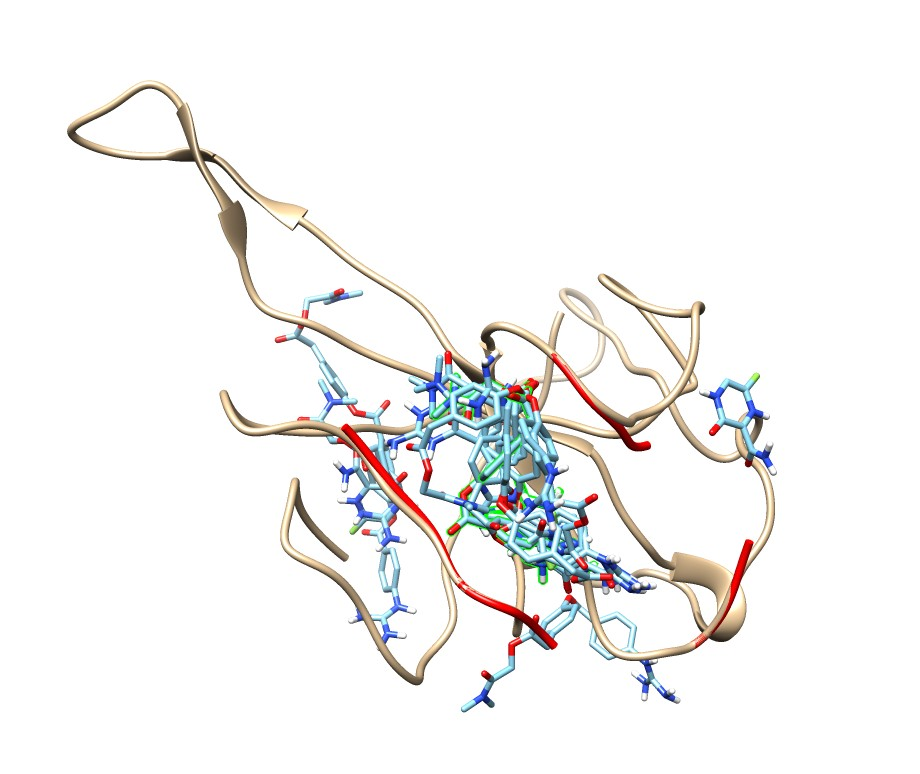  6VYO  Druggability score: -634 | 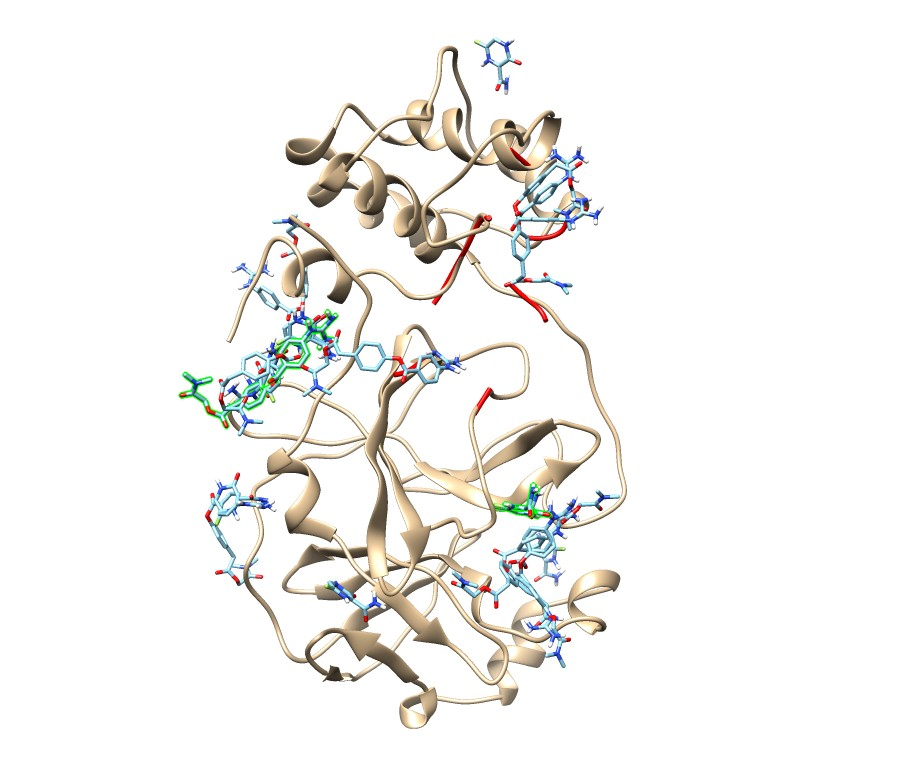  6Y84  Druggability score: -221 |
| **Figure S3 –** Camostat Mesylate+Favipiravir-protein complexes with 10 highest affinity energy conformations. The red colored amino acids indicate the highest energy druggabale cavity and the highest energy conformation is green colored. | |

| 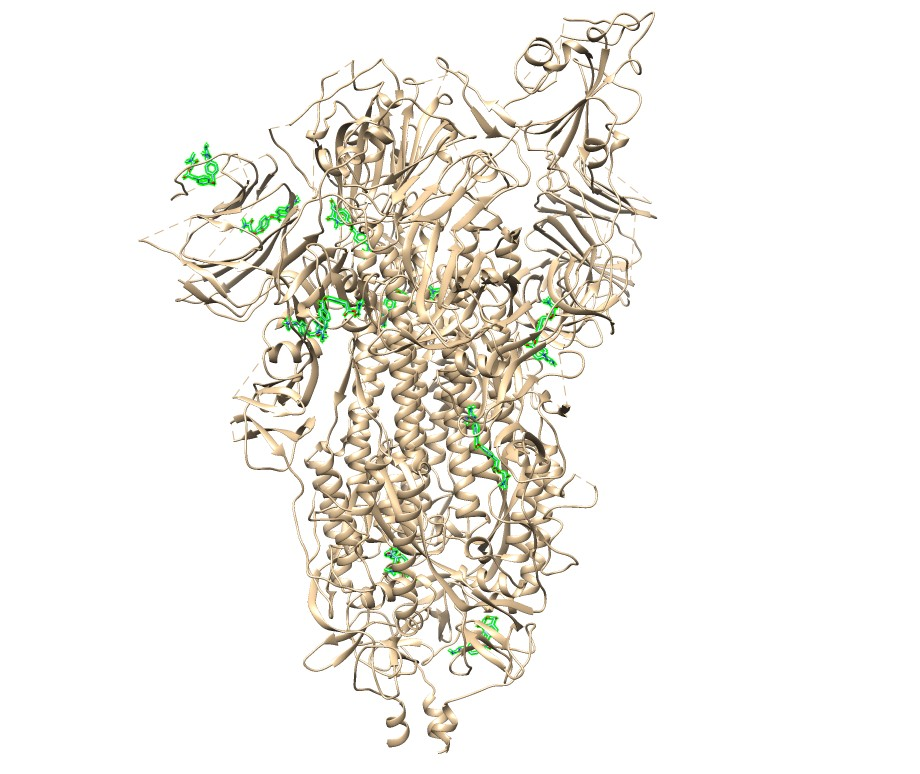  6VYB-Camostat Mesylate | 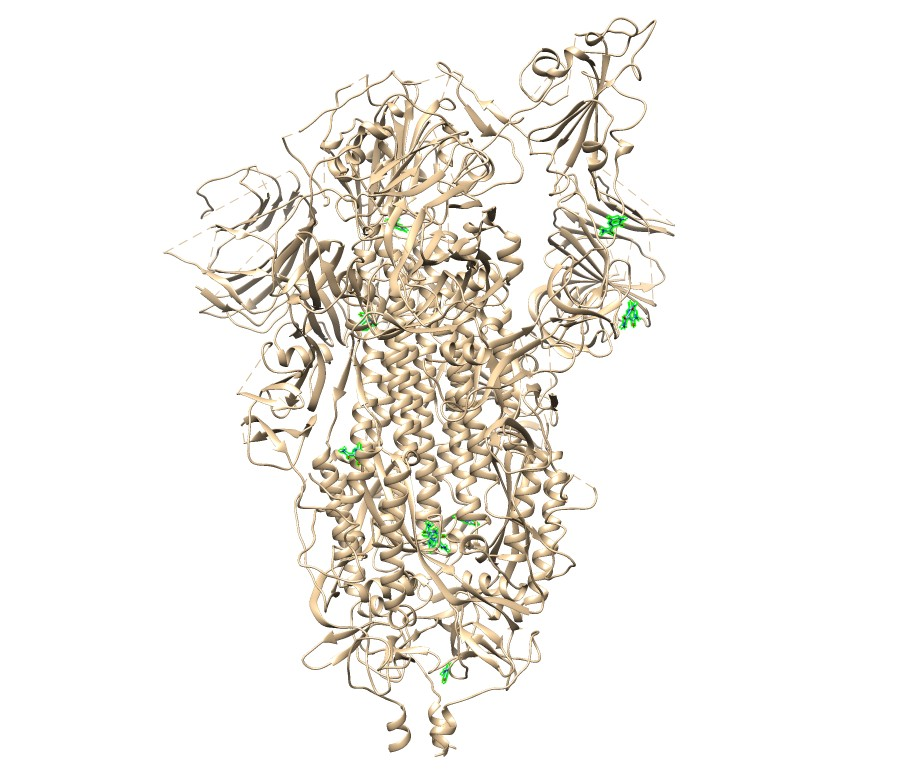  6VYB-Favipiravir |
| --- | --- |
| 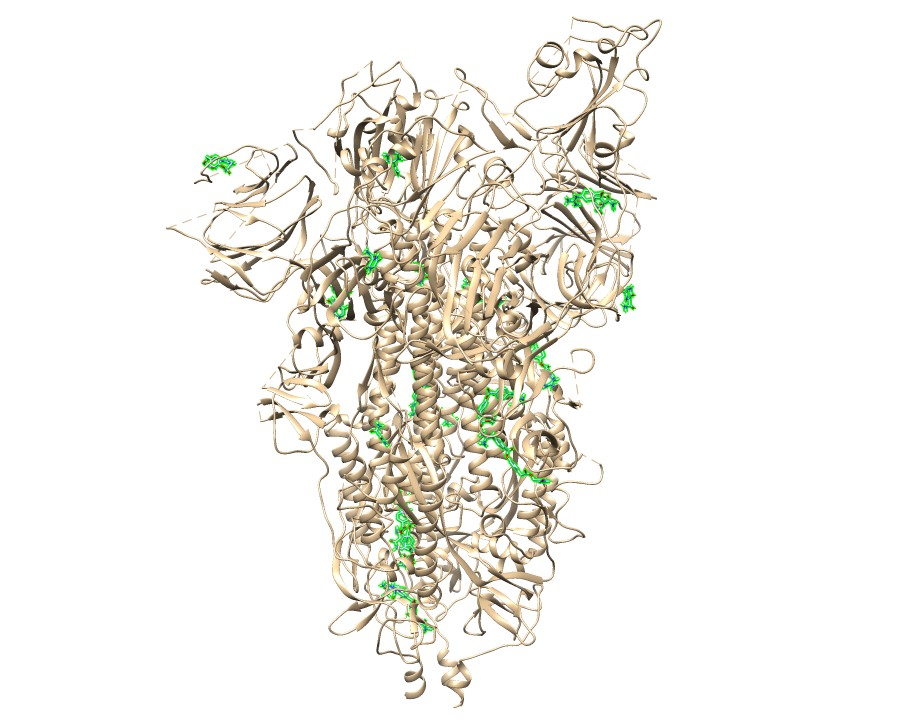  6VYB-Camostat Mesylate+Favipiravir |  |
| 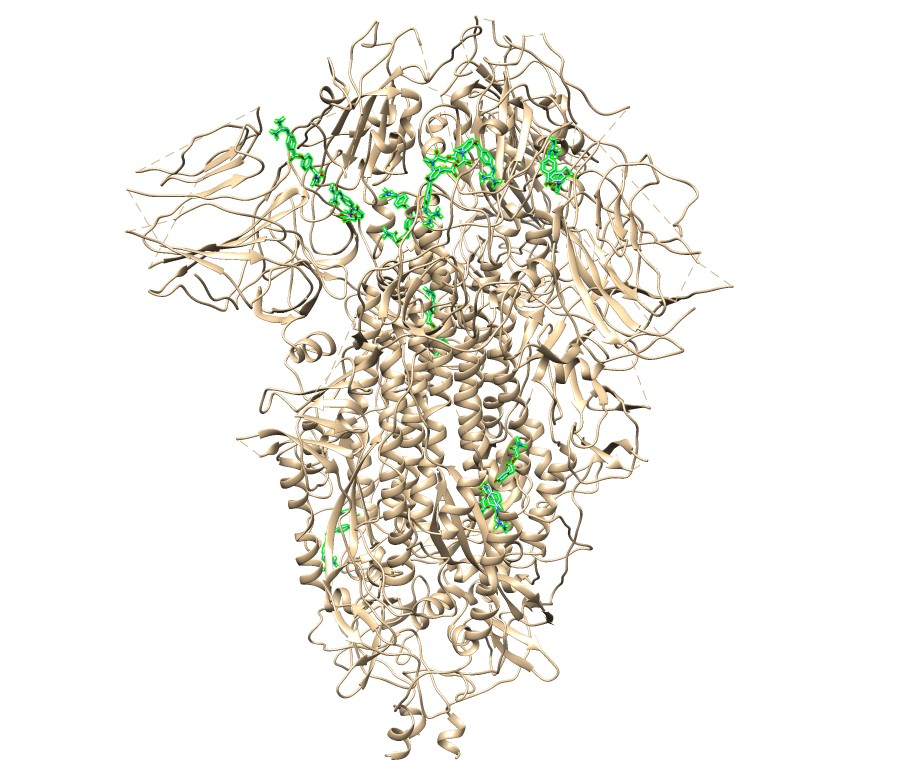  6VXX-Camostat Mesylate | 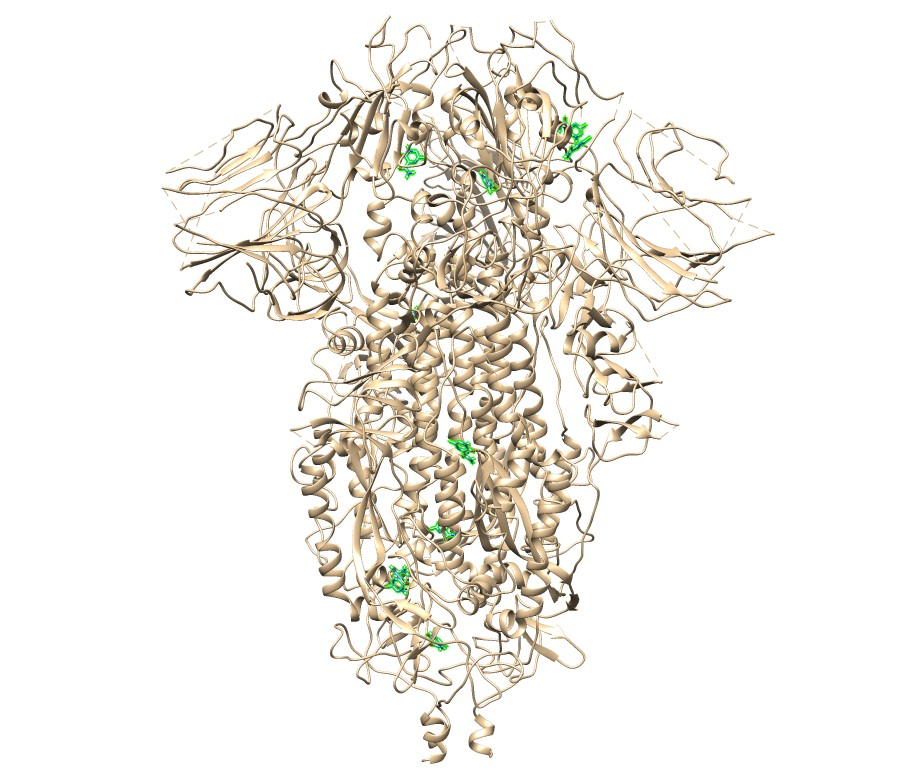  6VXX-Favipiravir |
| 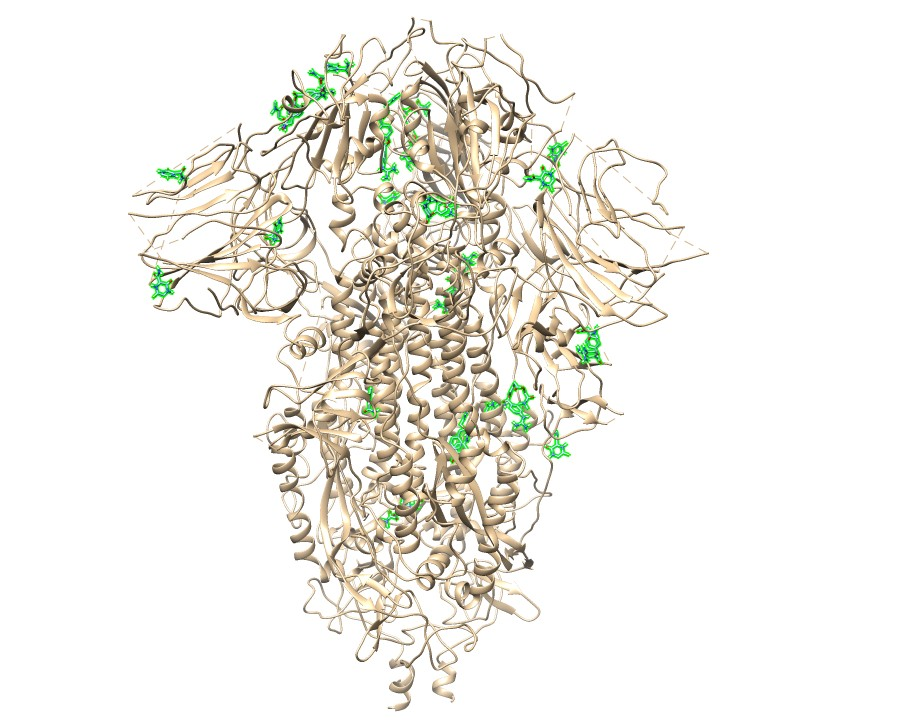  6VXX-Camostat Mesylate+Favipiravir |  |
| 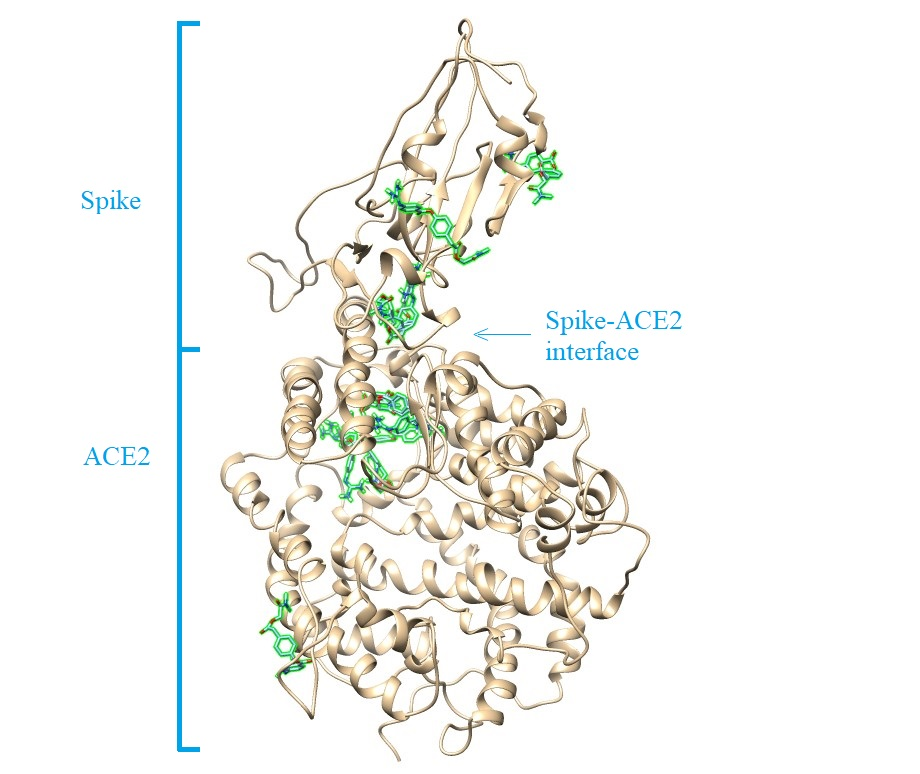  6M0J-Camostat Mesylate | 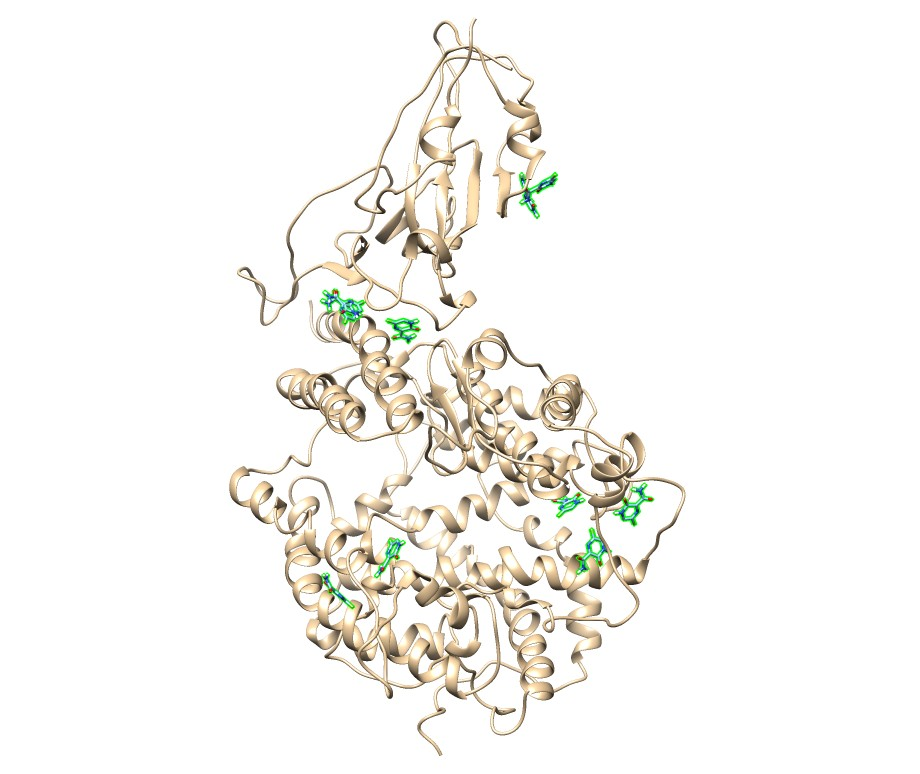  6M0J-Favipiravir |
| 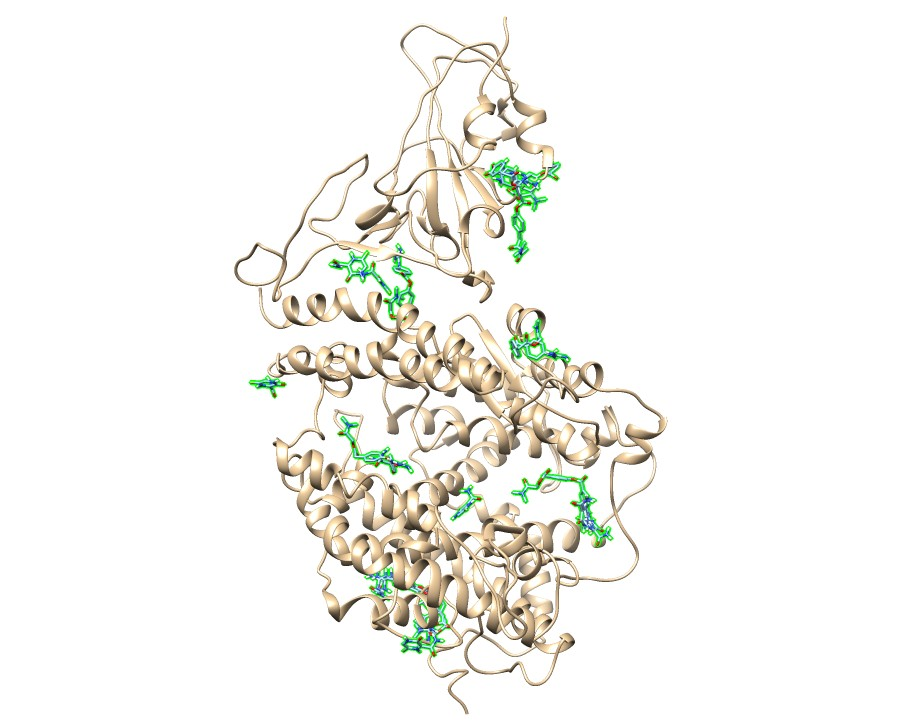  6M0J-Camostat Mesylate+Favipiravir |  |
| **Figure S4 –** 3D shots of camostat mesylate, favipiravir and camostatM+favipiravir-6VYB, 6VXX and 6M0J protein complexes. | |

| 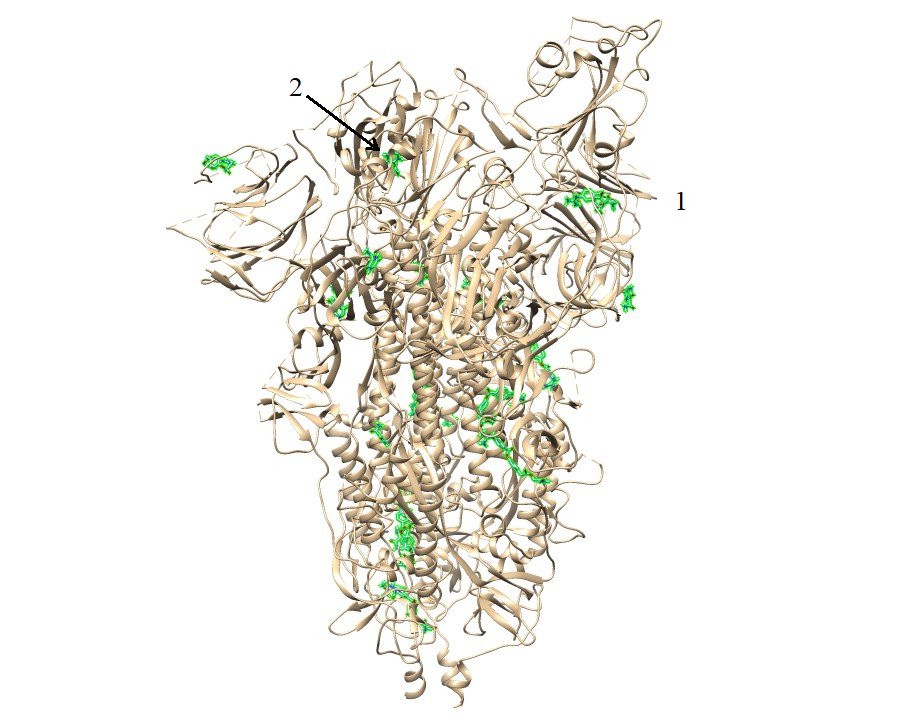  6VYB-Camostat Mesylate+Favipiravir  A | 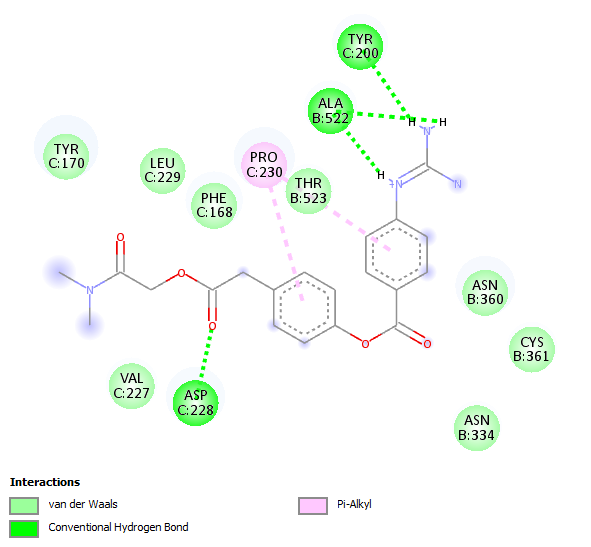  Mode-1 2D |
| --- | --- |
| 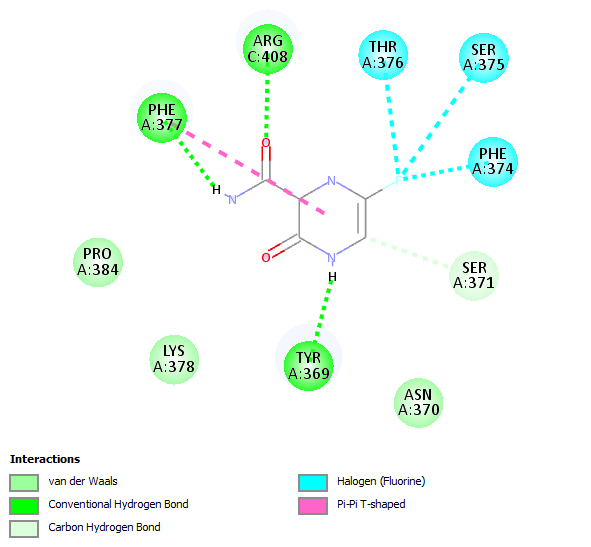  Mode-2 2D |  |
| 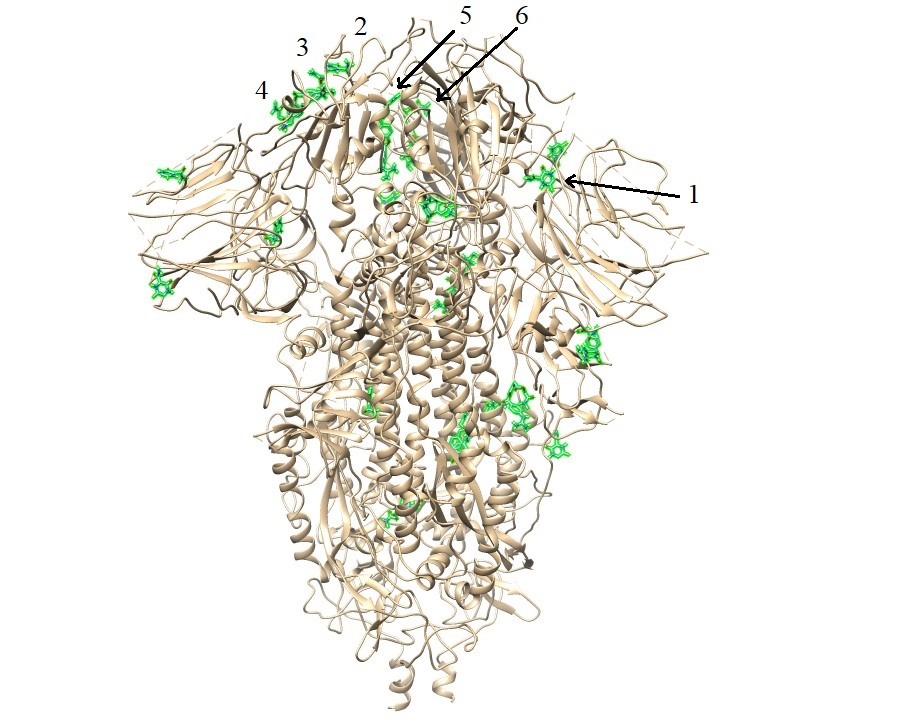  6VXX-Camostat Mesylate+Favipiravir  B | 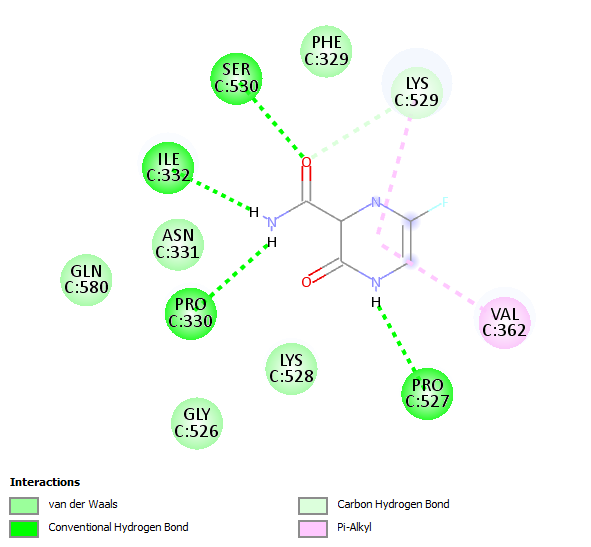  Mode-1 2D |
| 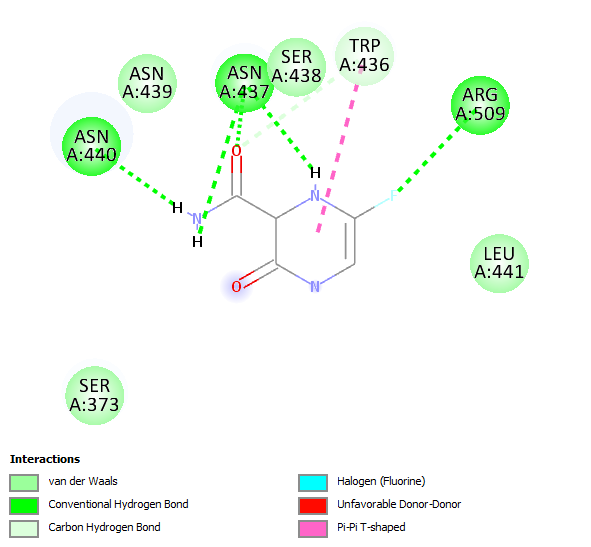  Mode-2 2D | 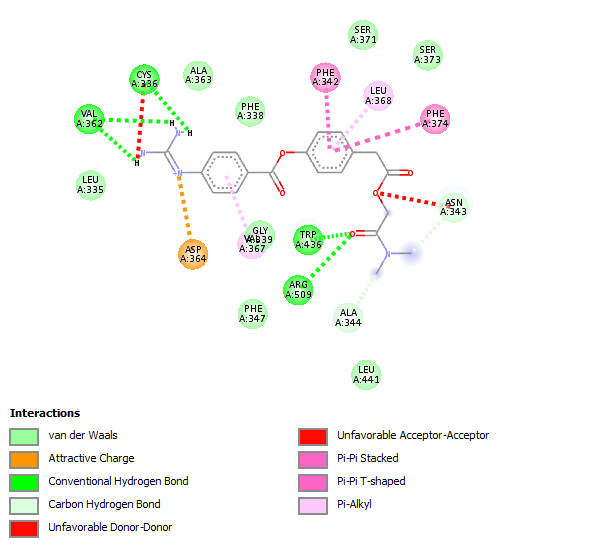  Mode-3 2D |
| 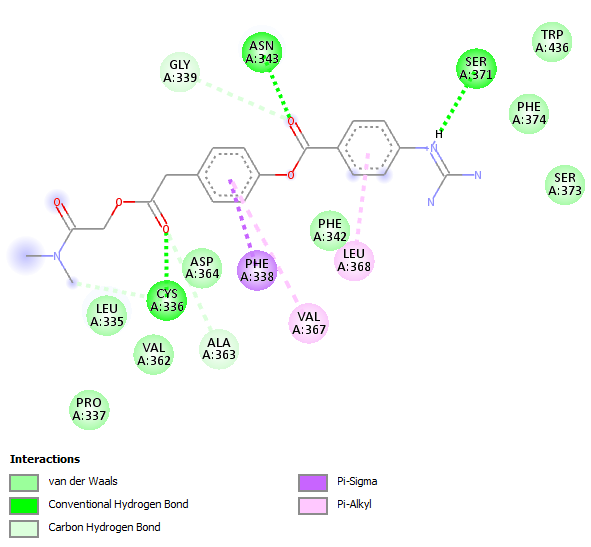  Mode-4 2D | 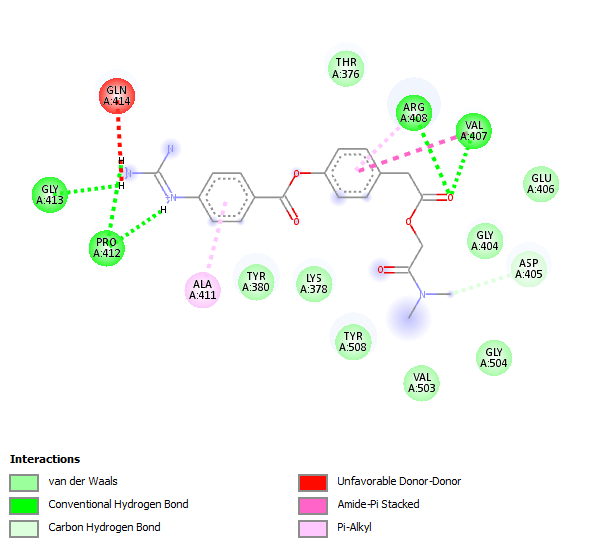  Mode-5 2D |
| 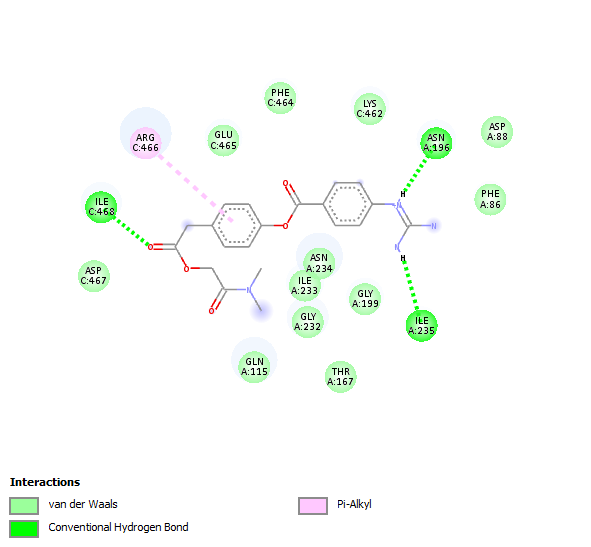  Mode-6 2D |  |
| 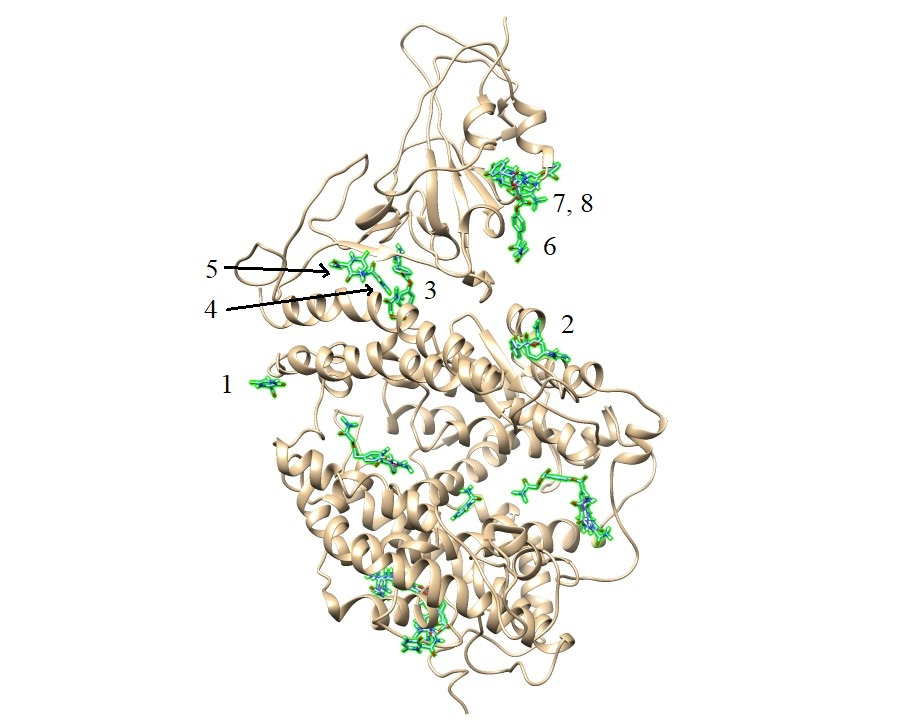  6M0J-Camostat Mesylate+Favipiravir  C | 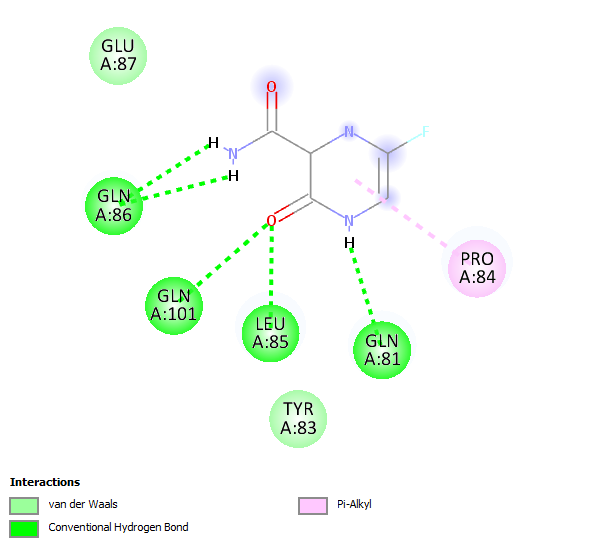  Mode-1 2D |
| 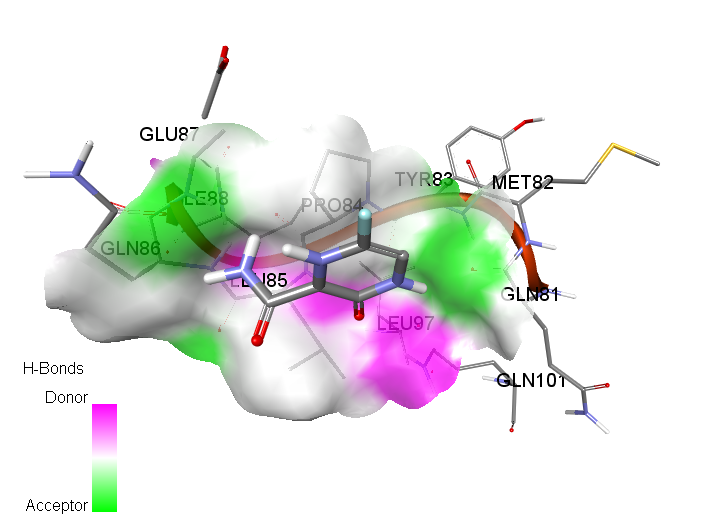  Mode-1 3D-Hydrogen bond | 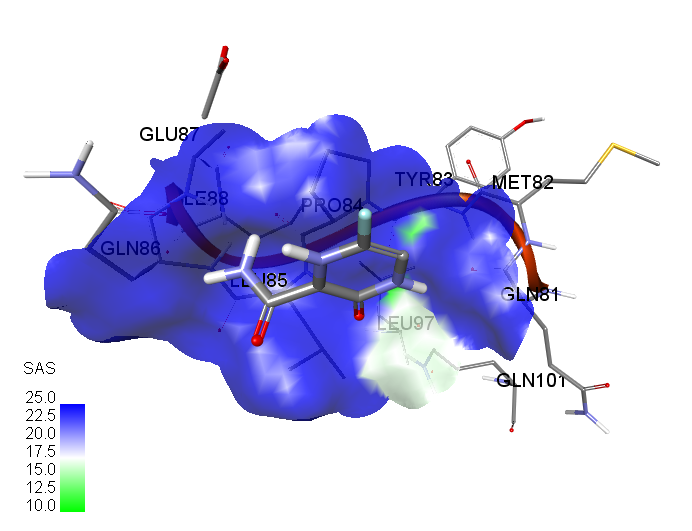  Mode-1 3D-SAS (Solvent Accesibility Surface) |
| 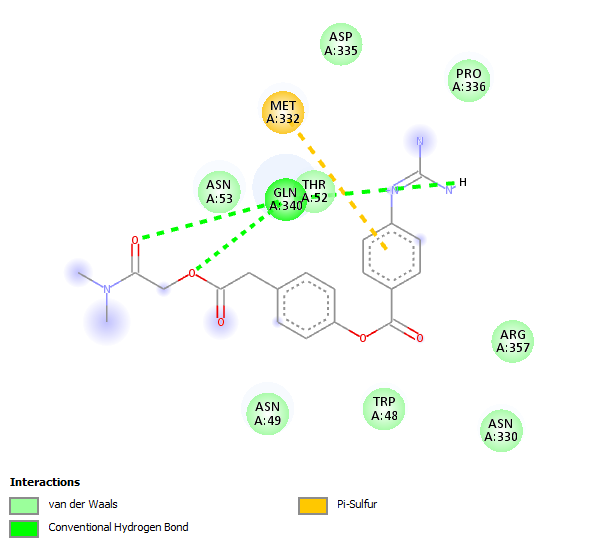  Mode-2 2D | 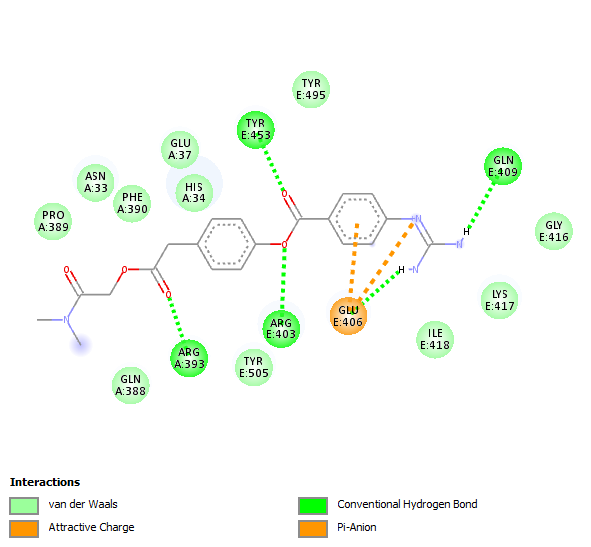  Mode-3 2D |
| 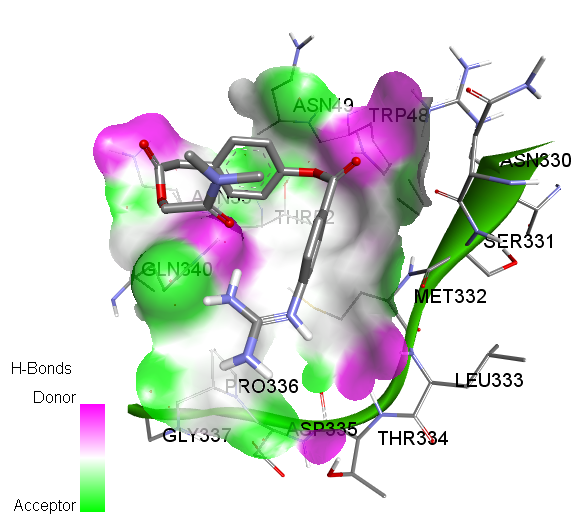  Mode-2 3D-Hydrogen bond | 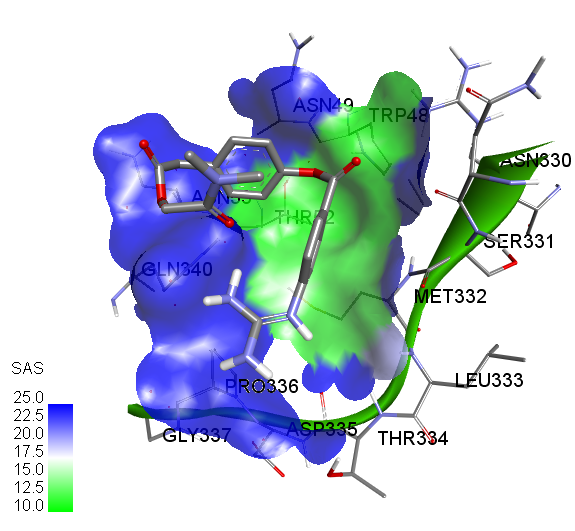  Mode-2 3D-SAS |
| 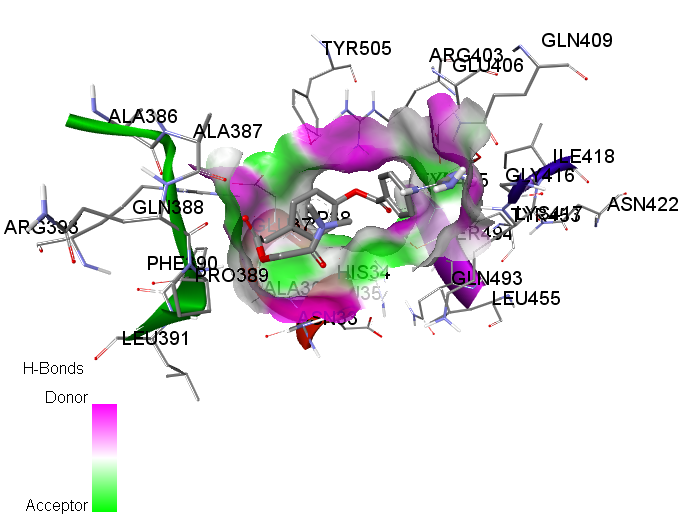  Mode-3 3D-Hydrogen bond | 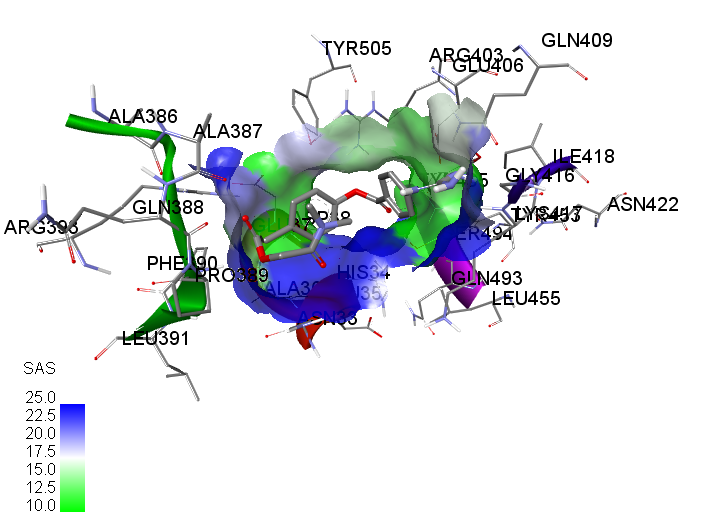  Mode-3 3D-SAS |
| 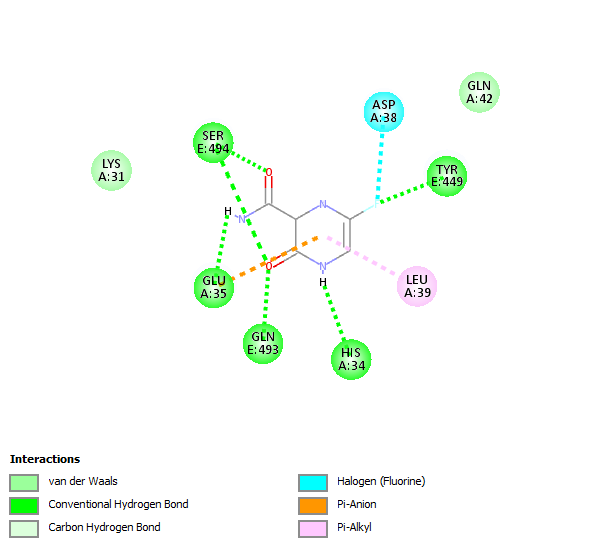  Mode-4 2D | 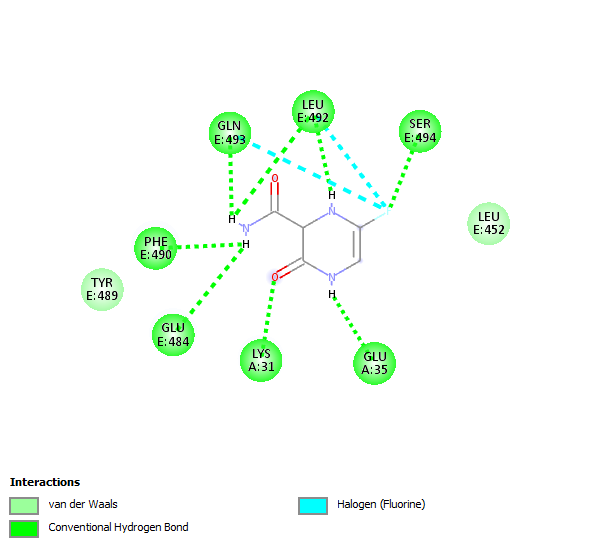  Mode-5 2D |
| 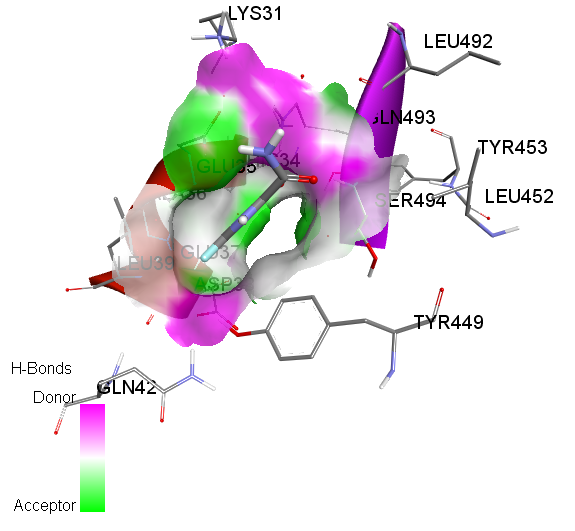  Mode-4 3D-Hydrogen bond | 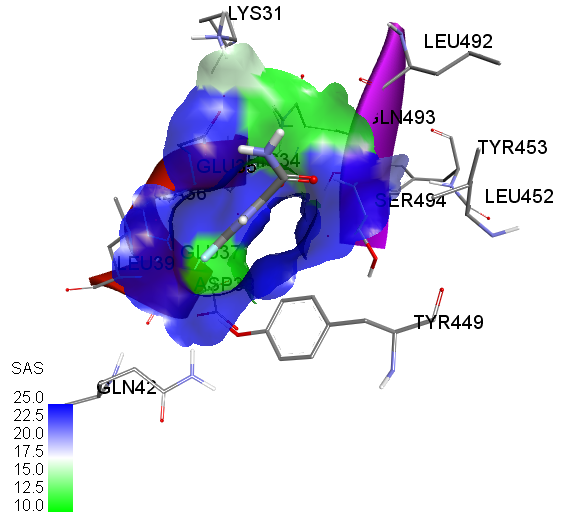  Mode-4 3D-SAS |
| 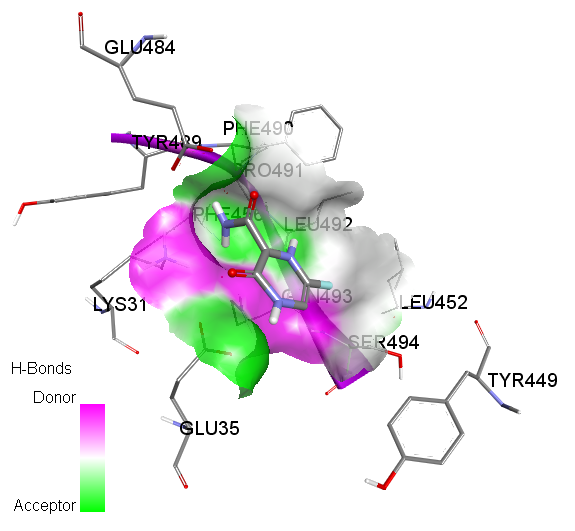  Mode-5 3D-Hydrogen bond | 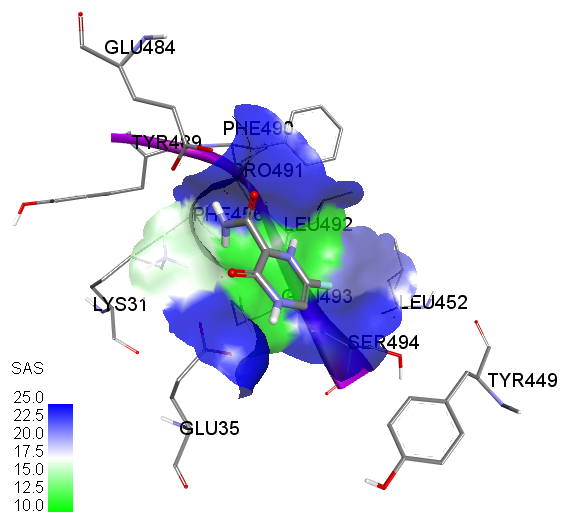  Mode-5 3D-SAS |
| 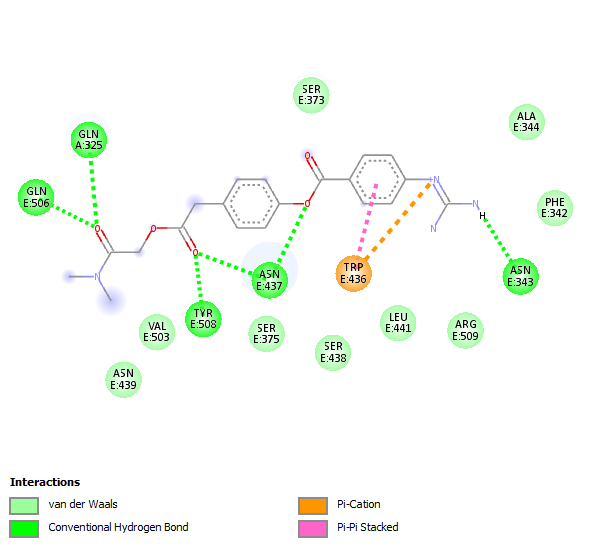  Mode-6 2D | 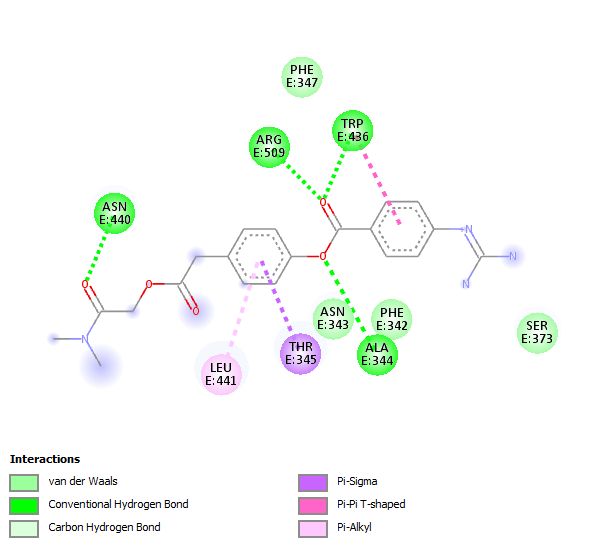  Mode-7 2D |
| 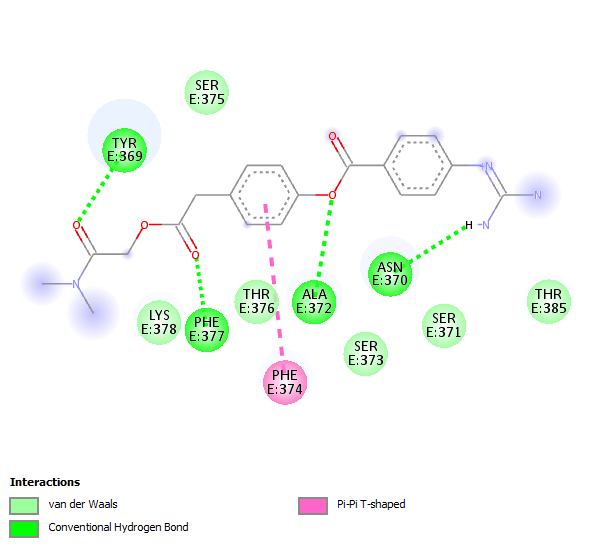  Mode-8 2D |  |
| **Figure S5 –** Affinity of the 10 highest energy conformations of the camostatM+favipiravir combination for the active amino acids of the A) 6VYB, B) 6VXX and C) 6M0J proteins | |

| 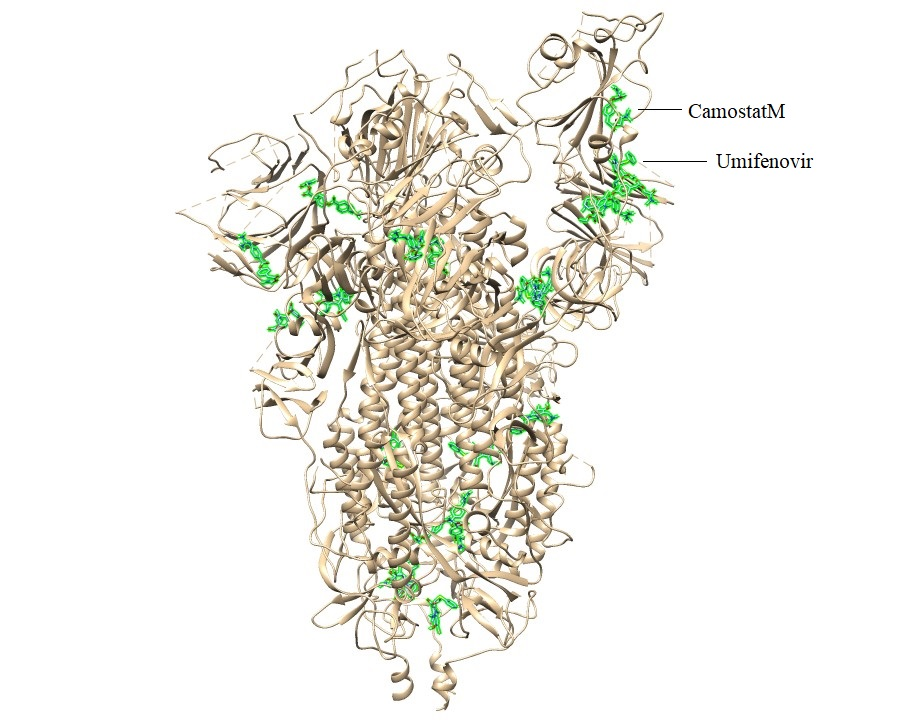  6VYB-Camostat Mesylate+Umifenovir  A1 | 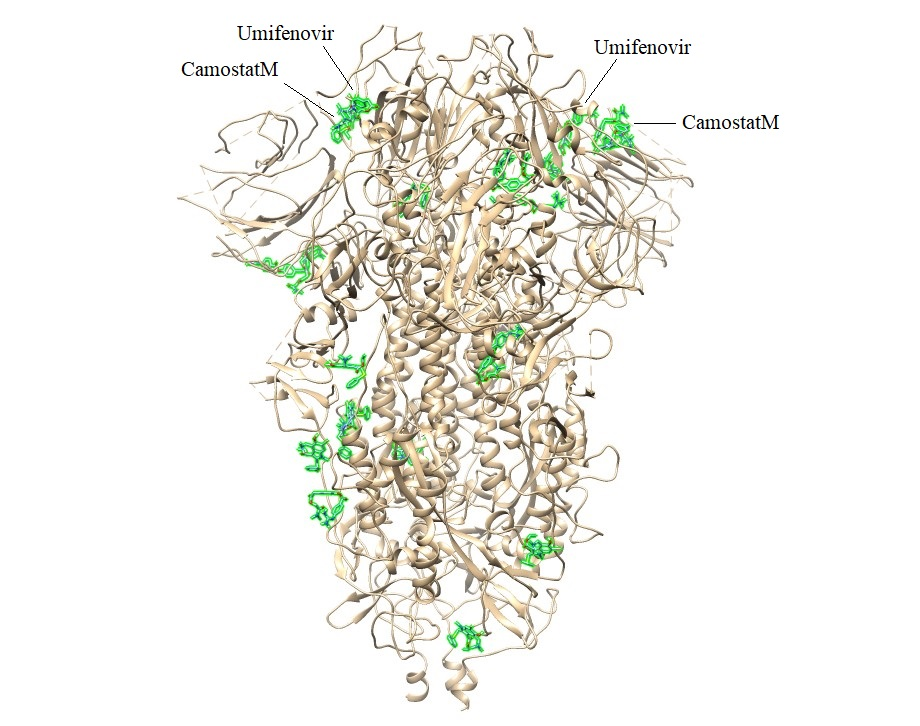  6VXX-Camostat Mesylate+Umifenovir  A2 |
| --- | --- |
| 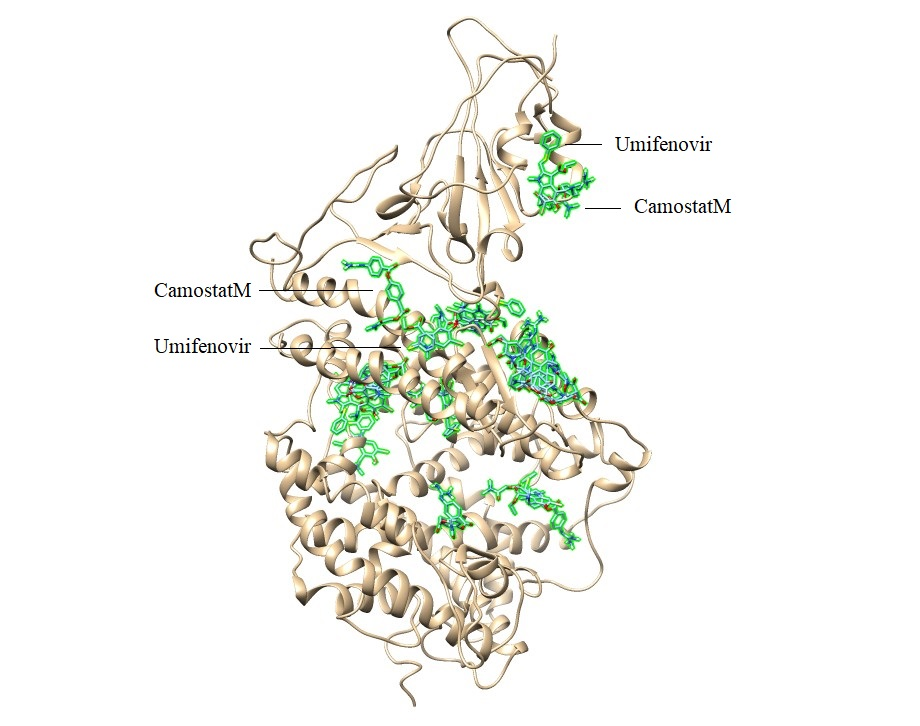  6M0J-Camostat Mesylate+Umifenovir  A3 | 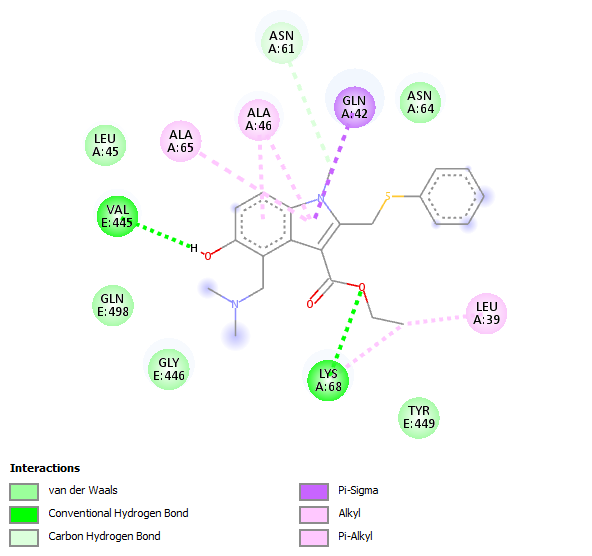  2D map of umifenovir at interface of 6M0J-Camostat Mesylate+Umifenovir complex |
| 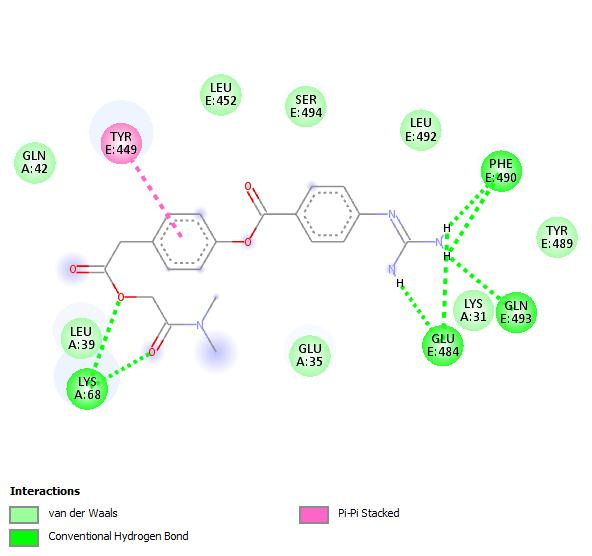  2D map of camostatM at interface of 6M0J-Camostat Mesylate+Umifenovir complex |  |
| 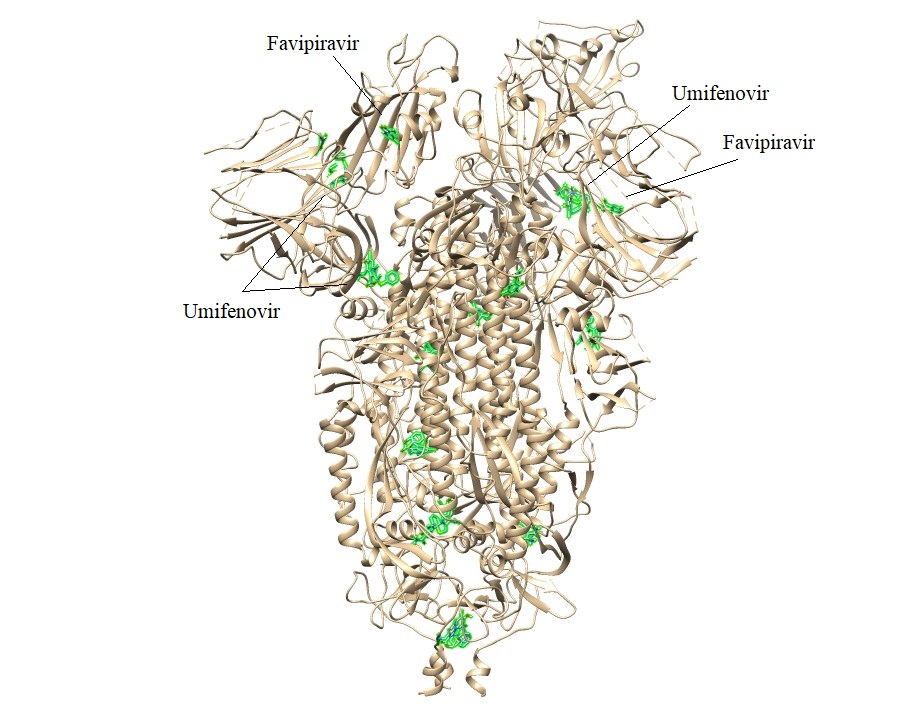  6VYB-Favipiravir+Umifenovir  B1 | 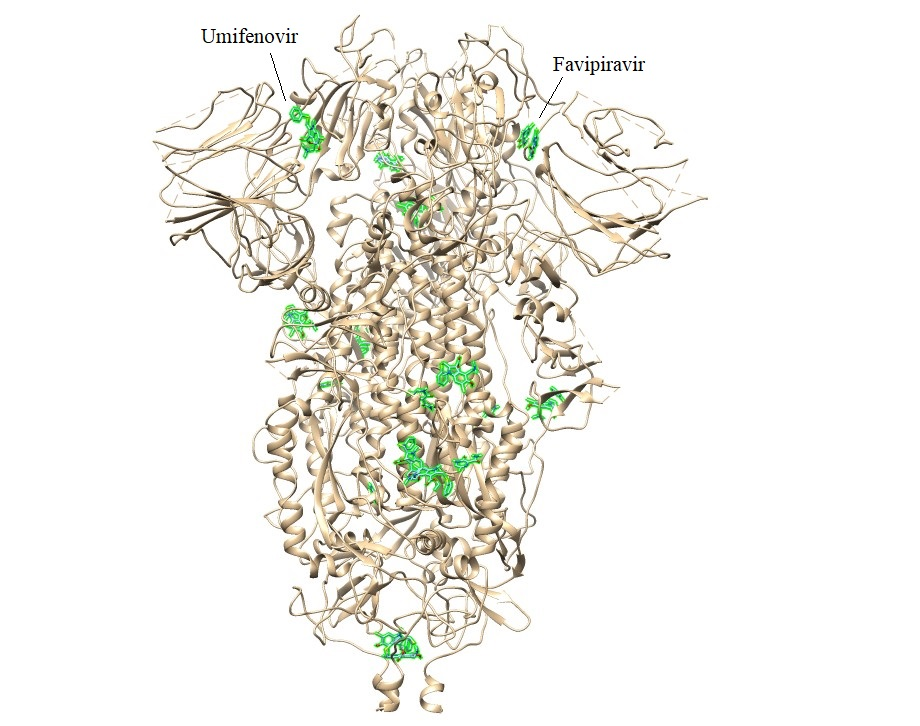  6VXX-Favipiravir+Umifenovir  B2 |
| 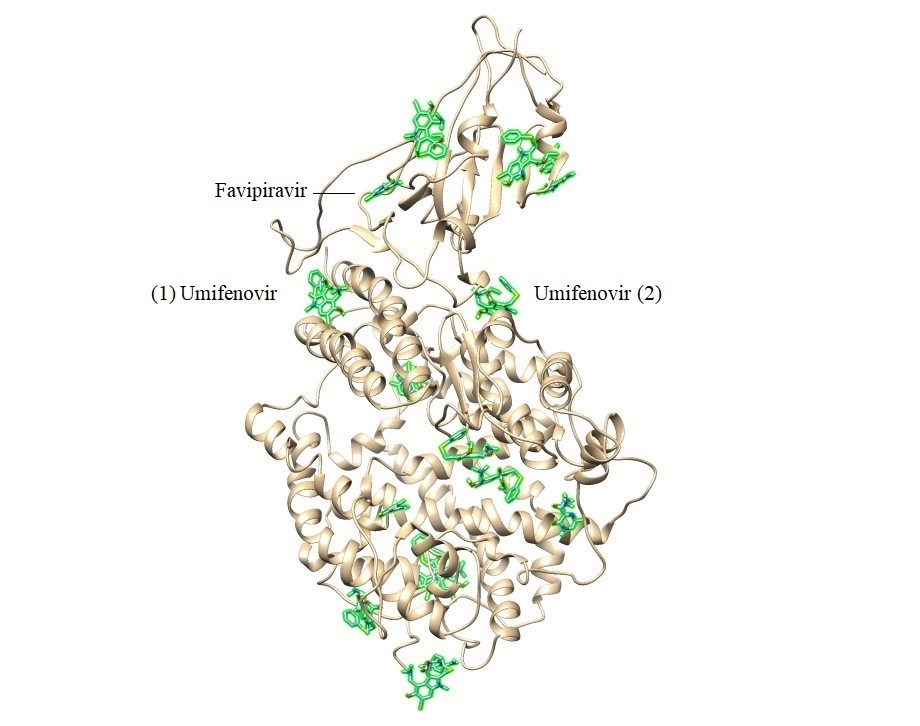  6M0J-Favipiravir+Umifenovir  B3 | 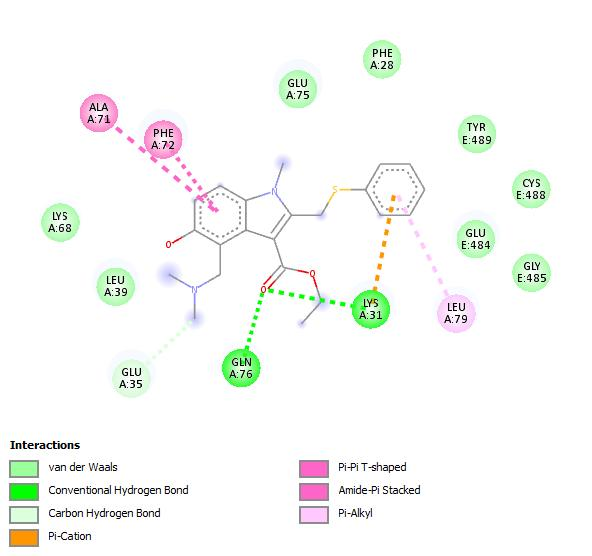  (1) 2D map of umifenovir at interface of 6M0J-Favipiravir +Umifenovir complex |
| 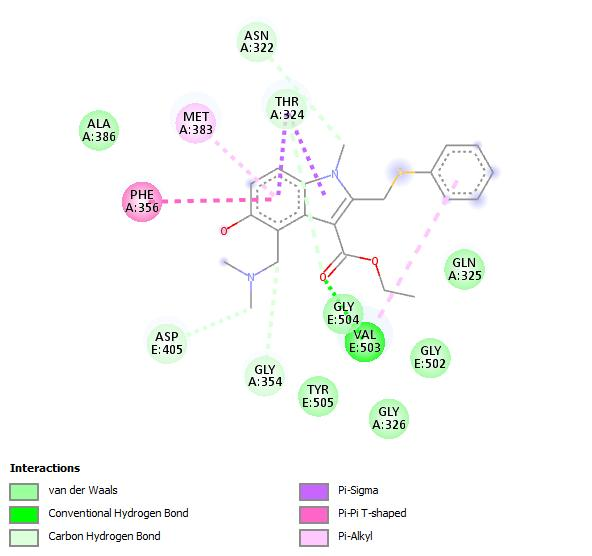  (2) 2D map of umifenovir at interface of 6M0J-Favipiravir +Umifenovir complex | 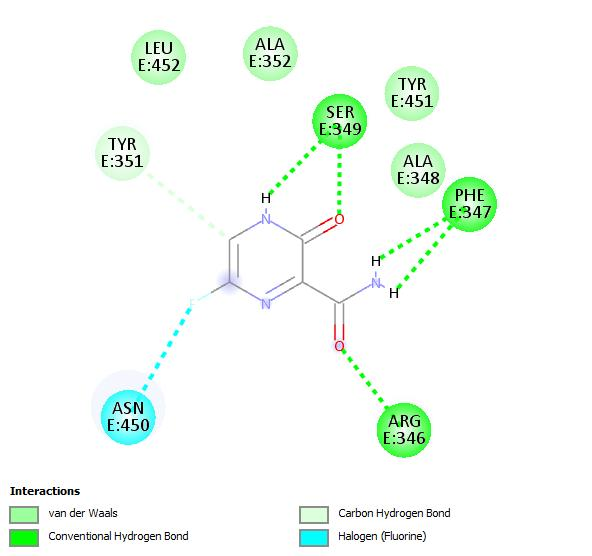  2D map of favipiravir at interface of 6M0J-Favipiravir +Umifenovir complex |
| **Figure S6 –** 3D shots of A) camostatM+umifenovir-6VYB (A1), 6VXX (A2) and 6M0J (A3) and B) favipiravir+umifenovir-6VYB (B1), 6VXX (B2) and 6M0J (B3) protein complexes. | |
