## Supplementary figures and images for "Favipiravir, umifenovir and camostat mesylate: a comparative study against SARS-CoV-2"

### camo_1r42.jpg

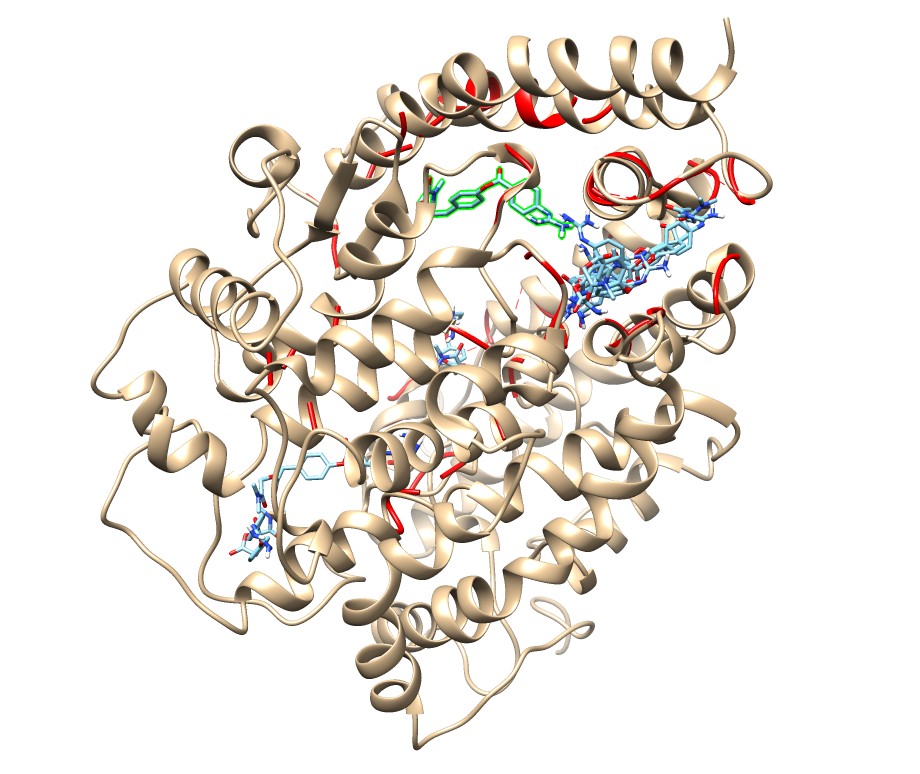

### camo_5x29.jpg

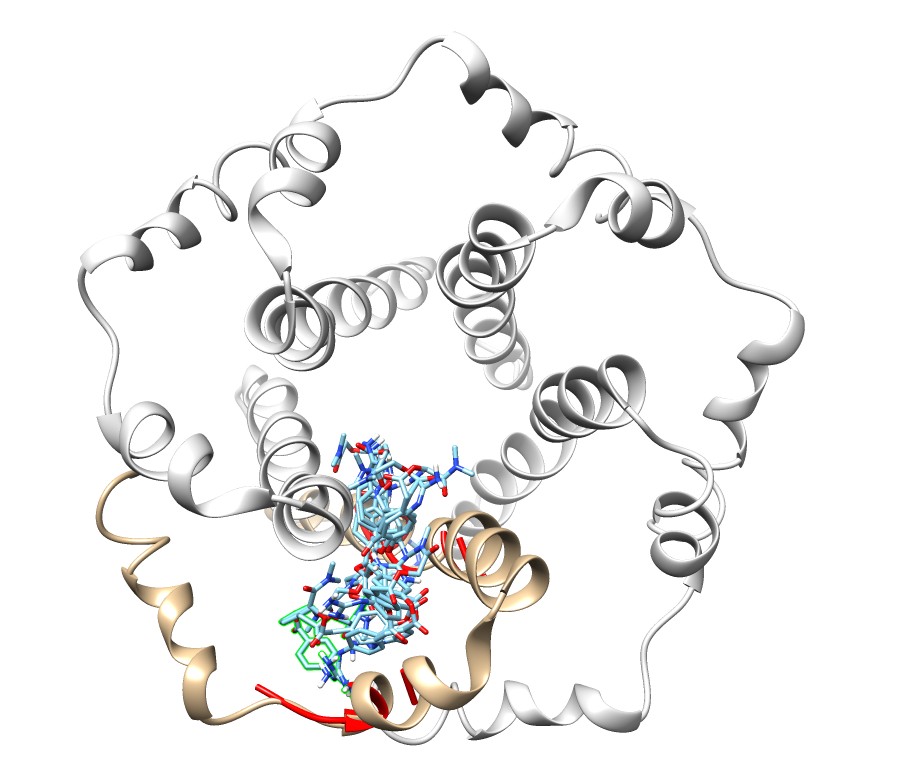

### camo_6lu7.jpg

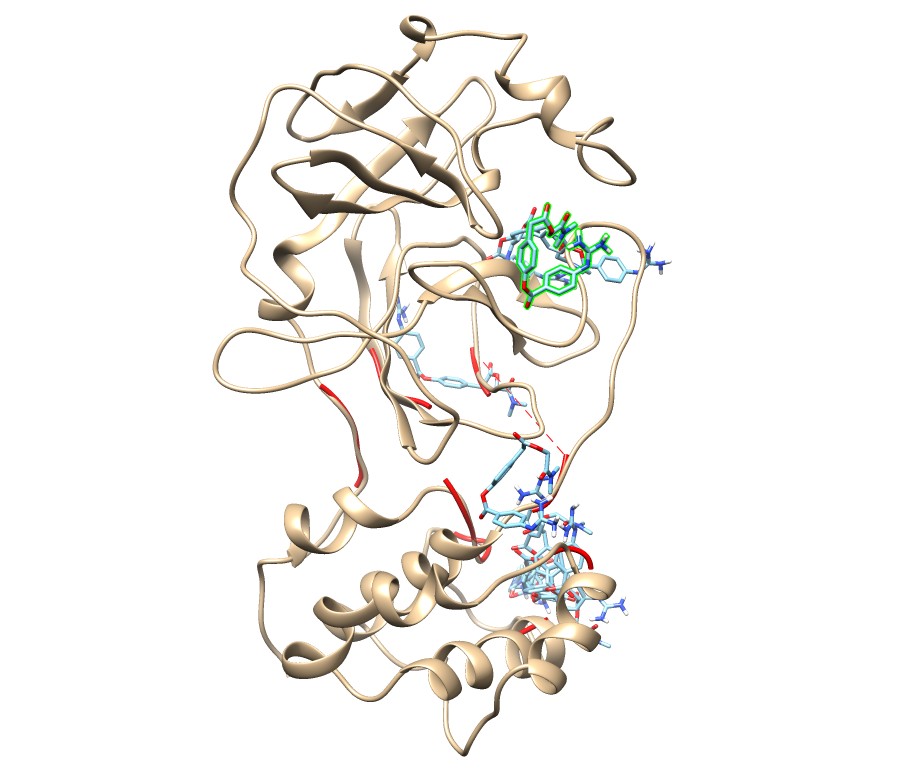

### camo_6lxt.jpg

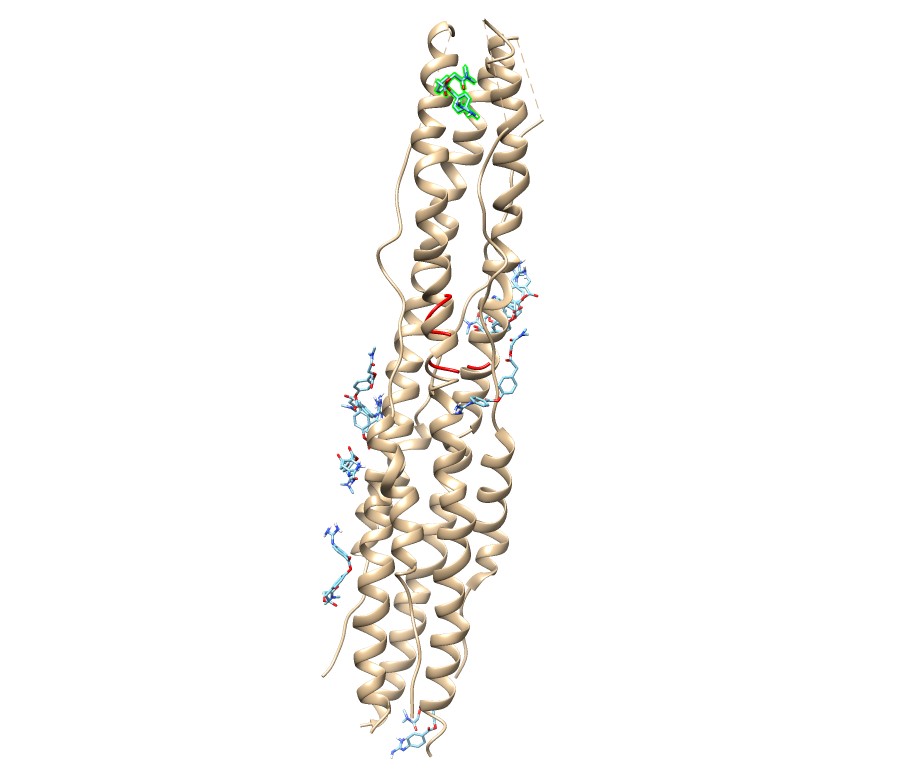

### camo_6m0j.jpg

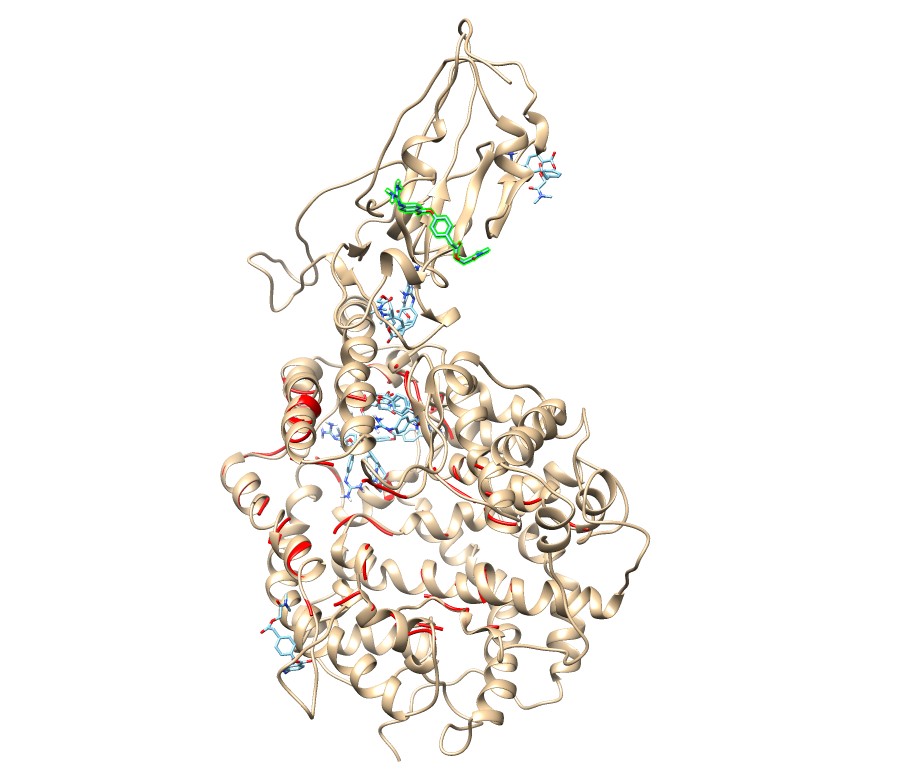

### camo_6m03.jpg

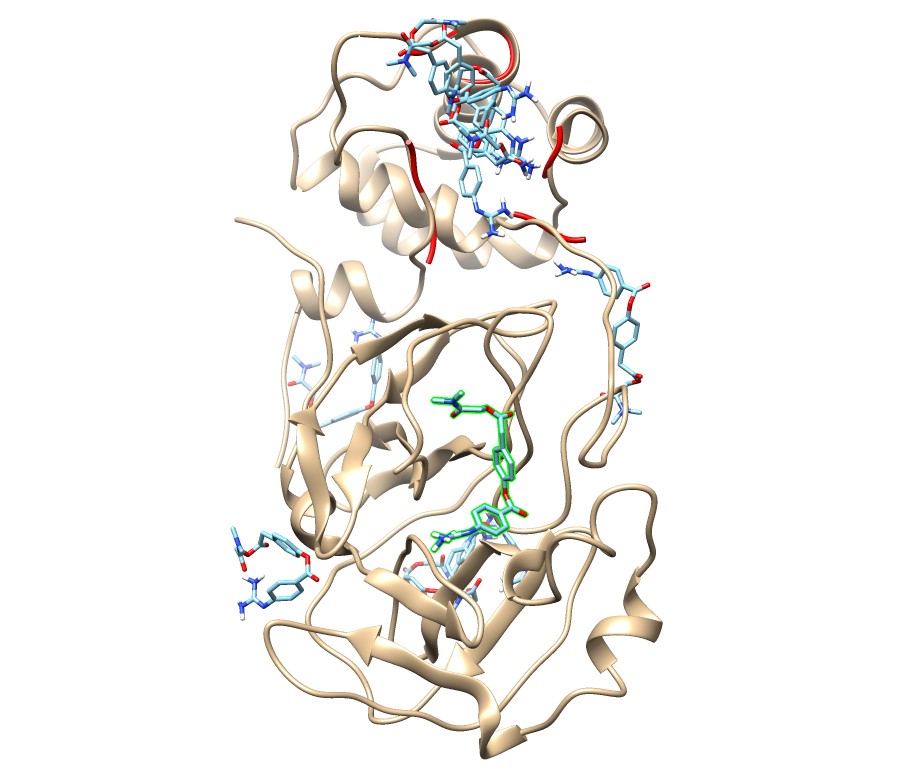

### camo_6m71.jpg

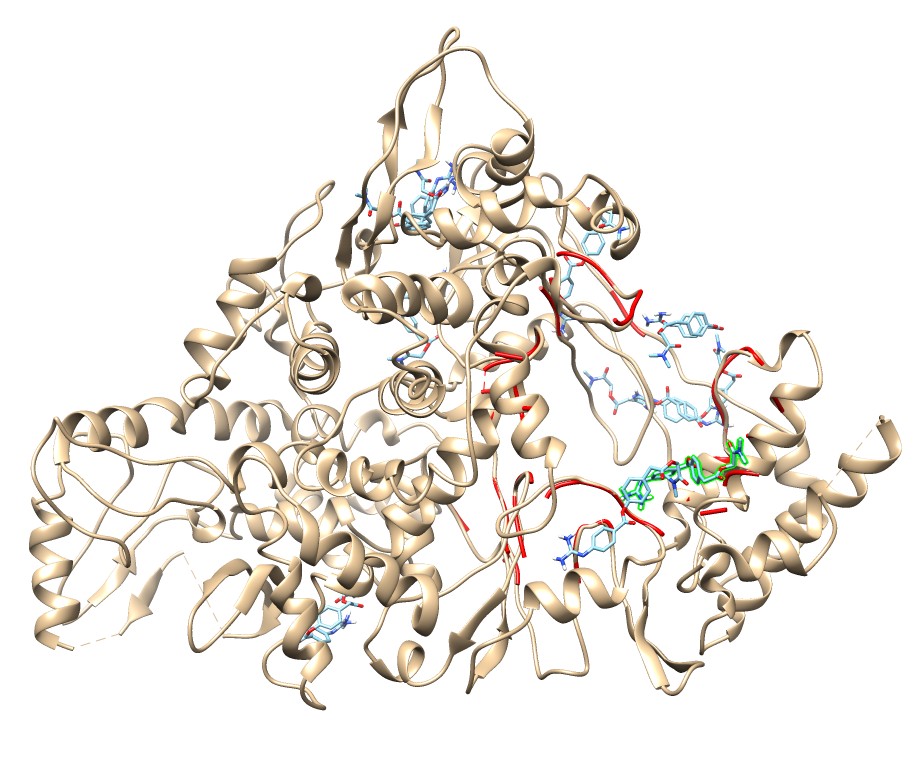

### camo_6vww.jpg

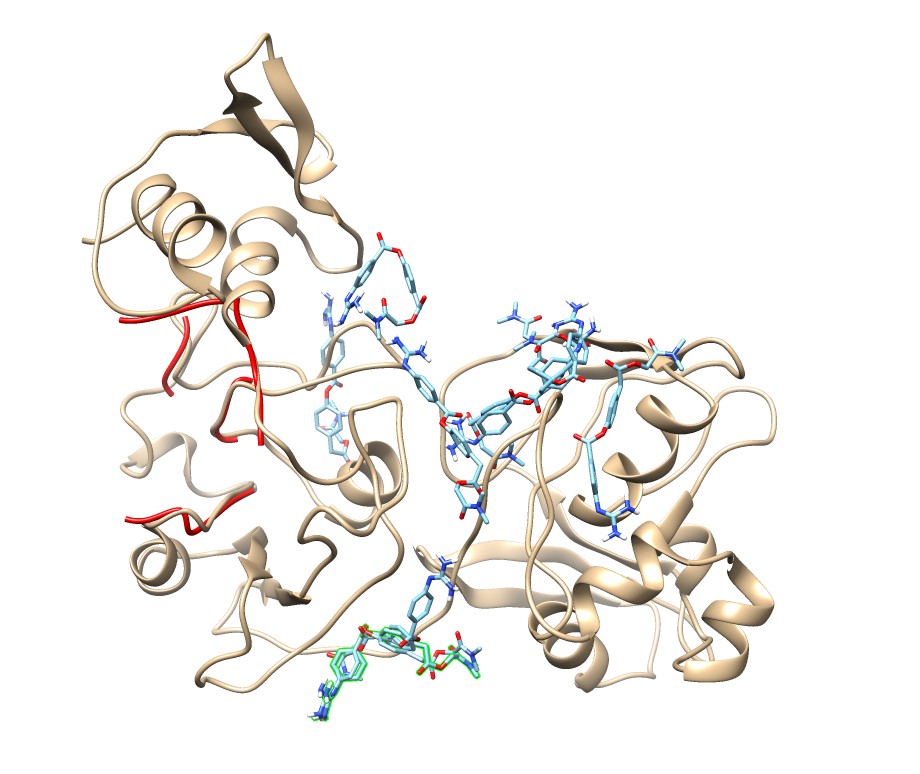

### camo_6vxx.jpg

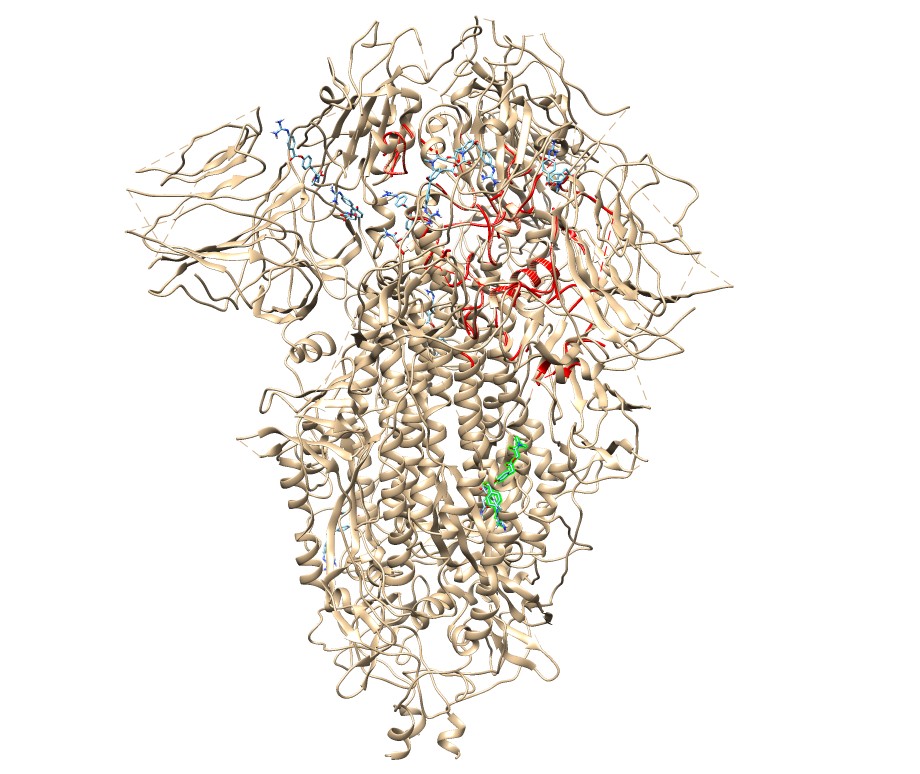

### camo_6vyb.jpg

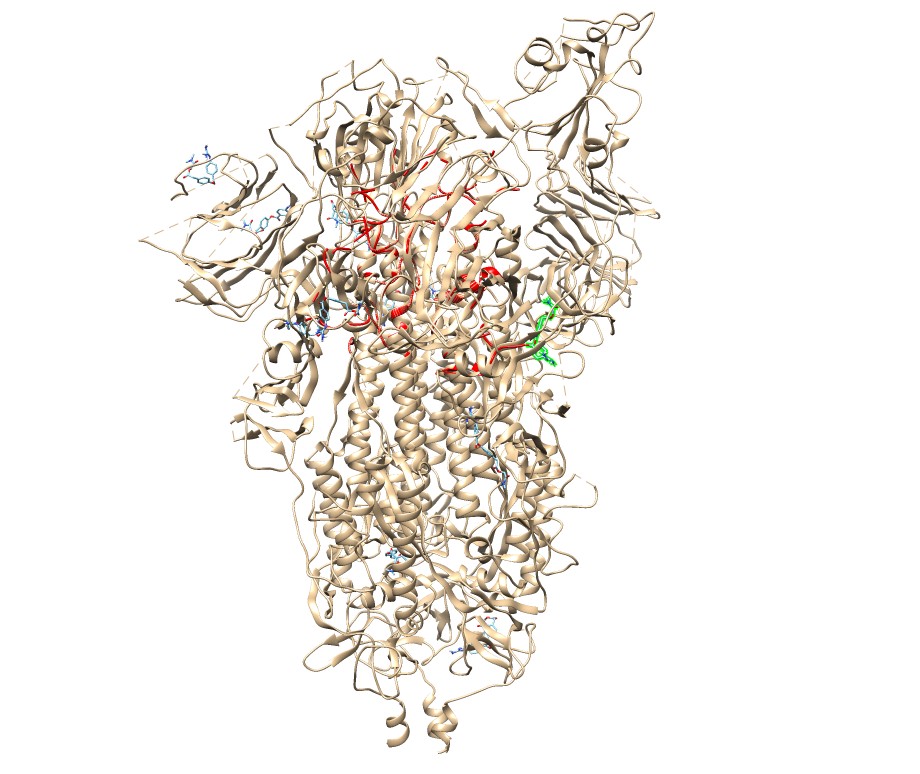

### camo_6vyo.jpg

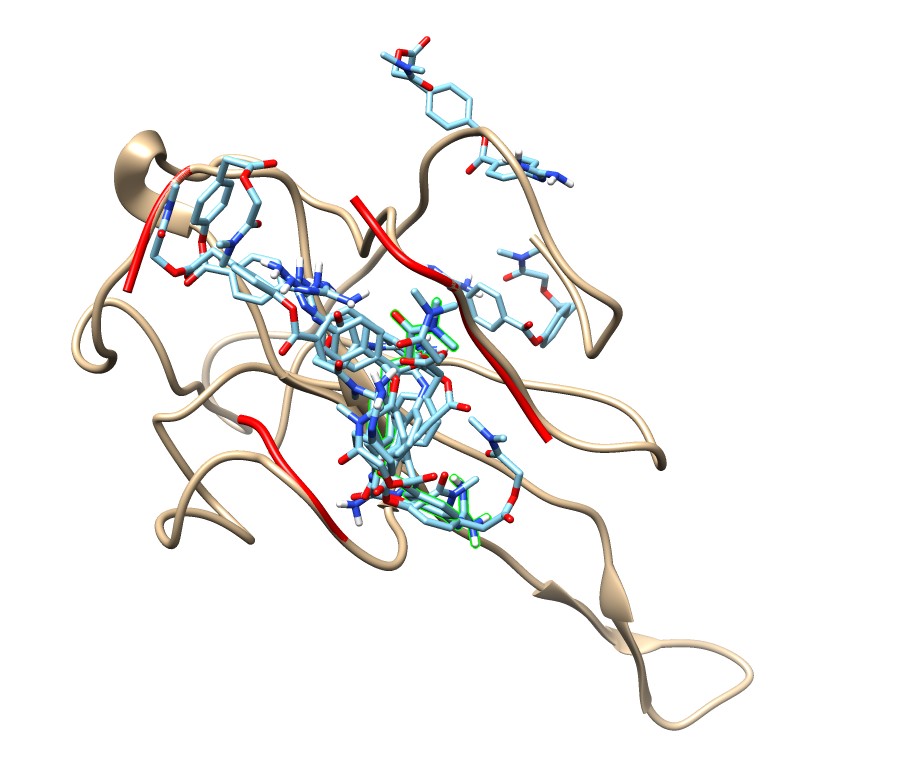

### camo_6y84.jpg

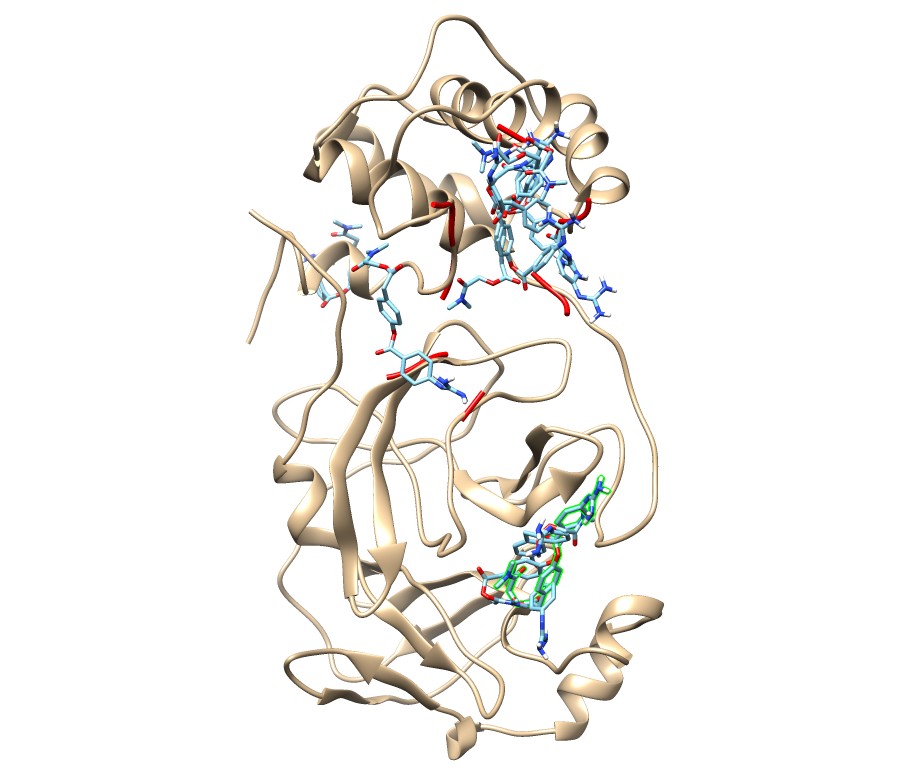

### camofavi_1r42.jpg

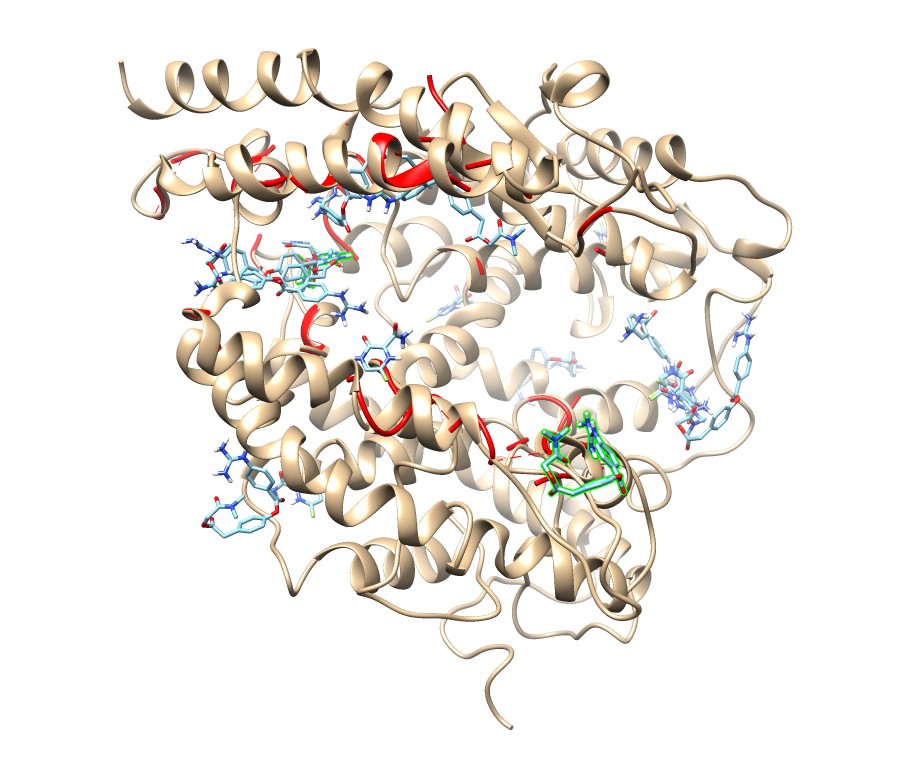

### camofavi_5x29.jpg

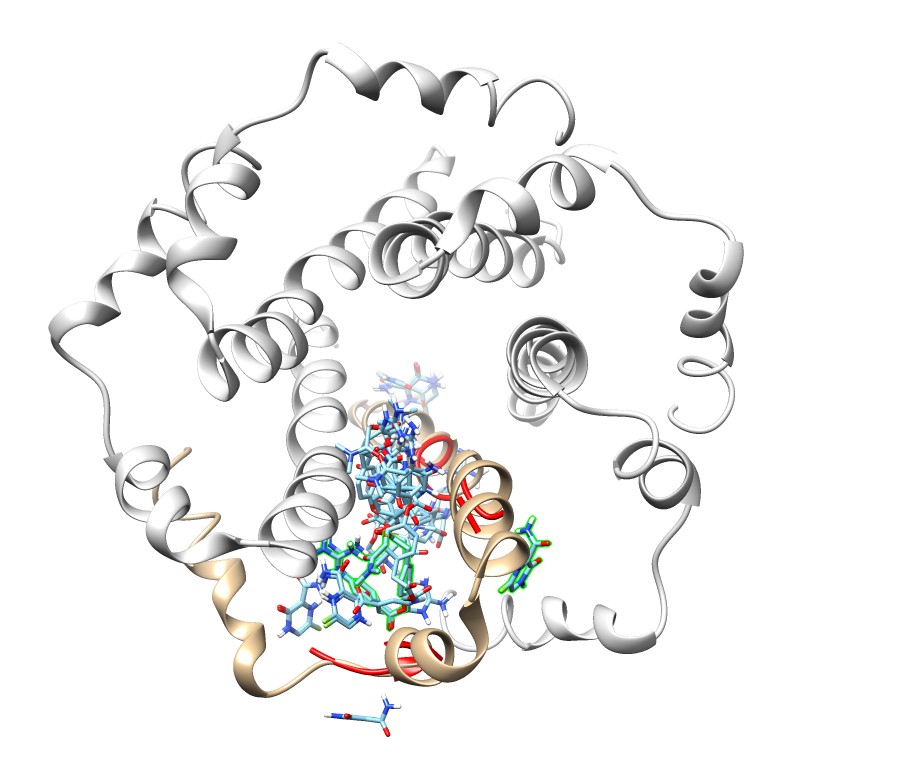

### camofavi_6lu7.jpg

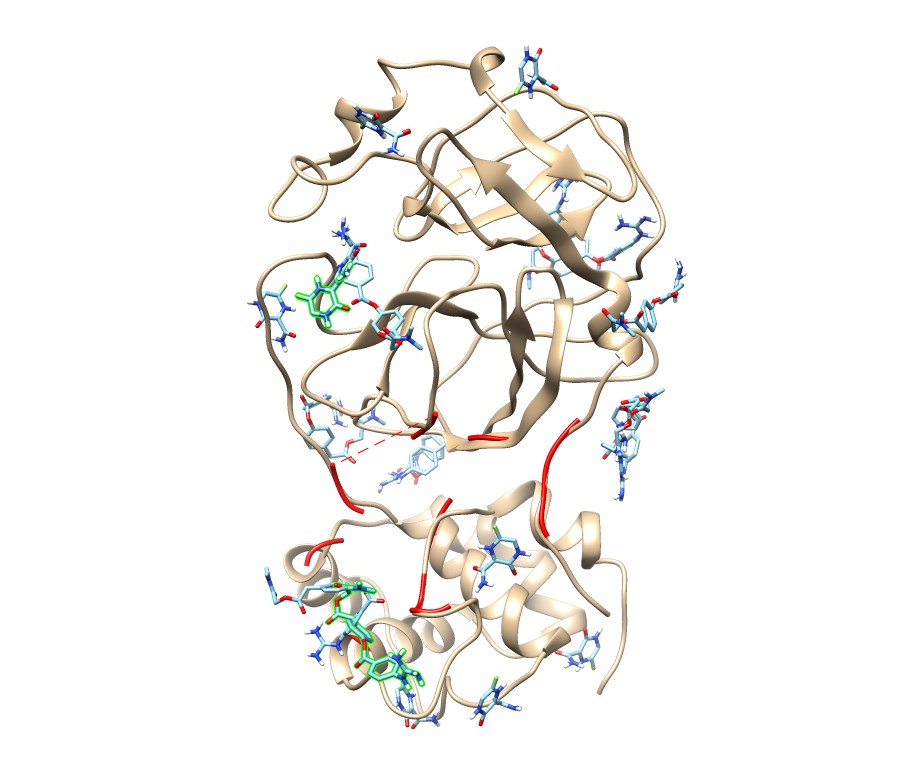

### camofavi_6lxt.jpg

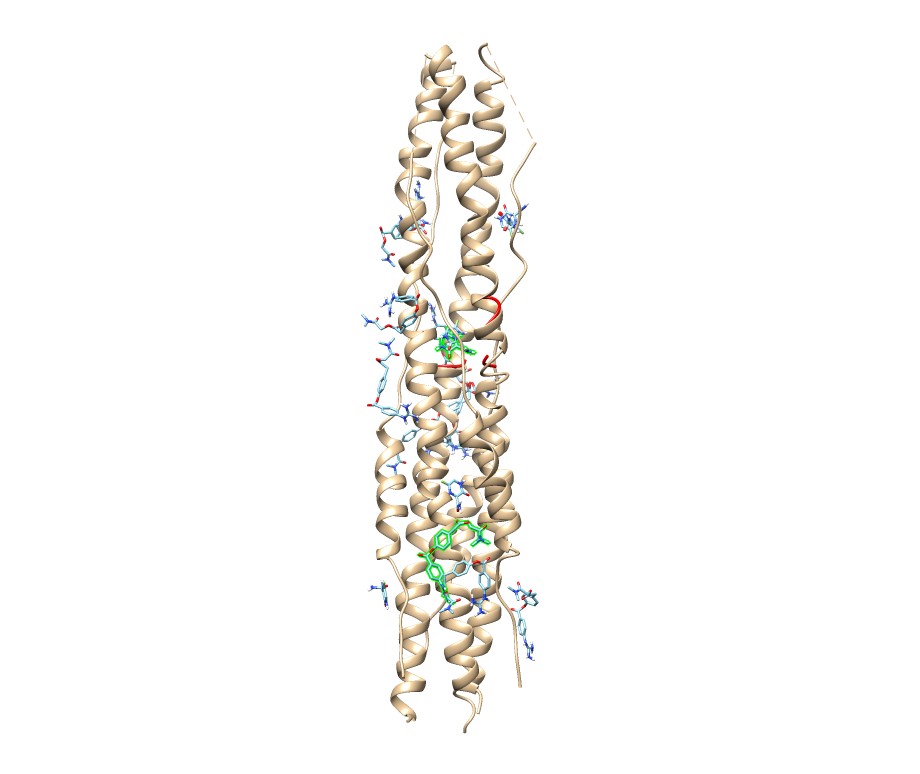

### camofavi_6m0j.jpg

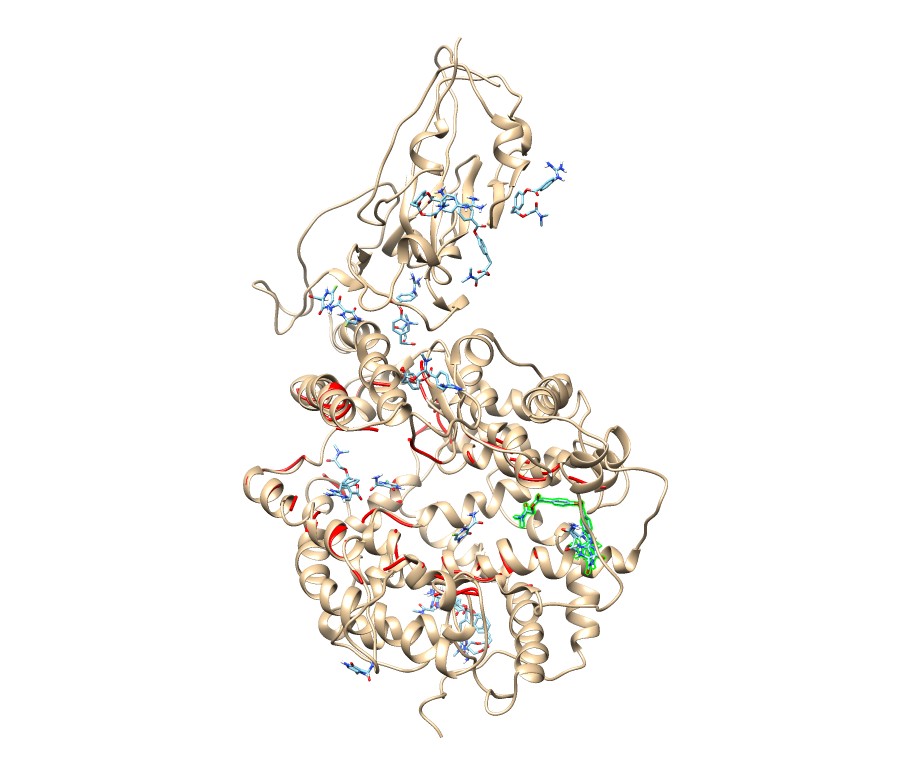

### camofavi_6m03.jpg

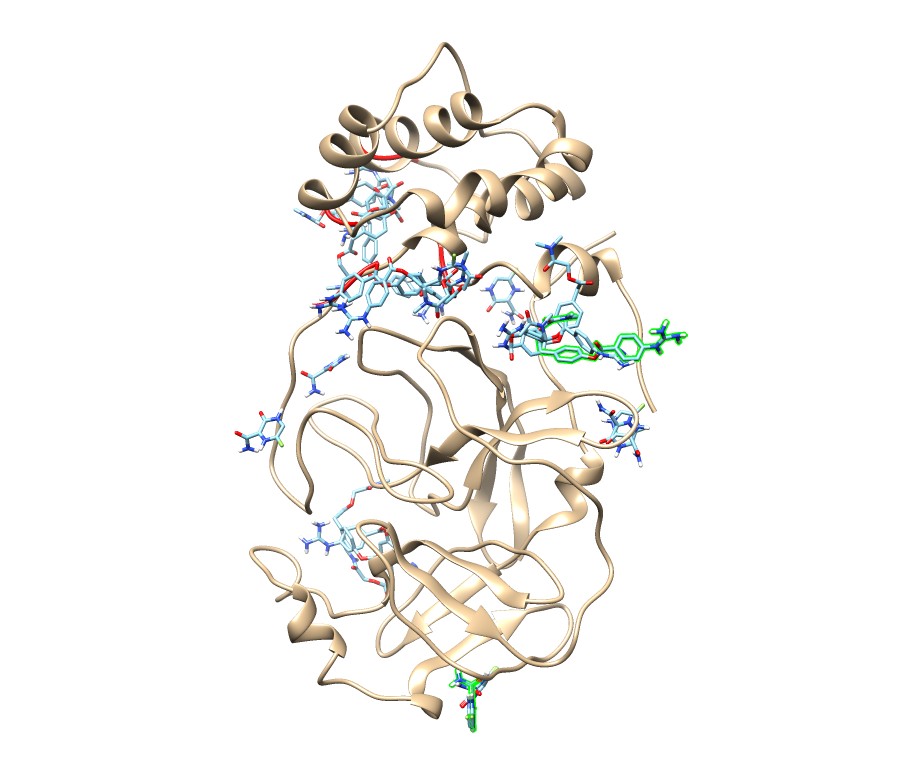

### camofavi_6m71.jpg

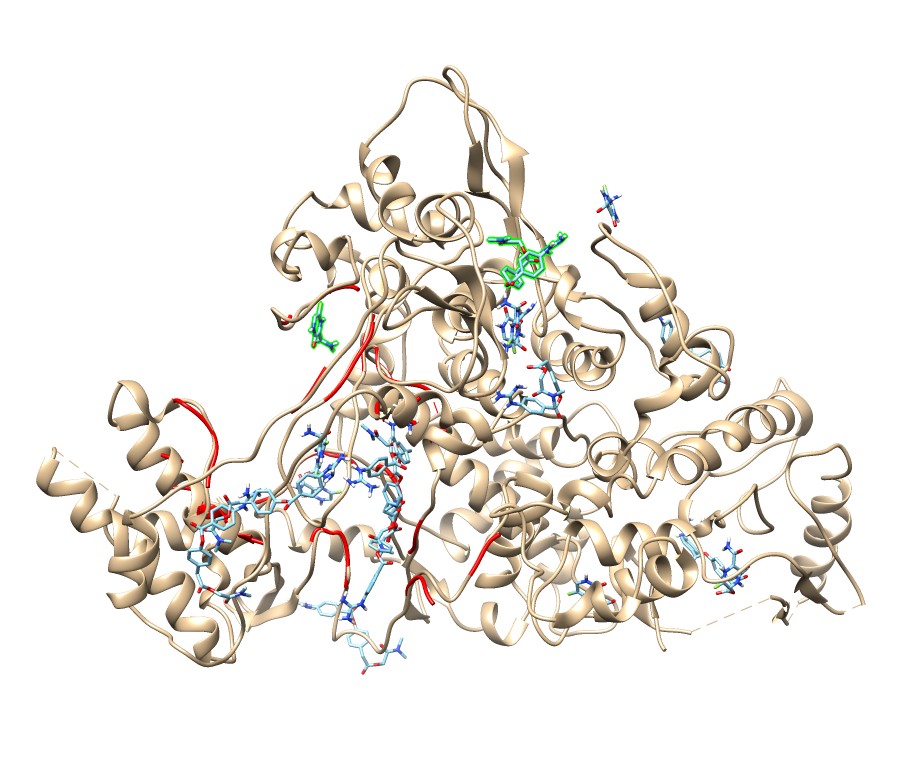

### camofavi_6vww.jpg

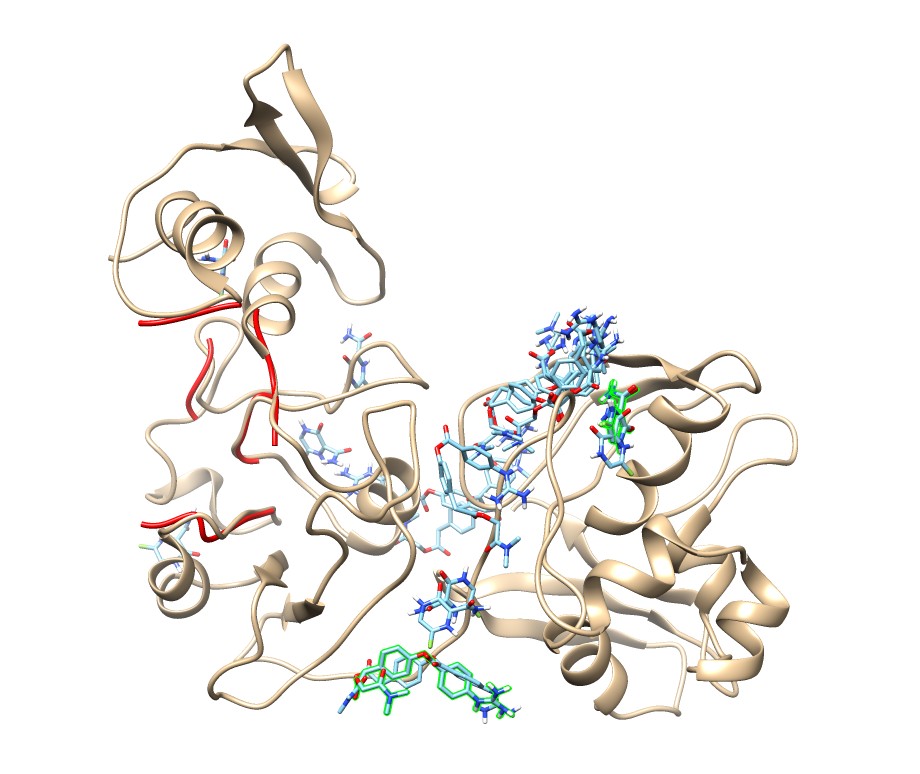

### camofavi_6vxx.jpg

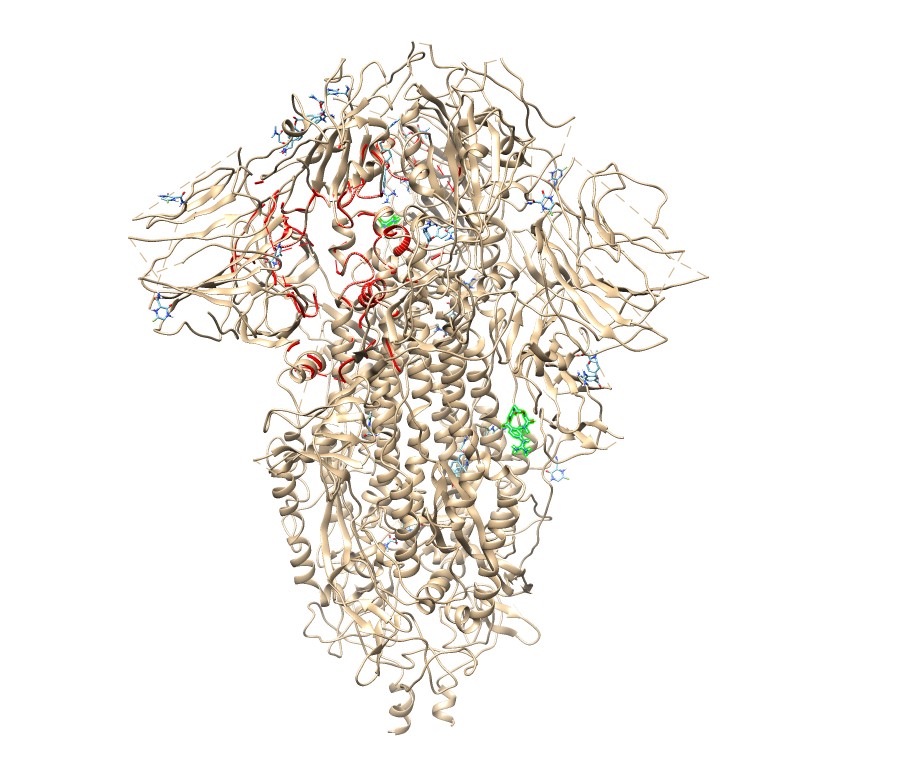

### camofavi_6vyb.jpg

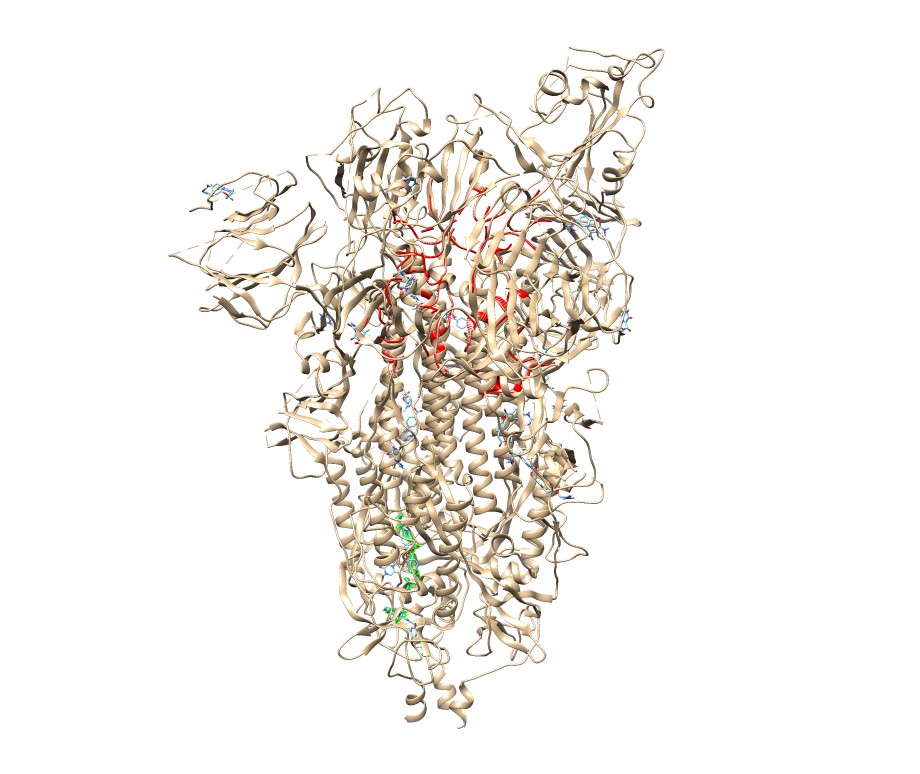

### camofavi_6vyo.jpg

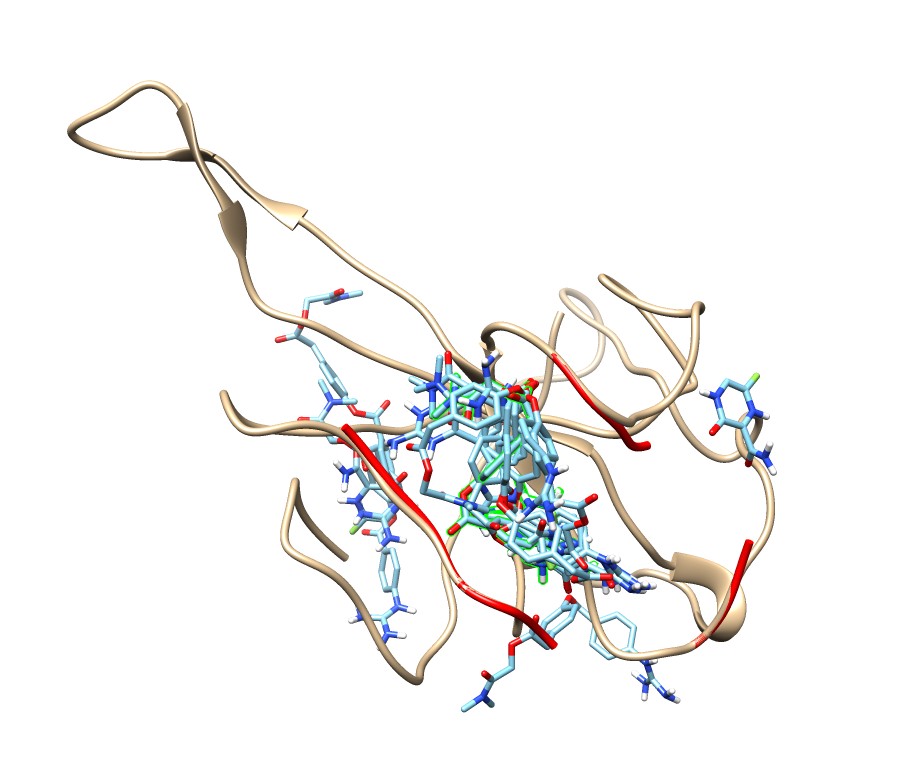

### camofavi_6y84.jpg

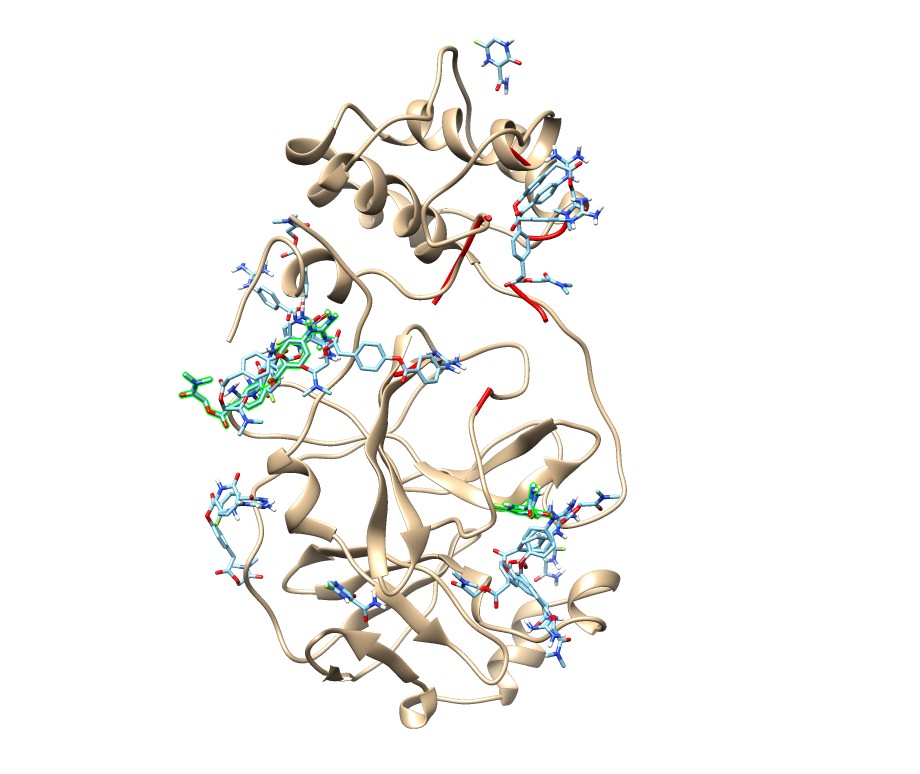

### camumif_1r42.jpg

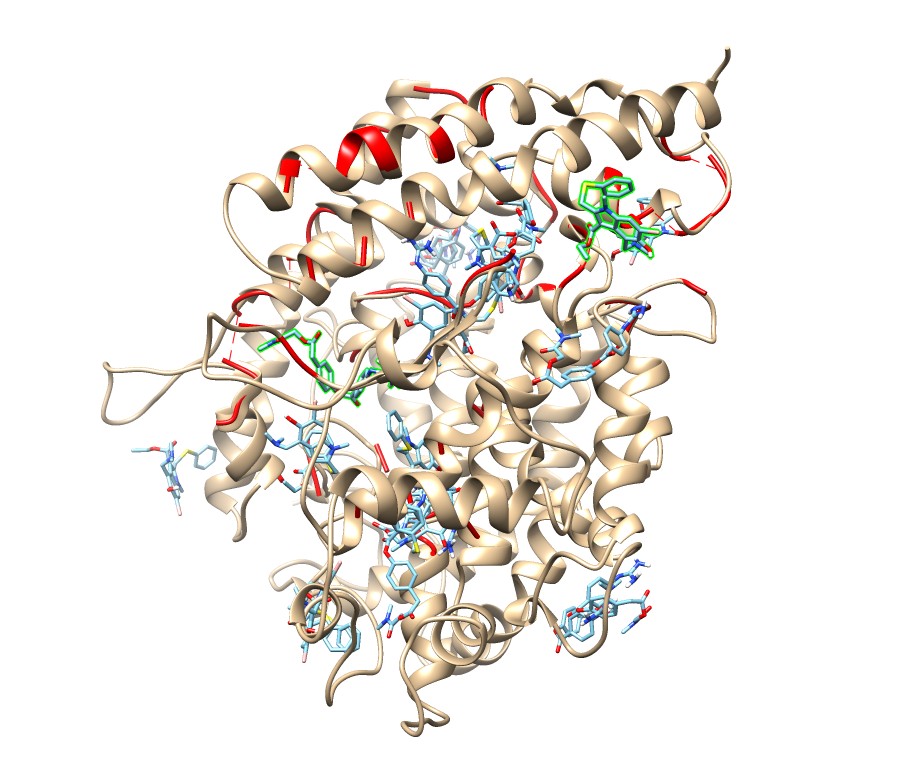

### camumif_5x29.jpg

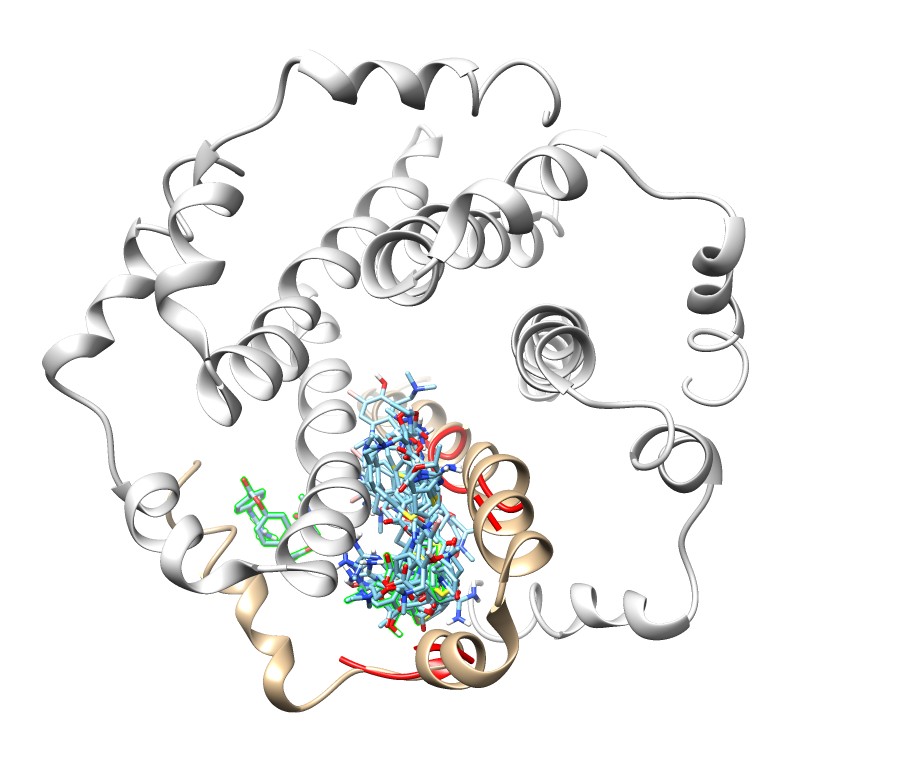

### camumif_6lu7.jpg

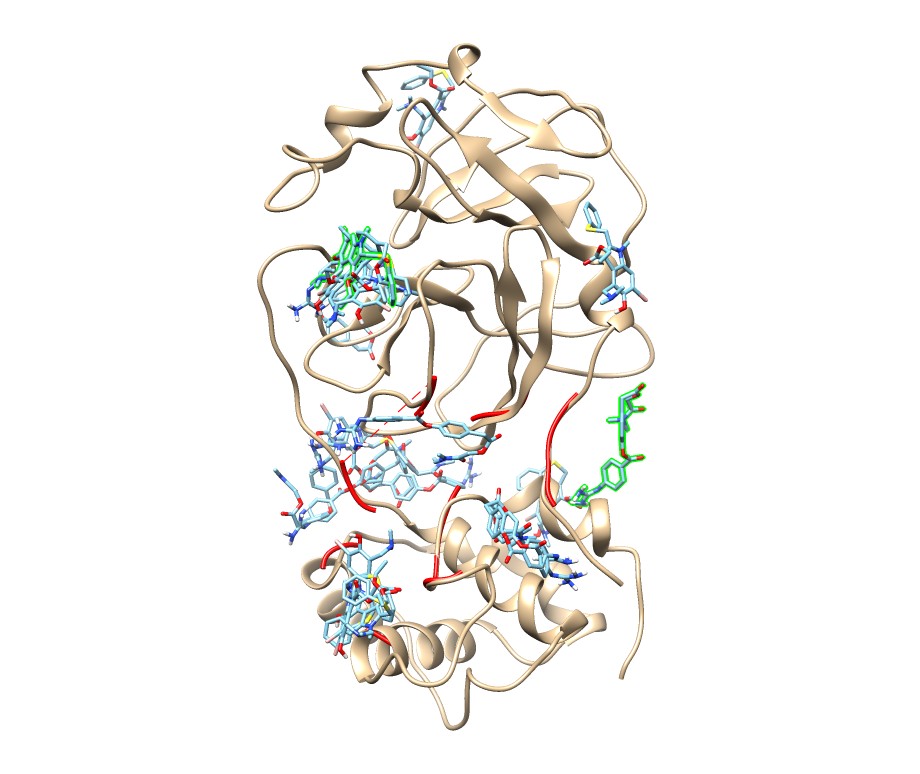

### camumif_6lxt.jpg

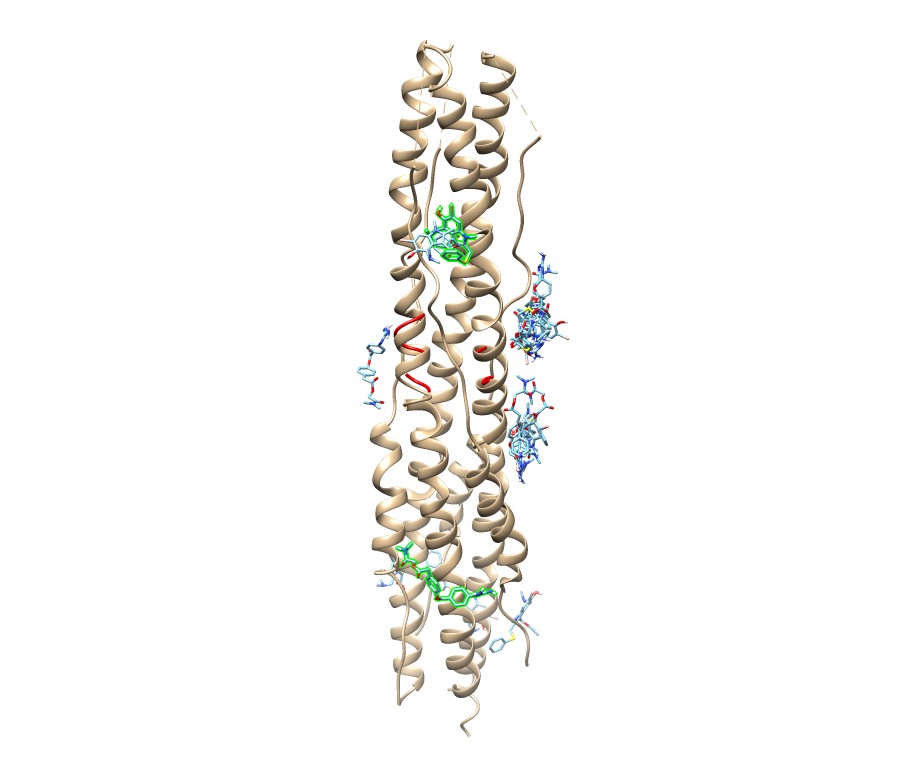

### camumif_6m0j.jpg

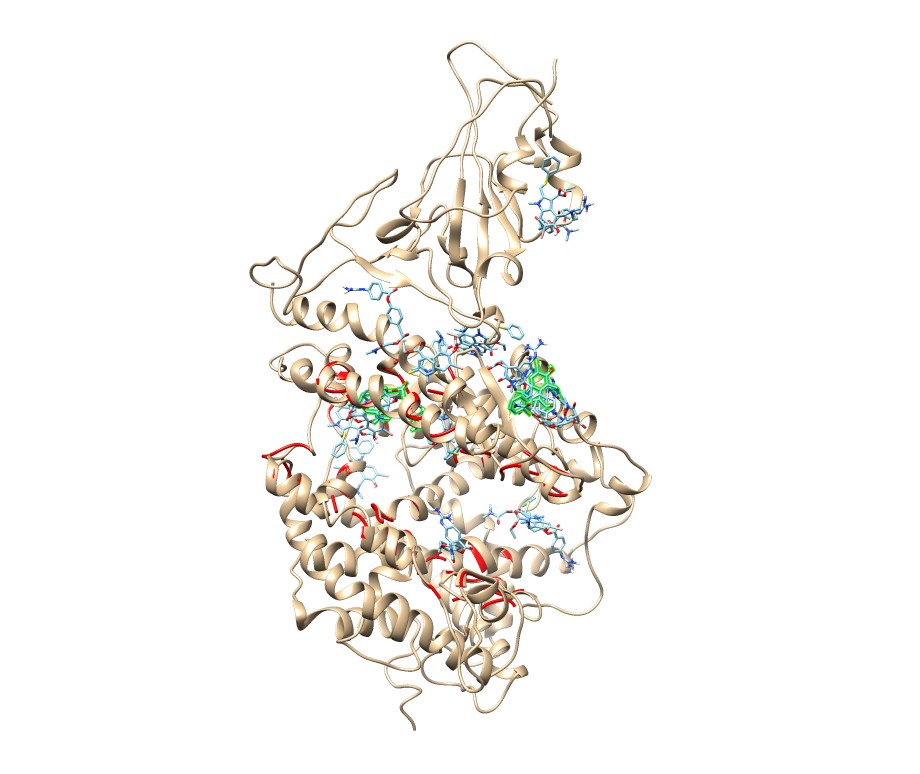

### camumif_6m03.jpg

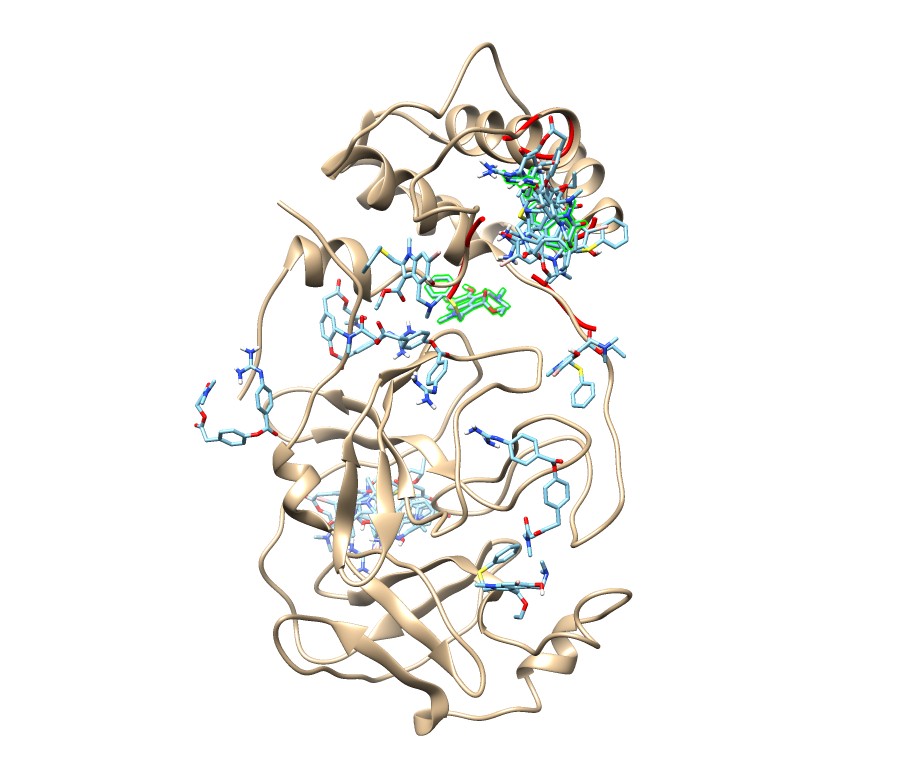

### camumif_6m71.jpg

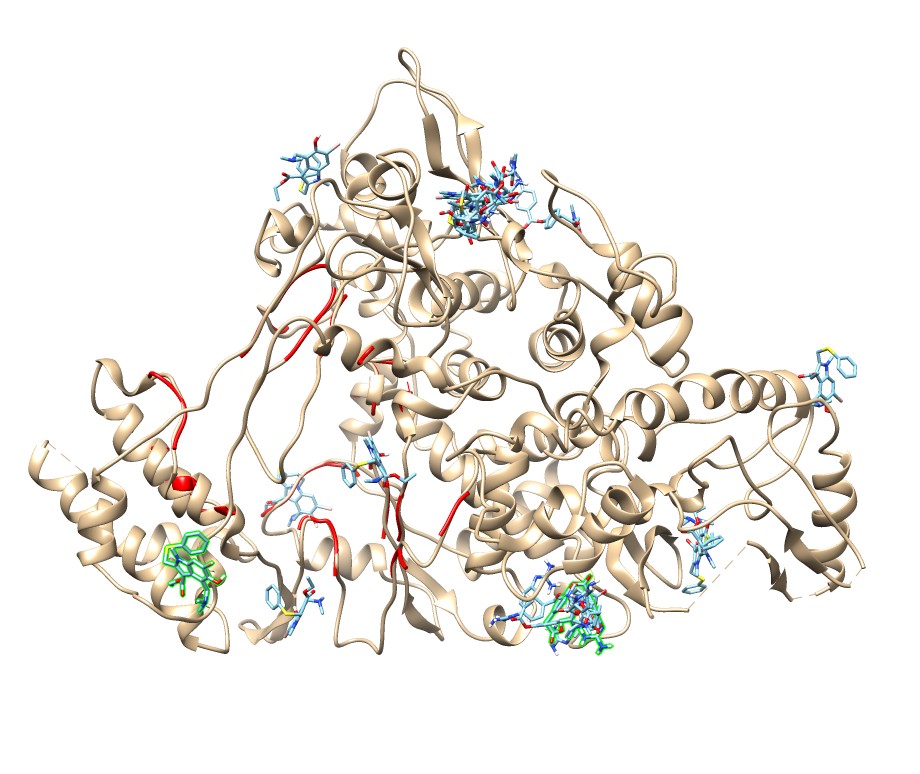

### camumif_6vww.jpg

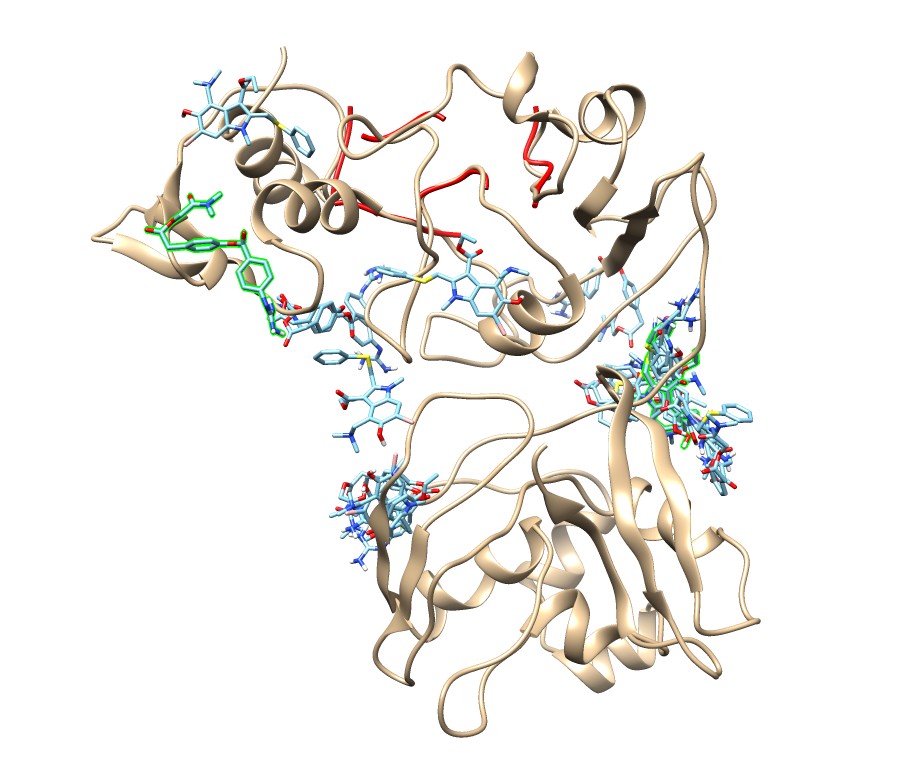

### camumif_6vxx.jpg

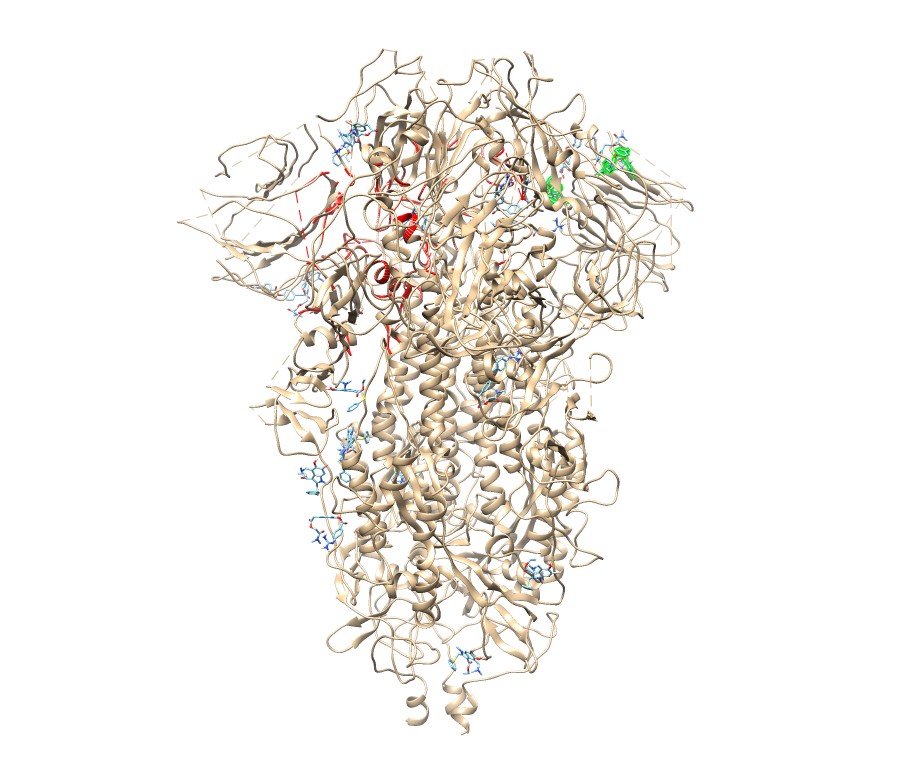

### camumif_6vyb.jpg

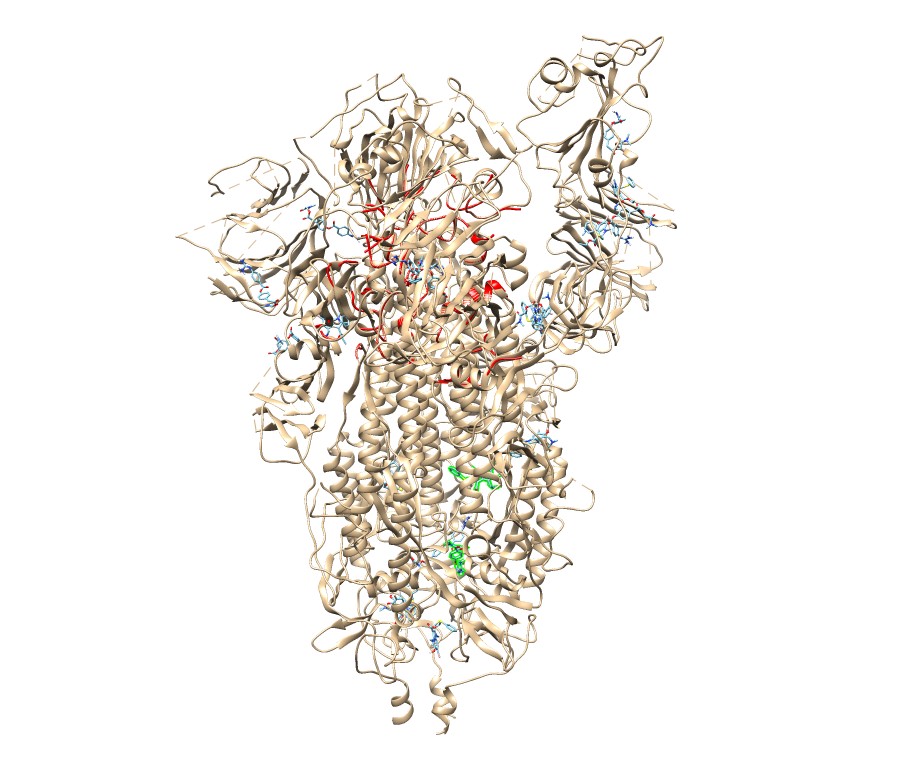

### camumif_6vyo.jpg

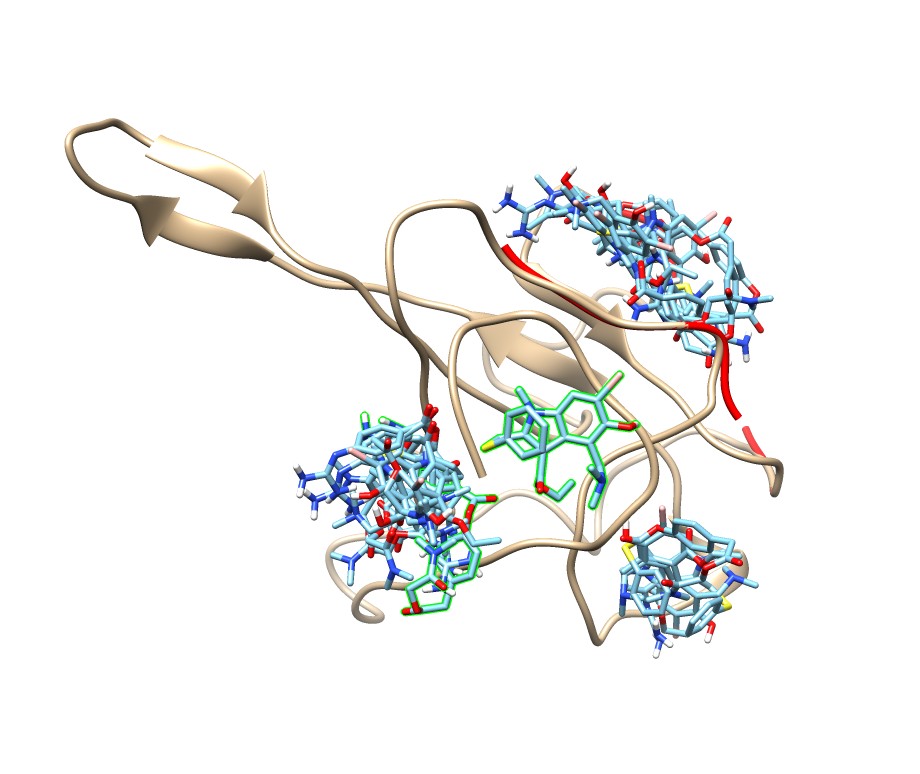

### camumif_6y84.jpg

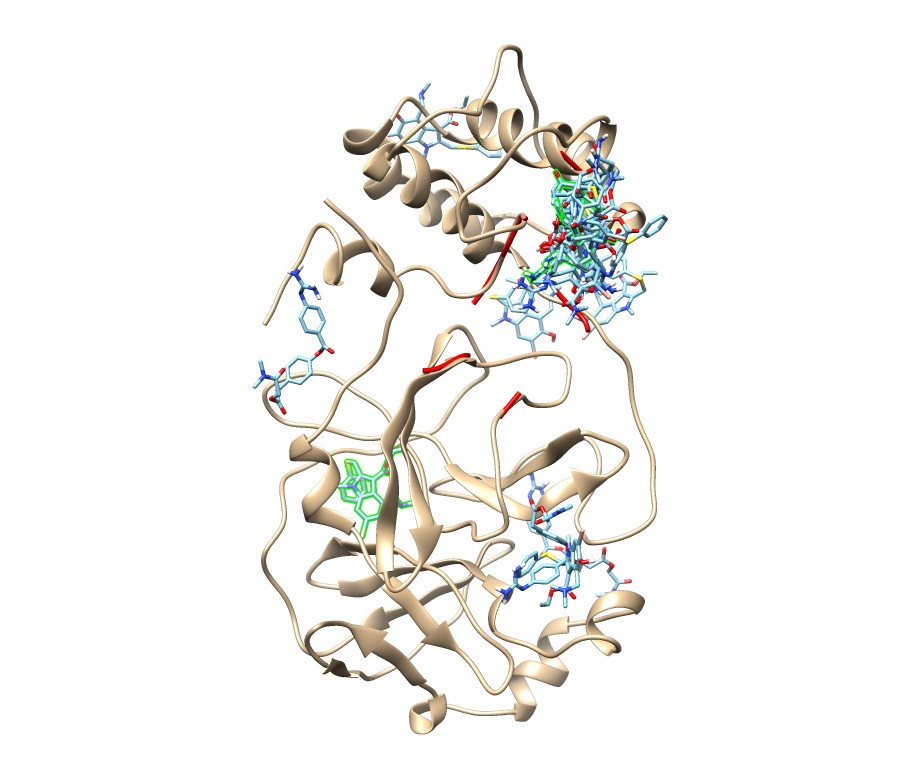

### favi_1r42.jpg

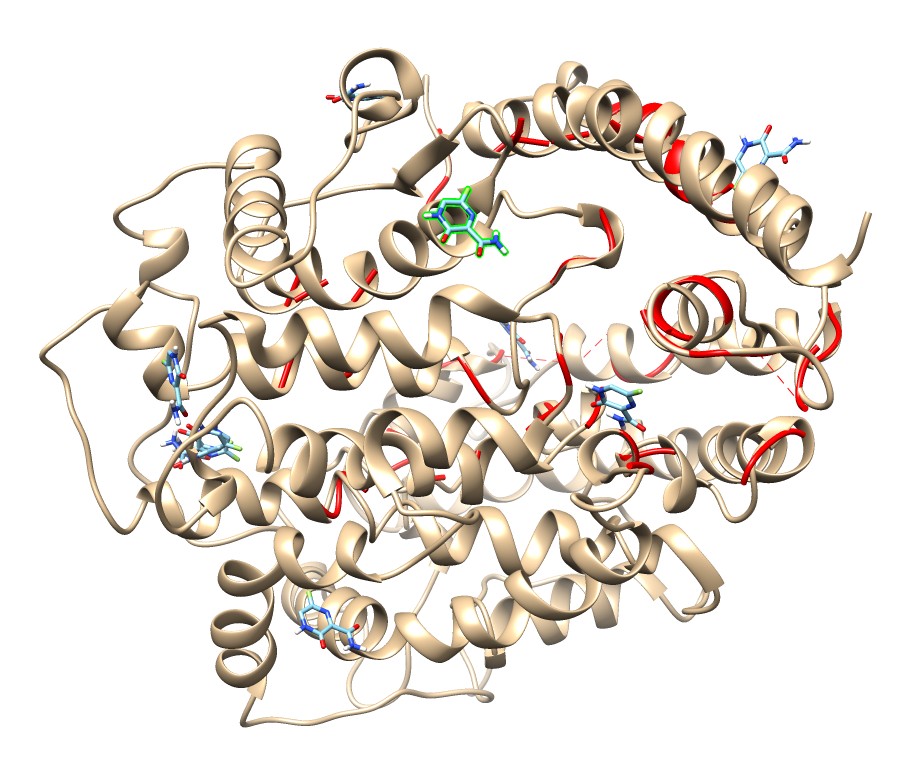

### favi_5x29.jpg

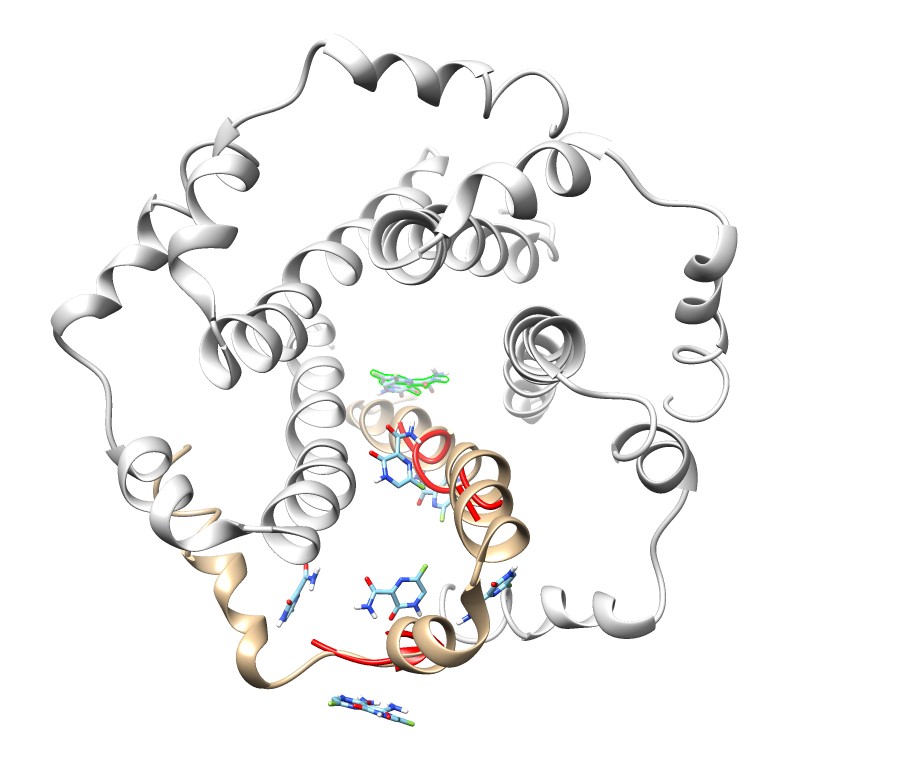

### favi_6lu7.jpg

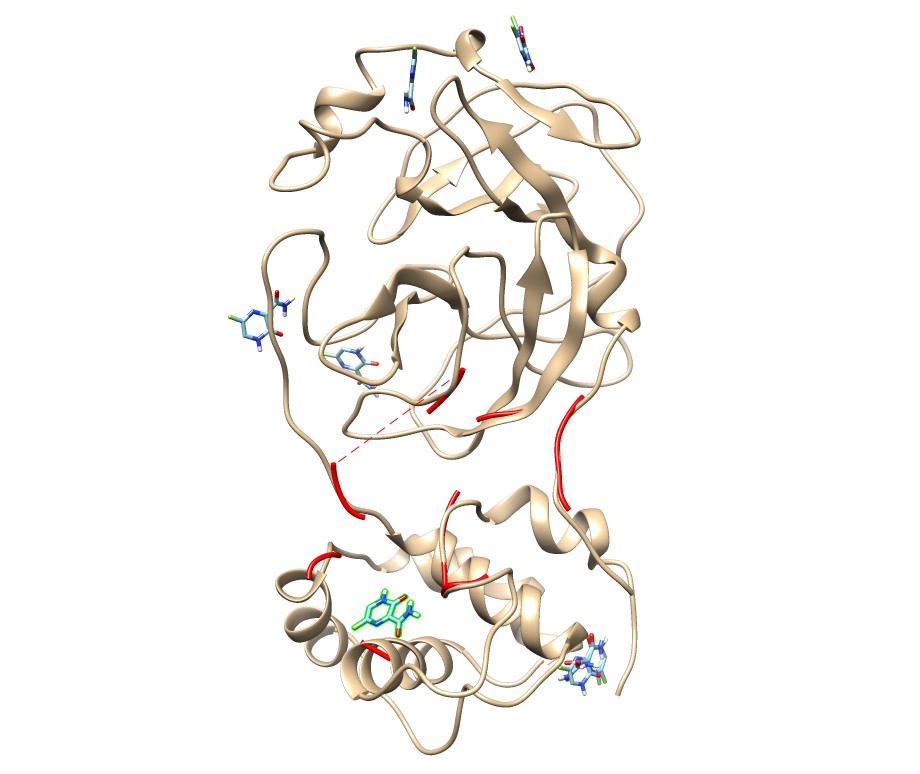

### favi_6lxt.jpg

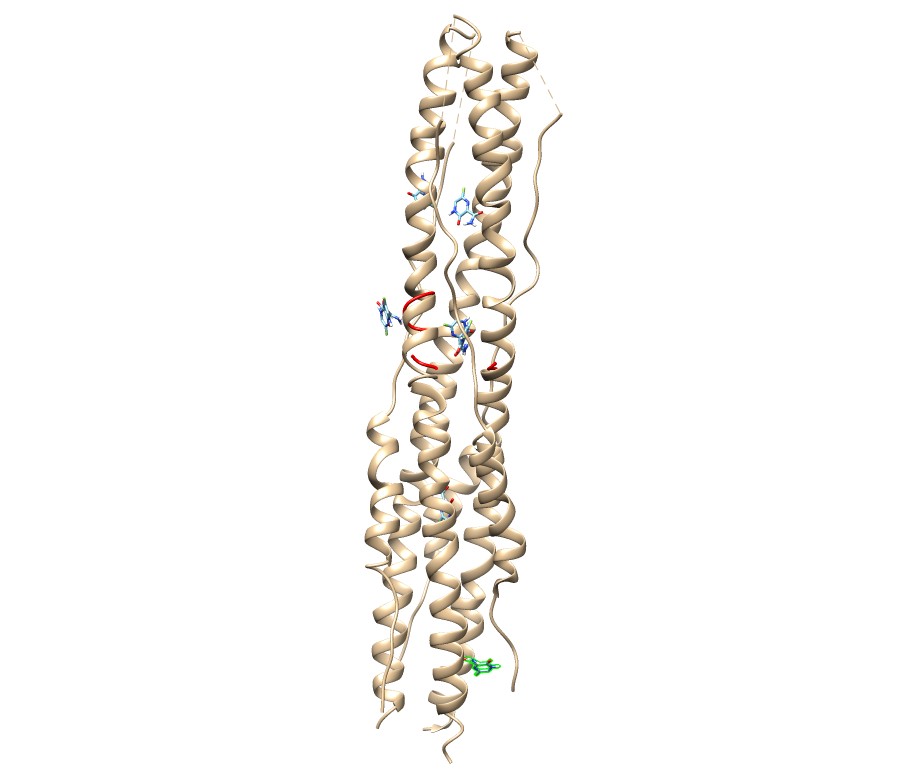

### favi_6m0j.jpg

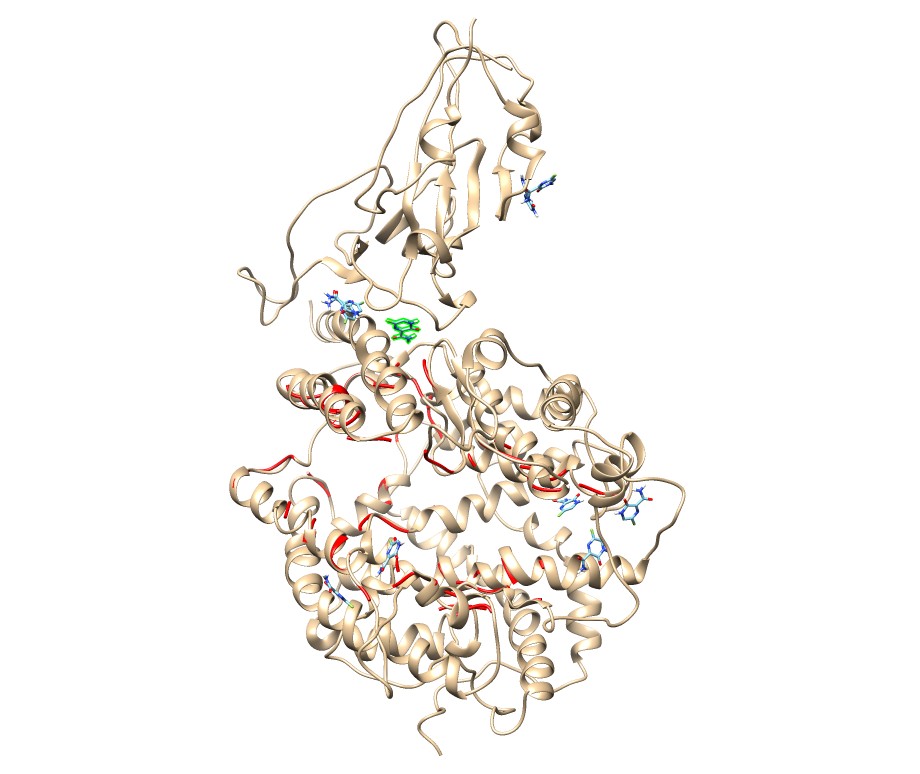

### favi_6m03.jpg

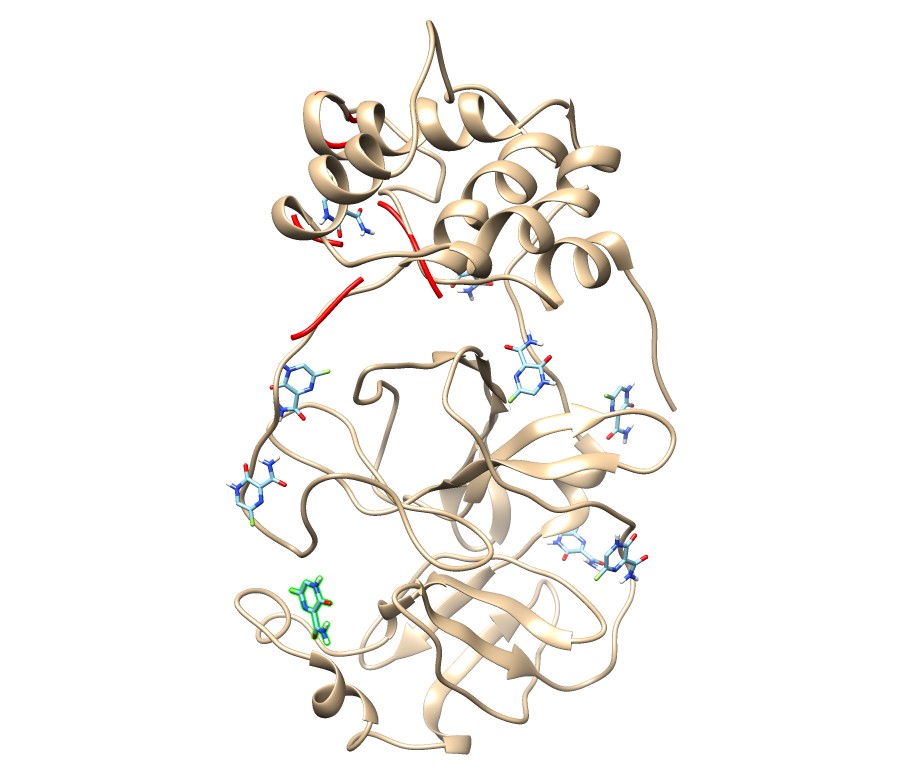

### favi_6m71.jpg

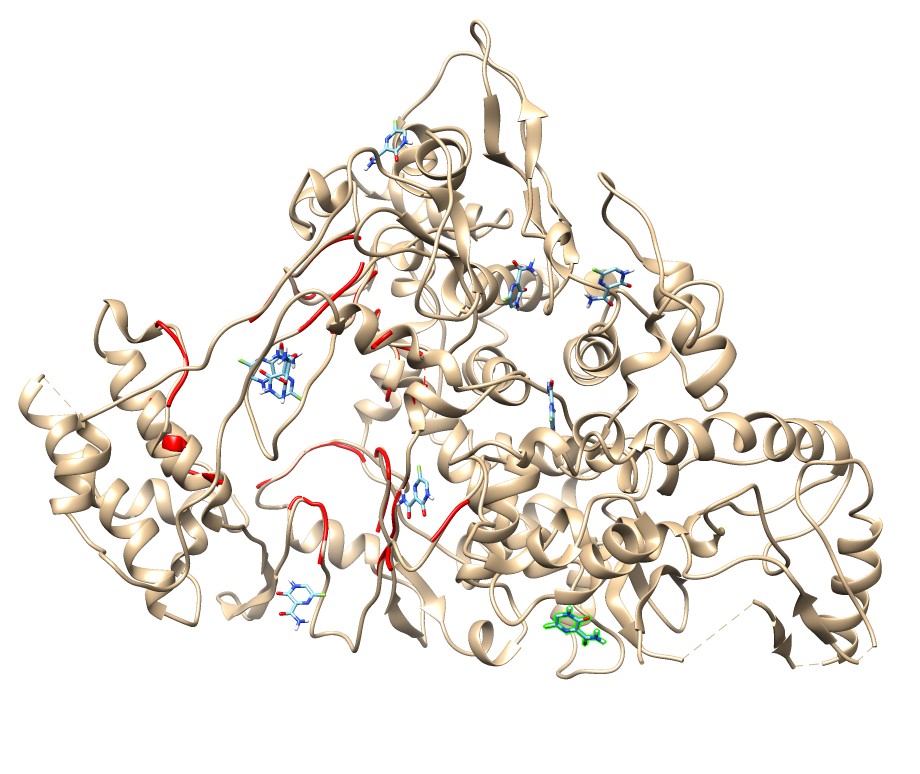

### favi_6vww.jpg

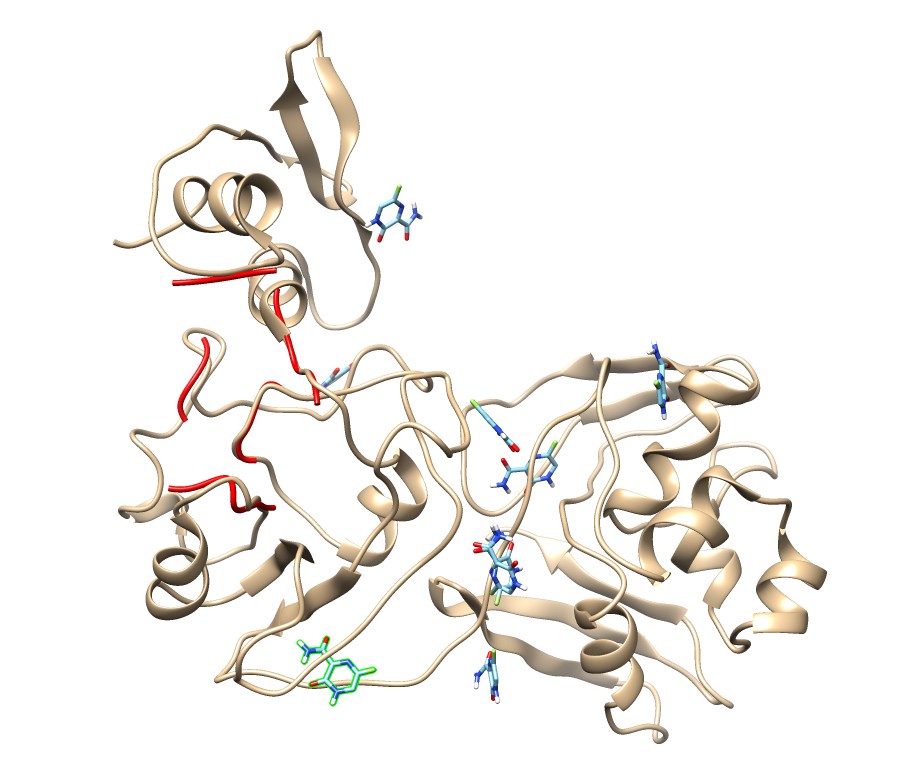

### favi_6vxx.jpg

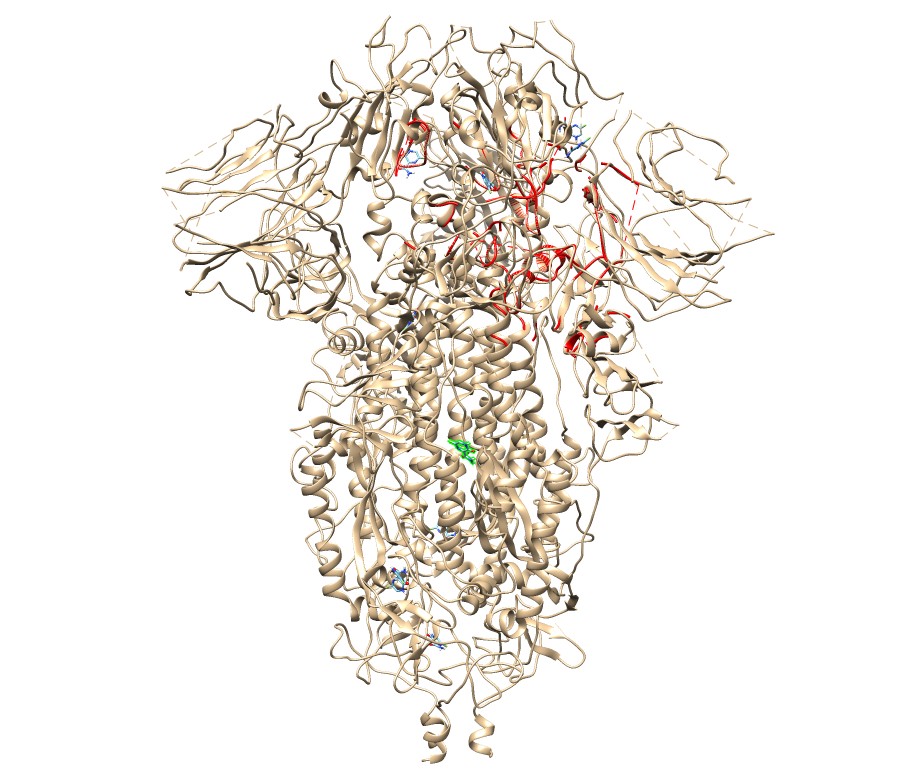

### favi_6vyb.jpg

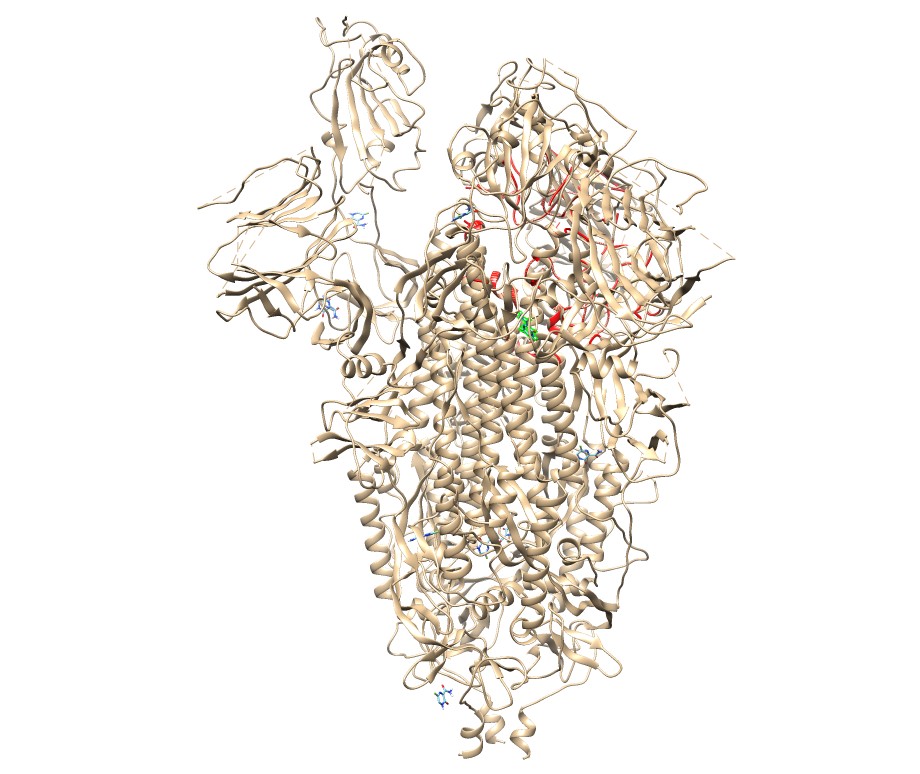

### favi_6vyo.jpg

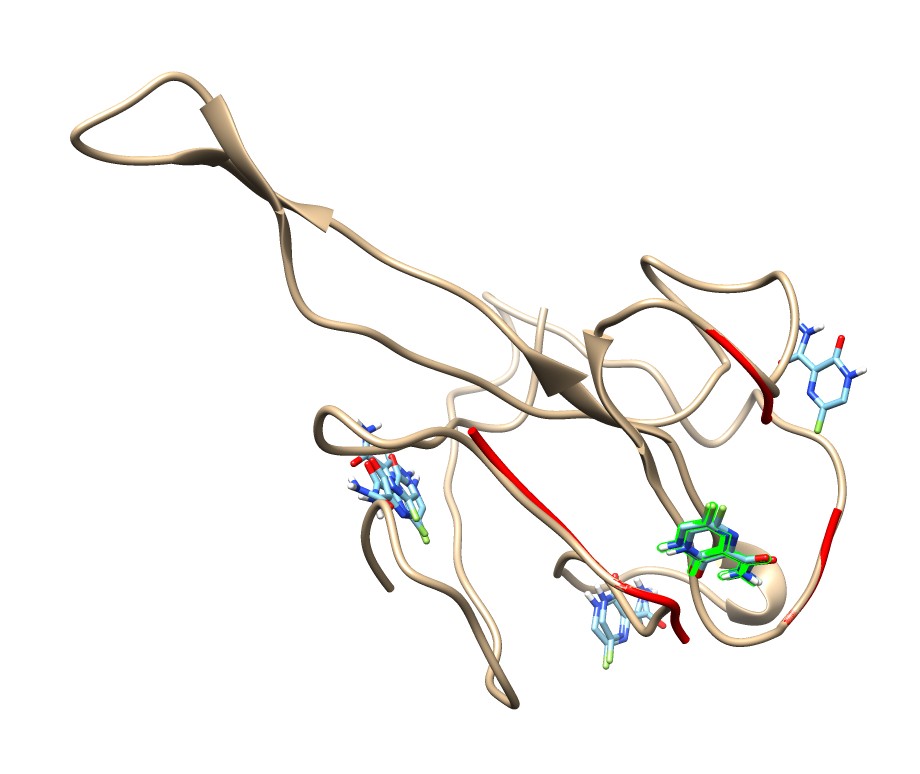

### favi_6y84.jpg

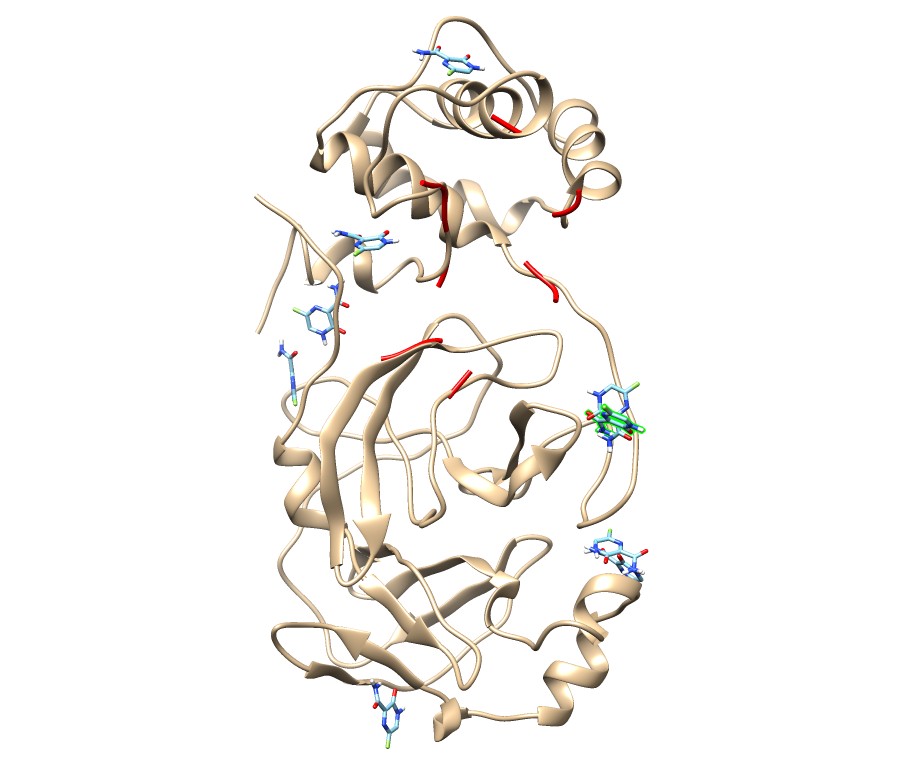

### umif_1r42.jpg

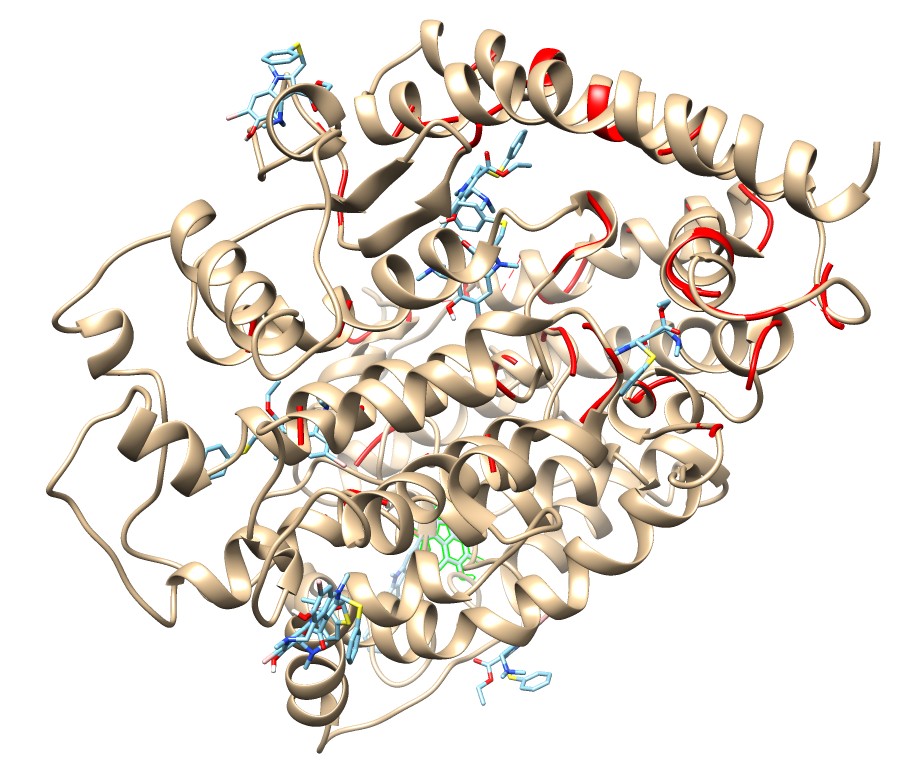

### umif_5x29.jpg

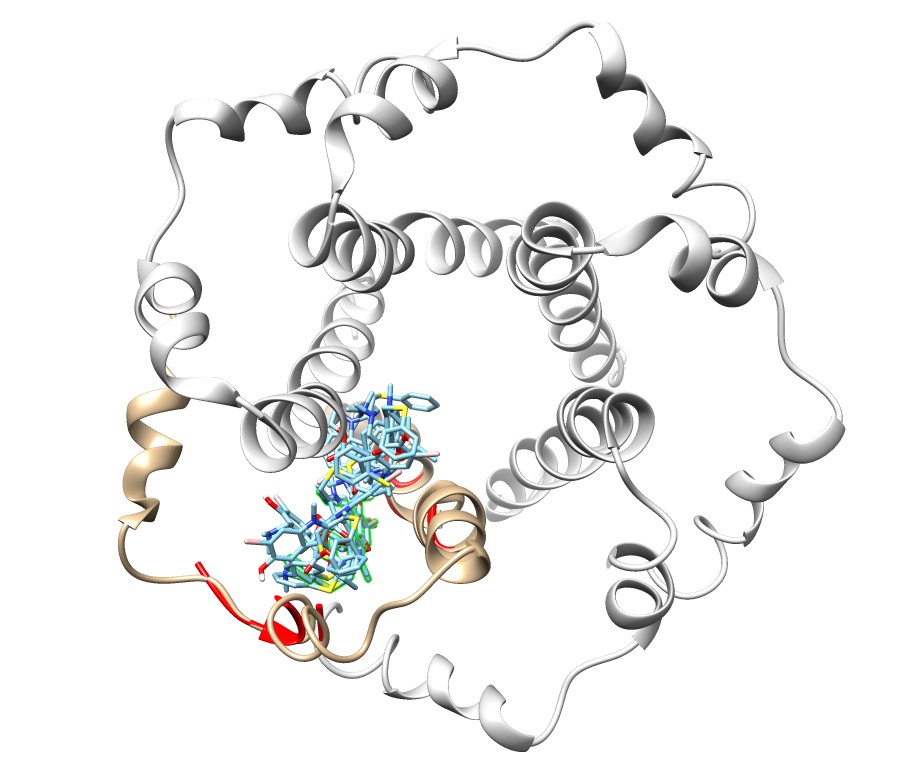

### umif_6lu7.jpg

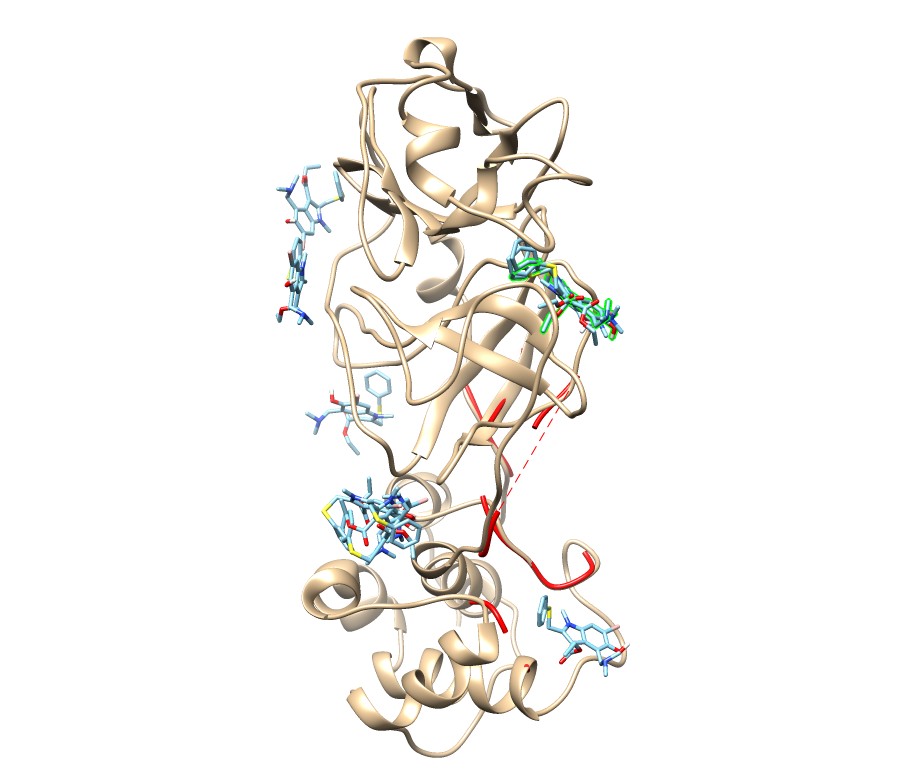

### umif_6lxt.jpg

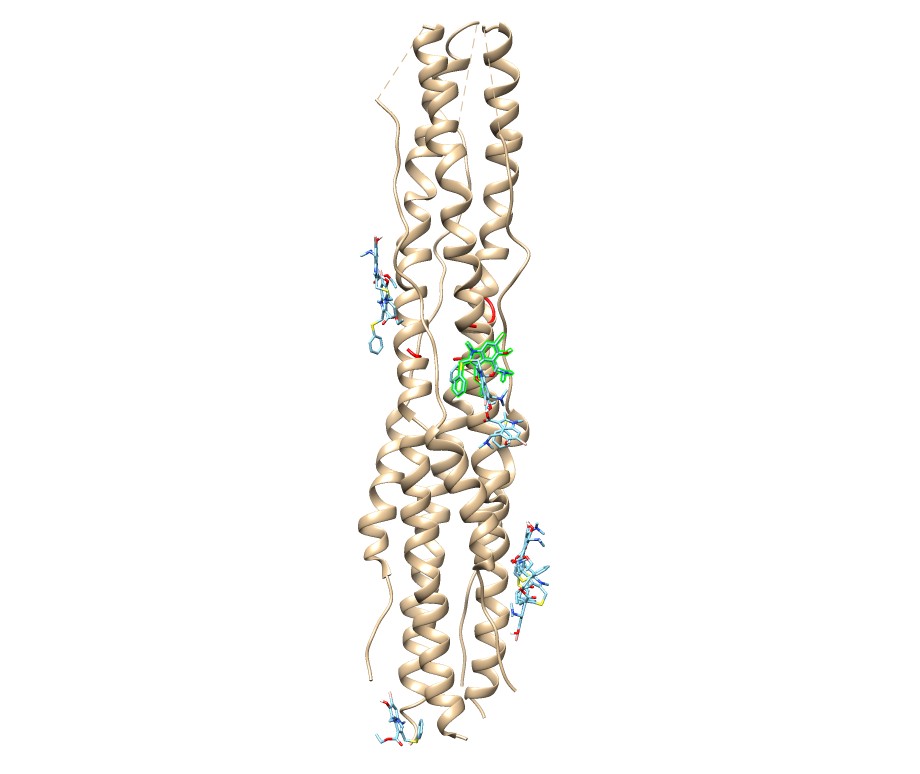

### umif_6m0j.jpg

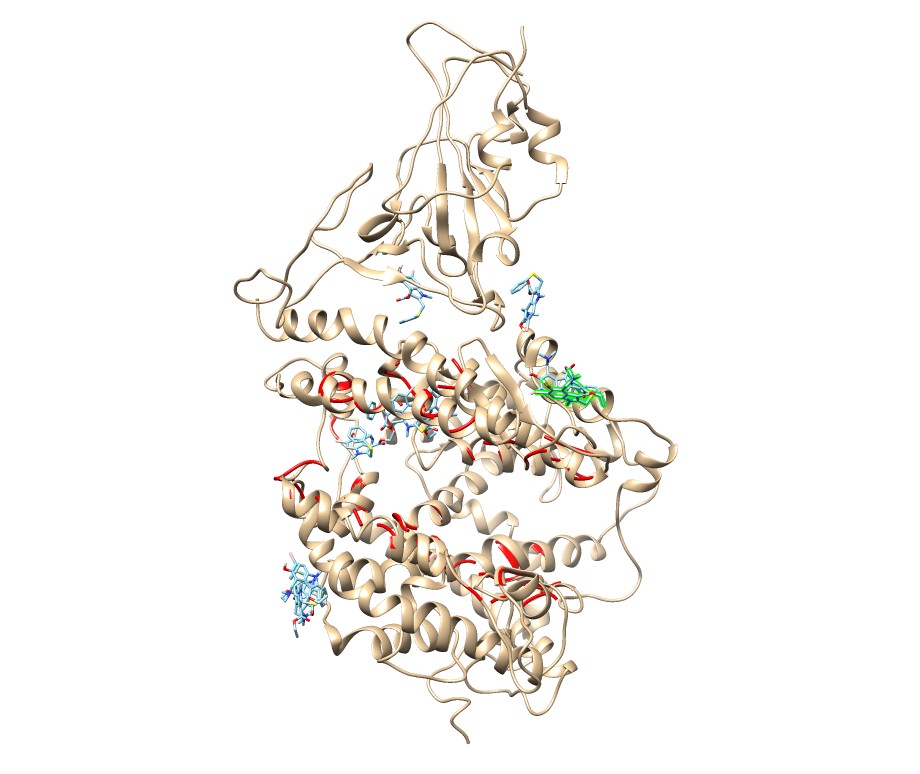

### umif_6m03.jpg

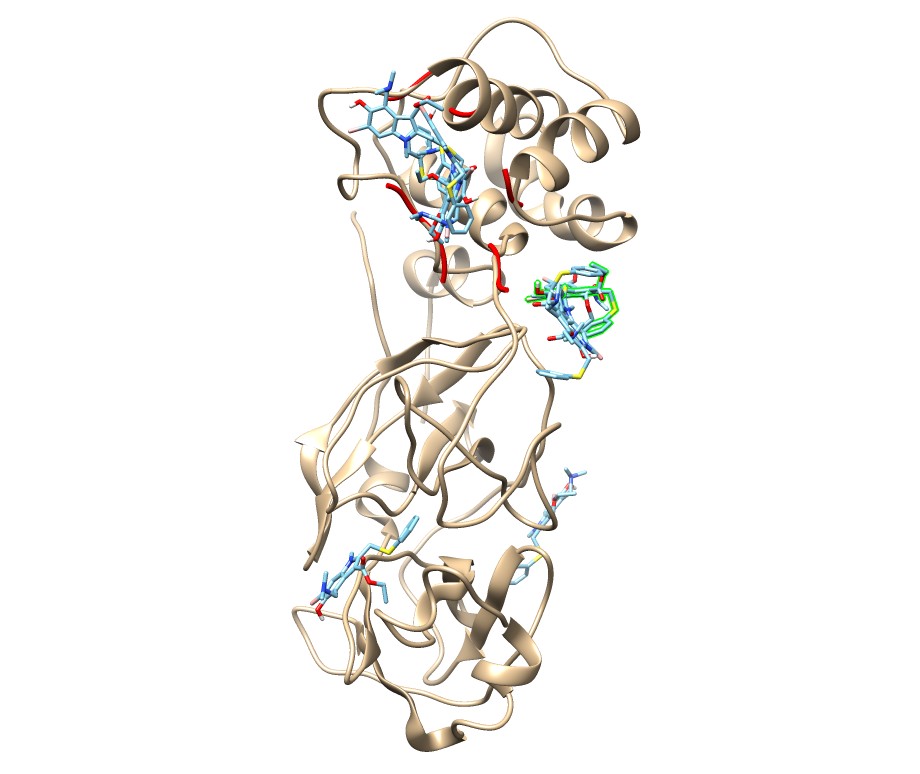

### umif_6m71.jpg

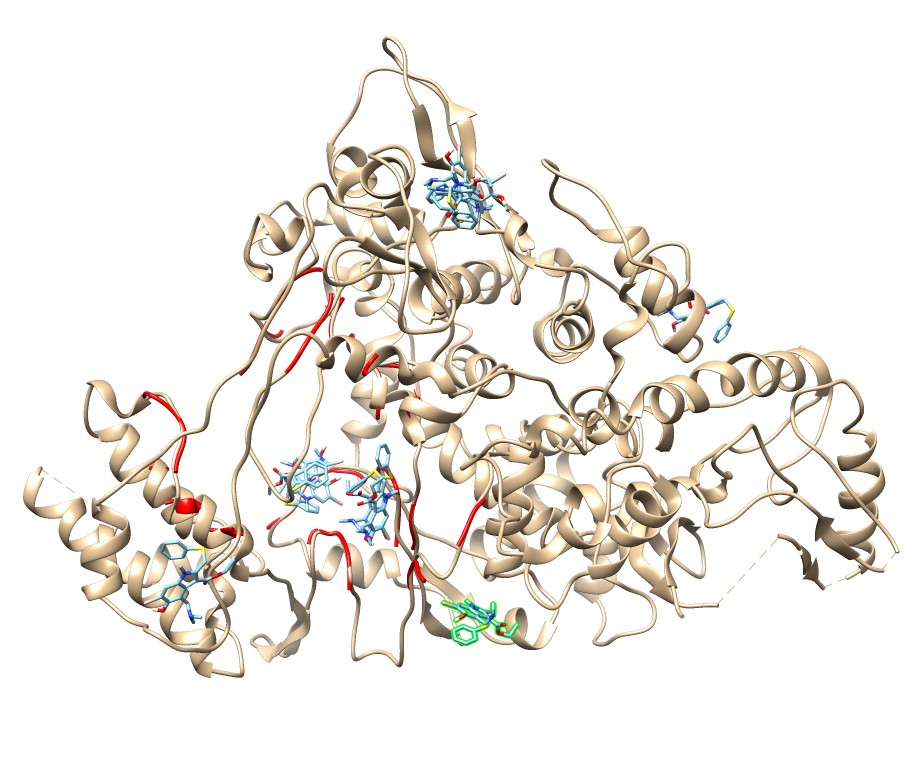

### umif_6vww.jpg

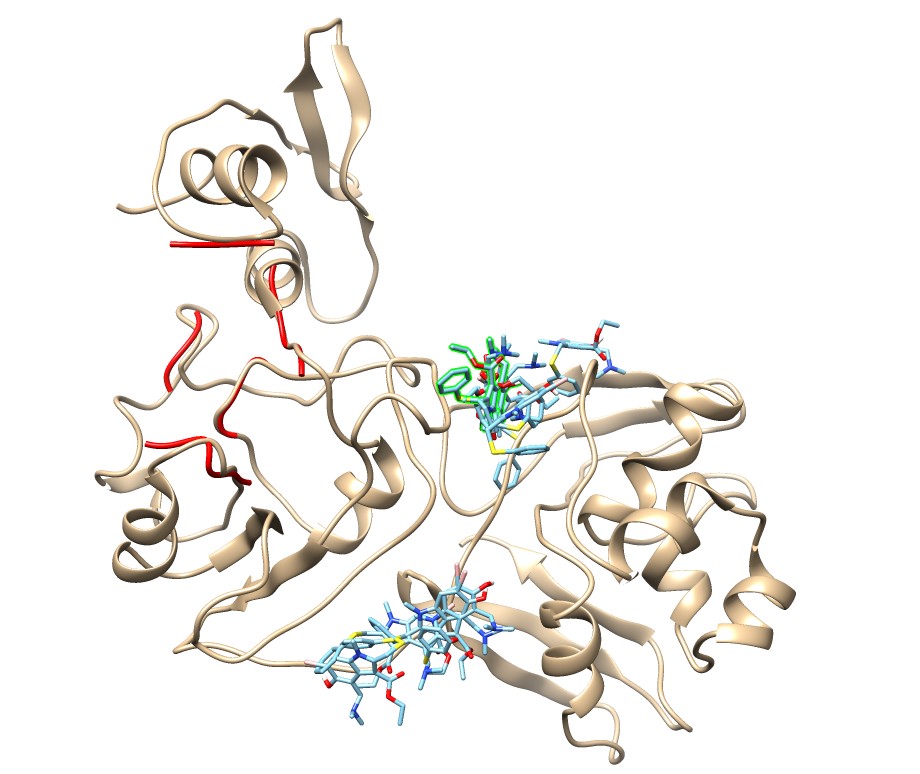

### umif_6vxx.jpg

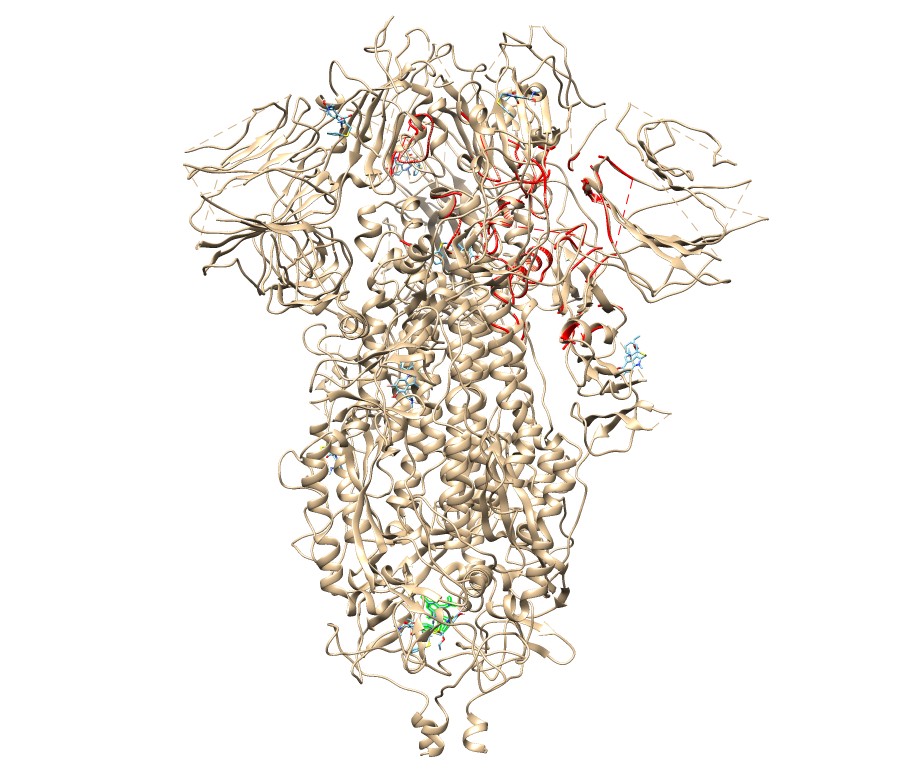

### umif_6vyb.jpg

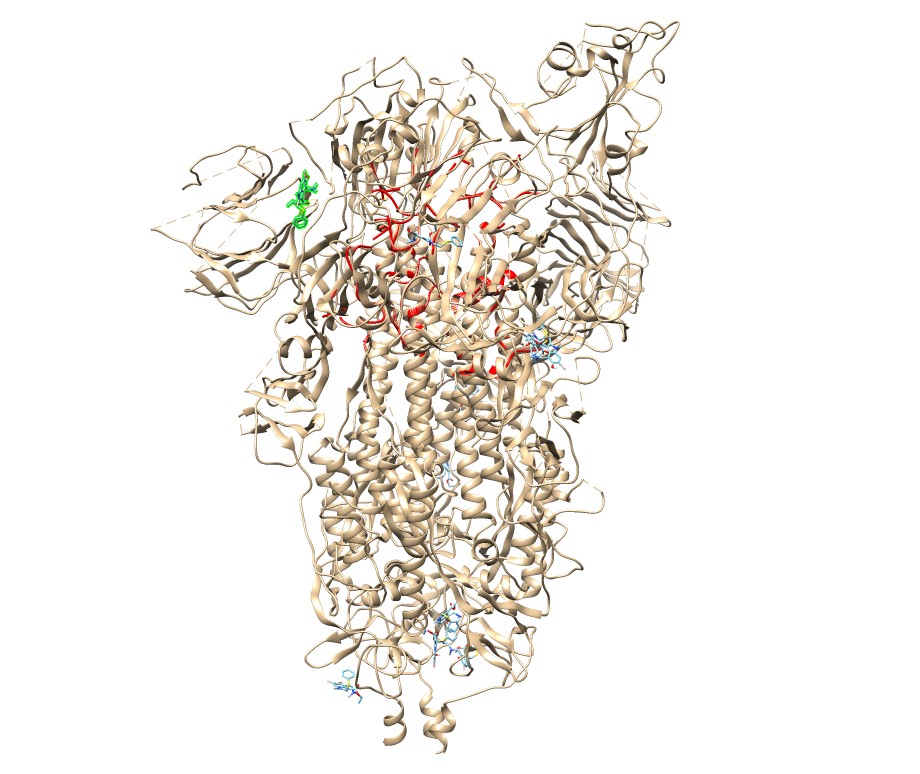

### umif_6vyo.jpg

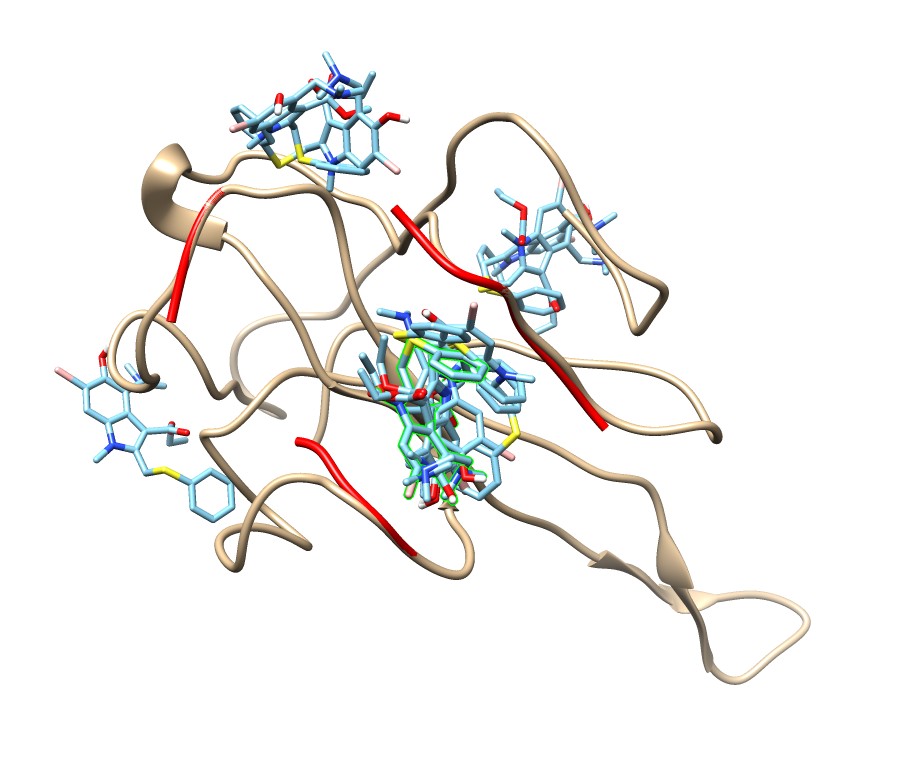

### umif_6y84.jpg

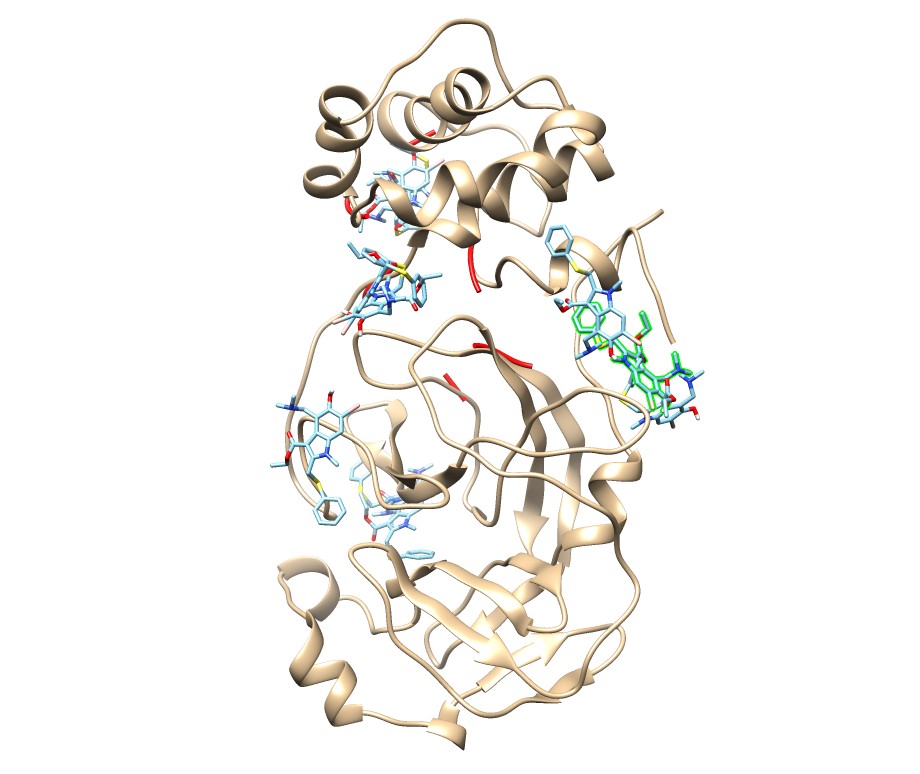

### umiffavi_1r42.jpg

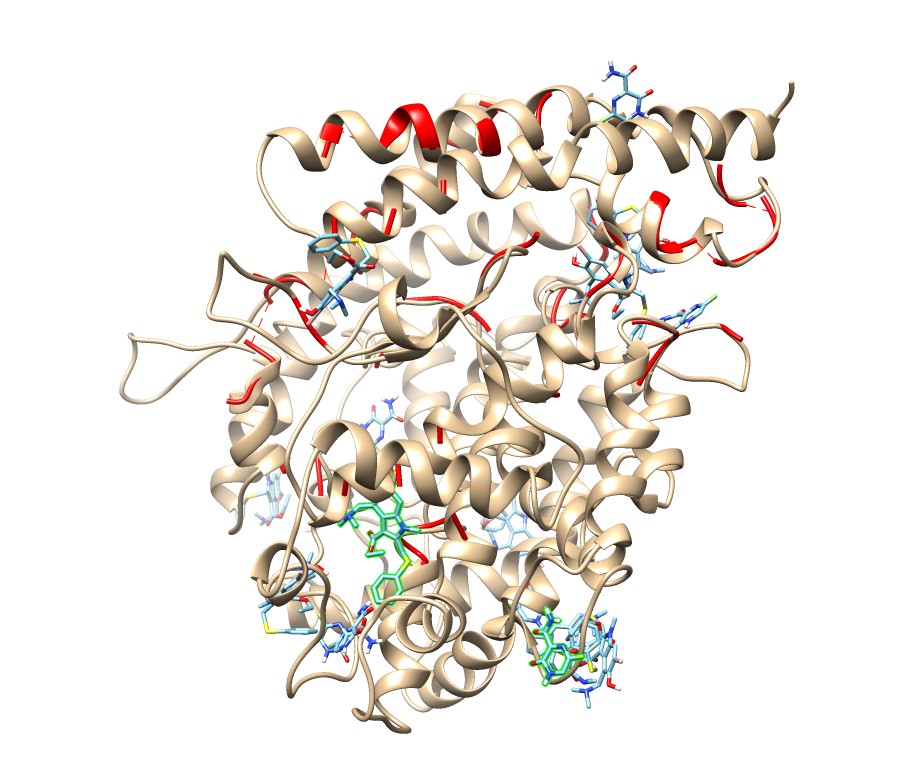

### umiffavi_5x29.jpg

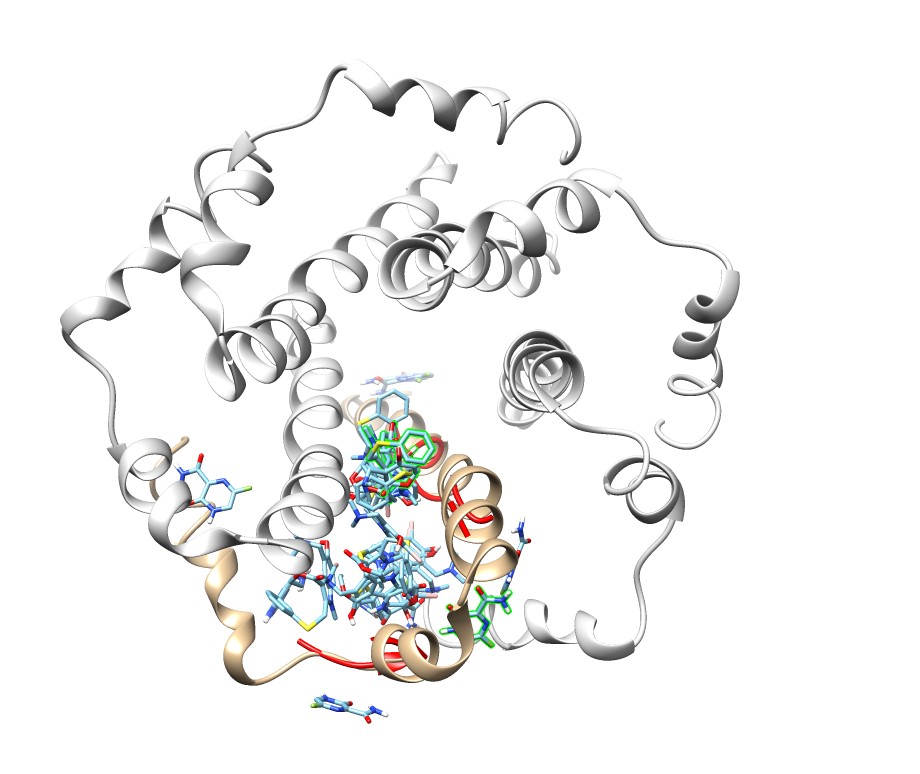

### umiffavi_6lu7.jpg

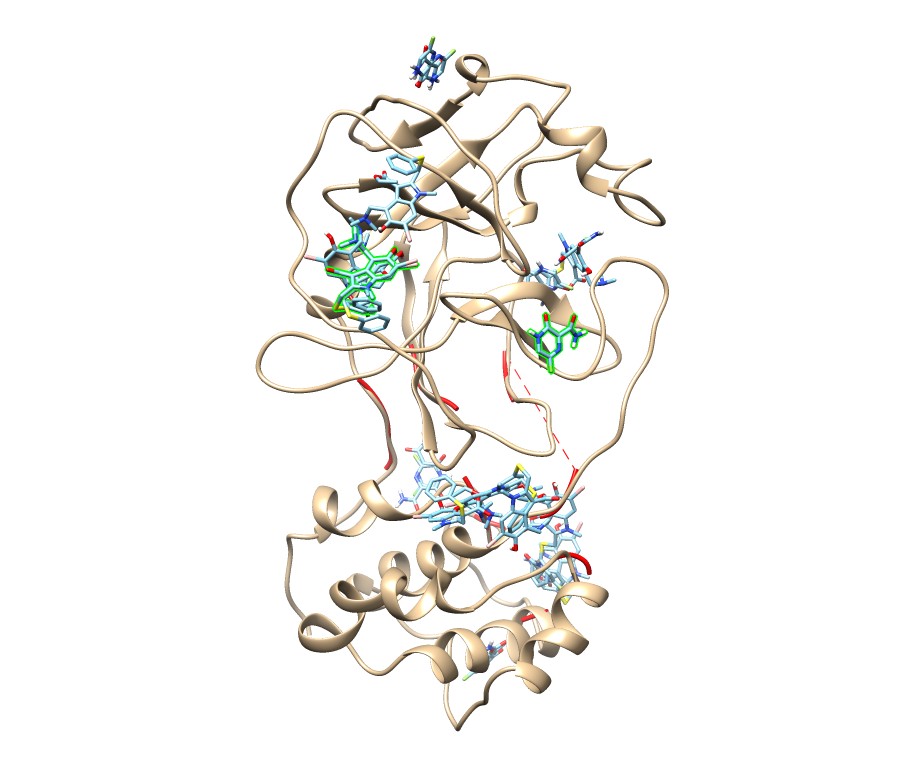

### umiffavi_6lxt.jpg

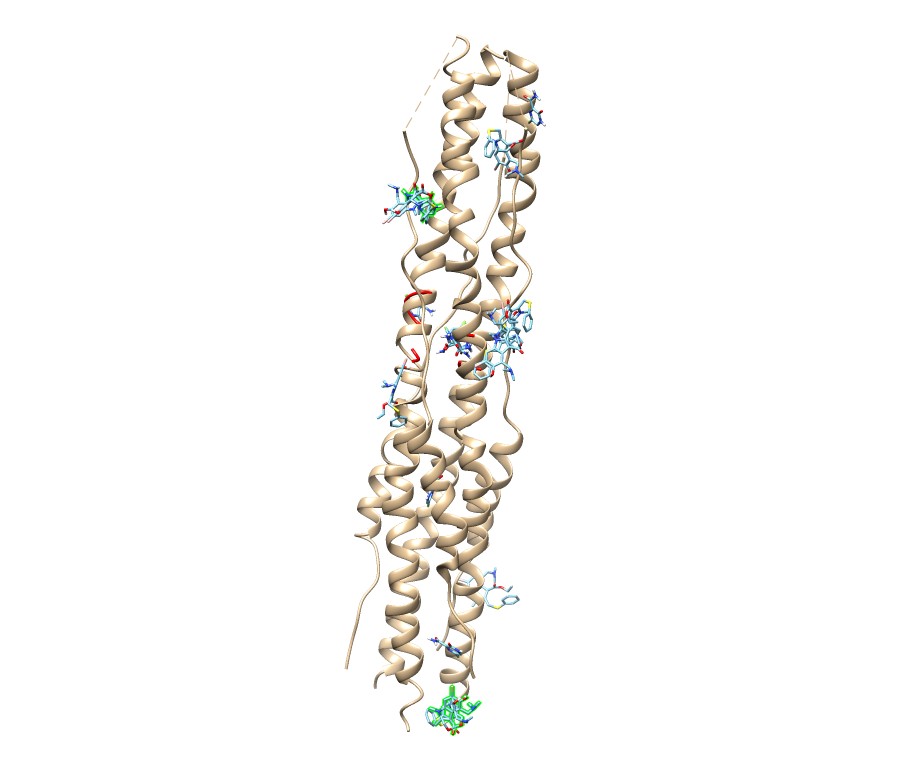

### umiffavi_6m0j.jpg

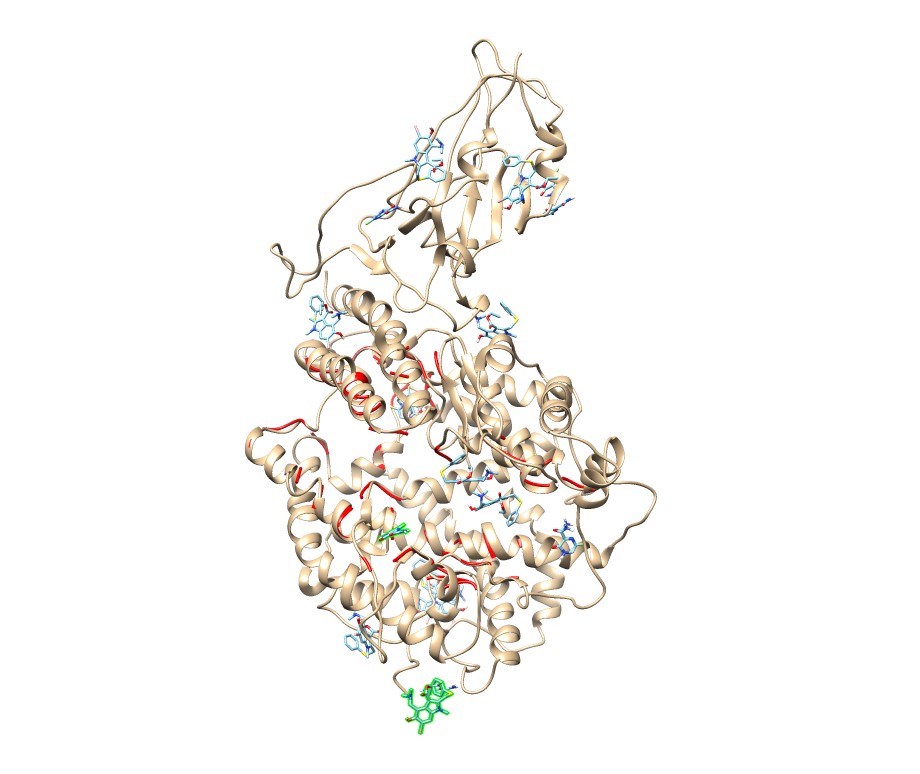

### umiffavi_6m03.jpg

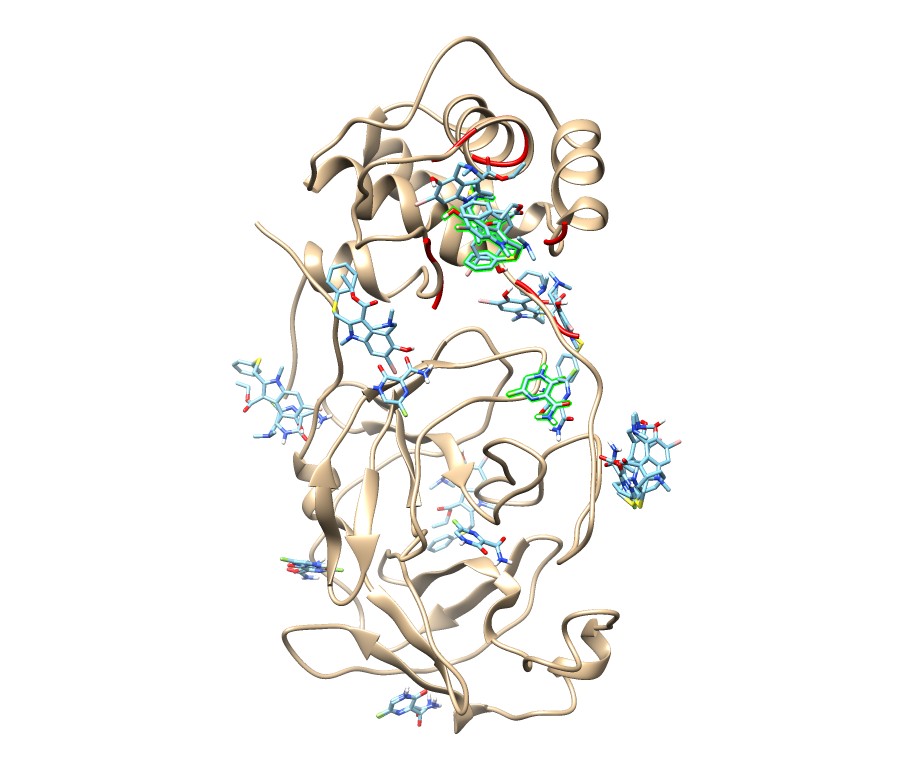

### umiffavi_6m71.jpg

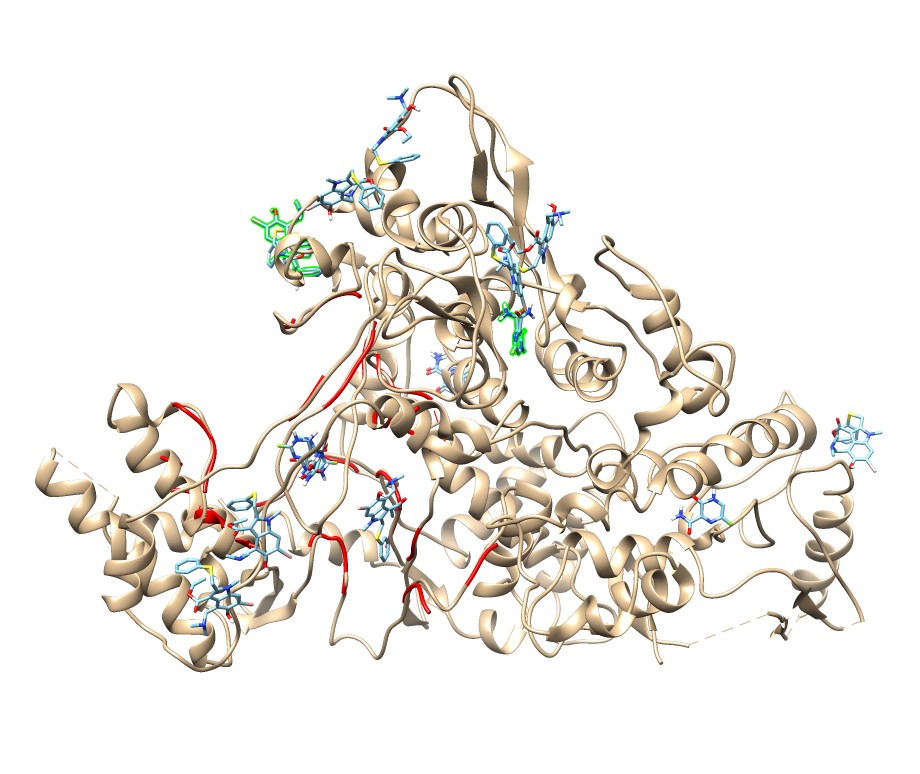

### umiffavi_6vww.jpg

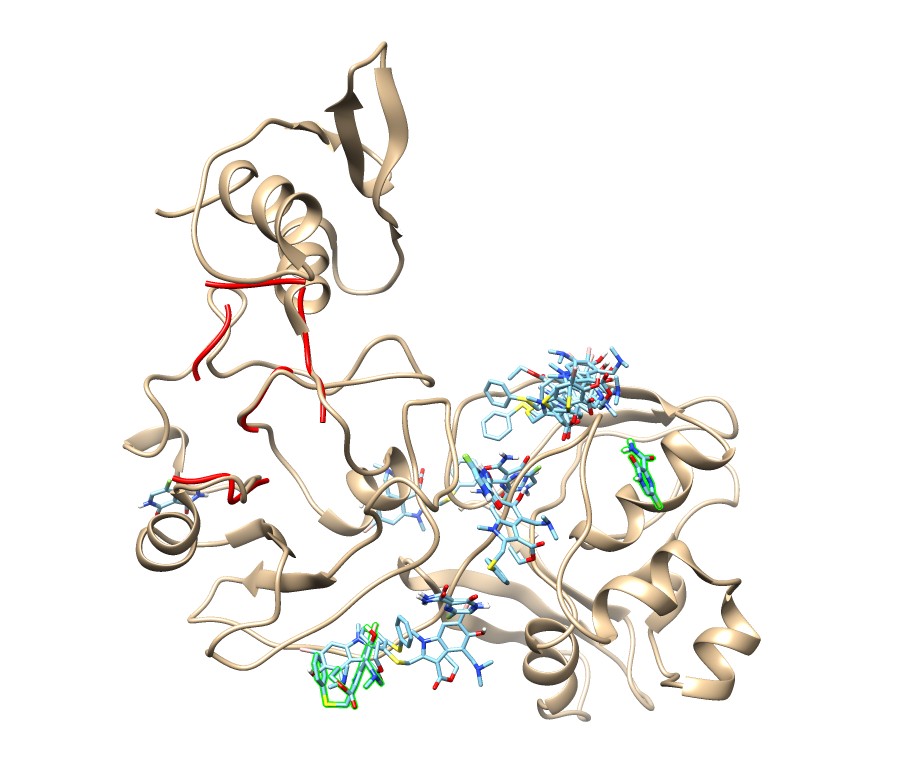

### umiffavi_6vxx.jpg

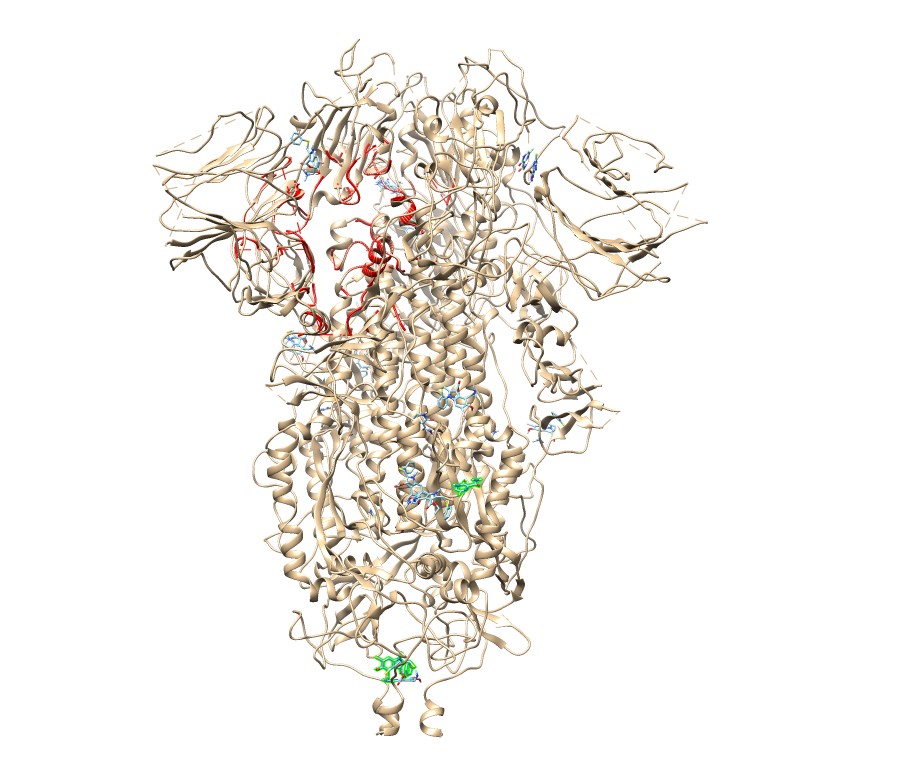

### umiffavi_6vyb.jpg

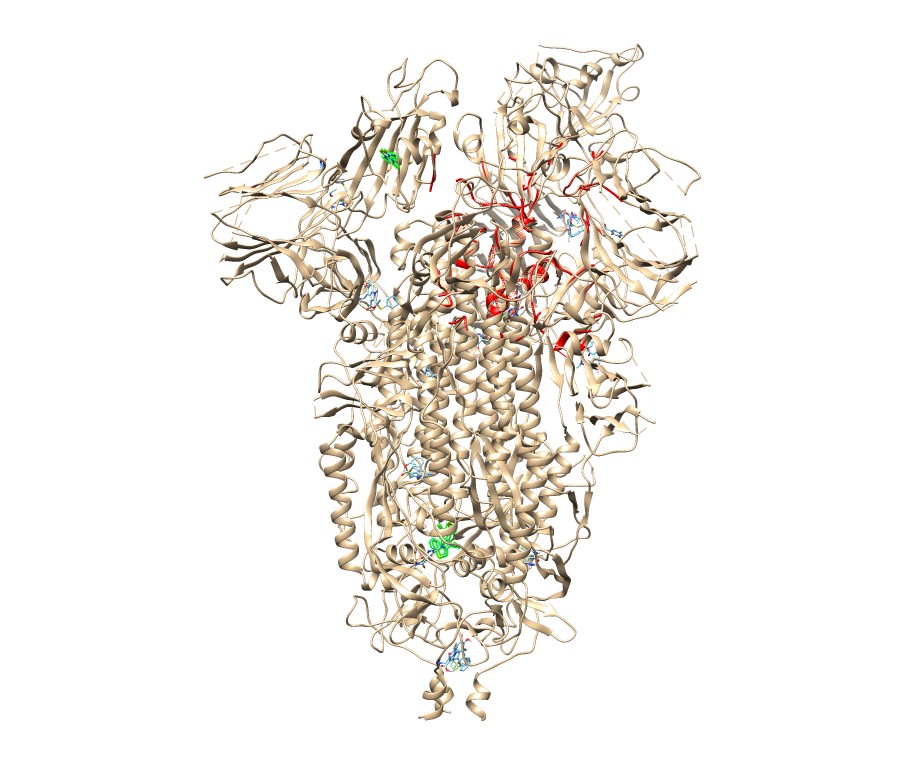

### umiffavi_6vyo.jpg

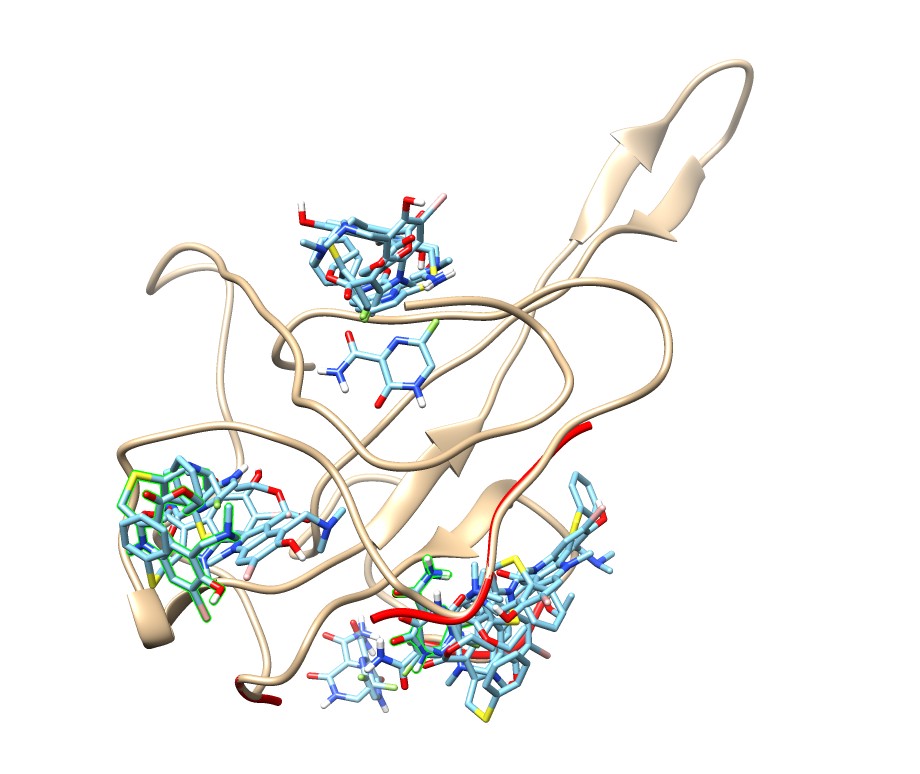

### umiffavi_6y84.jpg

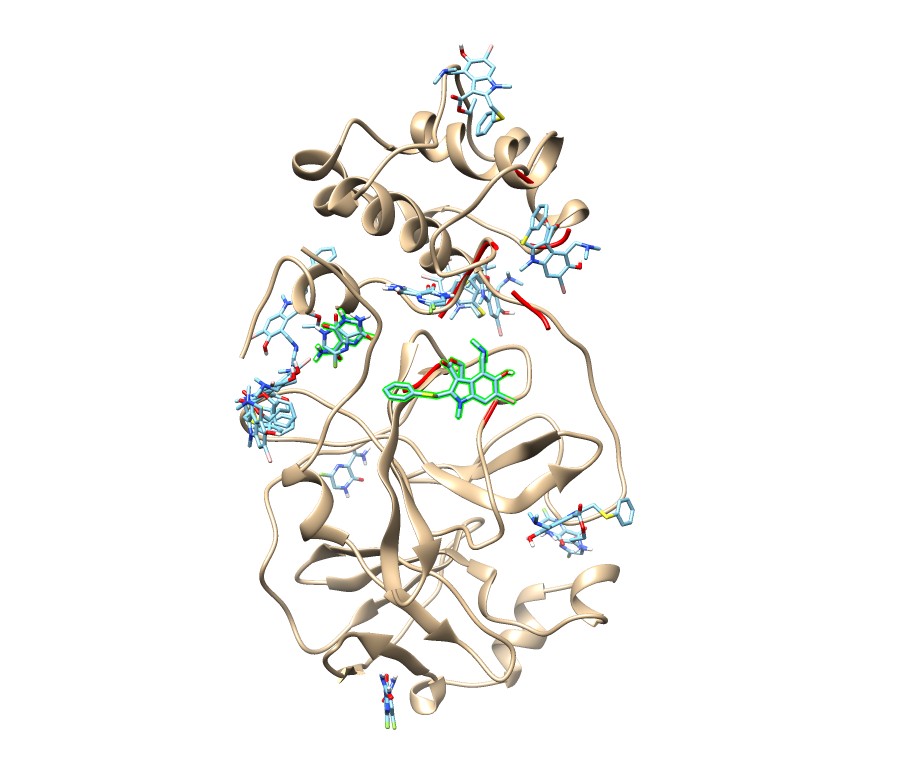
